# Structure-guided design and synthesis of C22- and C32-modified FK520 analogs with enhanced activity against human pathogenic fungi

**DOI:** 10.1101/2024.09.27.615491

**Authors:** Patrick A. Dome, Pyeonghwa Jeong, Gibeom Nam, Hongjun Jang, Angela Rivera, Anna Floyd Averette, Eunchong Park, Tzu-Chieh Liao, Maria Ciofani, Jianli Wu, Jen-Tsan Ashley Chi, Ronald A. Venters, Hyun-Ju Park, William J. Steinbach, Praveen R. Juvvadi, Joseph Heitman, Jiyong Hong

**Affiliations:** Department of Chemistry, Duke University, Durham, NC 27708, United States; School of Pharmacy, Sungkyunkwan University, Suwon 16419, Republic of Korea; Department of Pharmacology and Cancer Biology, Duke University Medical Center, Durham, NC 27710, United States; Department of Molecular Genetics and Microbiology, Duke University Medical Center, Durham, NC 27710, United States; Department of Integrative Immunobiology, Duke University Medical Center, Durham, NC 27710, United States; Duke University NMR Center, Duke University, Durham, NC 27710, United States; Department of Radiology, Duke University Medical Center, NC 27710, United States; Department of Pediatrics, University of Arkansas for Medical Sciences, Little Rock, AR 72202, United States

**Keywords:** calcineurin, FKBP12, FK506, antifungal

## Abstract

Invasive fungal infections are a leading cause of death worldwide. Translating molecular insights into clinical benefits is challenging because fungal pathogens and their hosts share similar eukaryotic physiology. Consequently, current antifungal treatments have limited efficacy, may be poorly fungicidal in the host, can exhibit toxicity, and are increasingly compromised by emerging resistance. We have established that the phosphatase calcineurin (CaN) is required for invasive fungal disease and an attractive target for antifungal drug development. CaN is a druggable target, and there is vast clinical experience with the CaN inhibitors FK506 and cyclosporin A (CsA). However, while FK506 and its natural analog FK520 exhibit antifungal activity, they are also immunosuppressive in the host and thus not fungal-selective. We leverage our pathogenic fungal CaN-FK506-FKBP12 complex X-ray structures and biophysical data to support structure-based ligand design as well as structure–activity relationship analyses of broad-spectrum FK506/FK520 derivatives with potent antifungal activity and reduced immunosuppressive activity. Here we apply molecular docking studies to develop antifungal C22- or C32-modified FK520 derivatives with improved therapeutic index scores. Among them, the C32-modified FK520 derivative JH-FK-44 (**7**) demonstrates a significantly improved therapeutic index compared to JH-FK-08, our lead compound to date. NMR binding studies with C32-derivatives are consistent with our hypothesis that C32 modifications disrupt the hydrogen bonding network in the human complex while introducing favorable electrostatic and cation–π interactions with the fungal FKBP12 R86 residue. These findings further reinforce calcineurin inhibition as a promising strategy for antifungal therapy.

**Significance:** Invasive fungal infections cause significant mortality worldwide, and current antifungal treatments are often ineffective, toxic, or face growing resistance. This research identifies calcineurin (CaN), a critical protein for fungal survival, as a potential target for developing new antifungal drugs. Although existing CaN inhibitors such as FK506 (tacrolimus) and FK520 (ascomycin) possess antifungal properties, their immunosuppressive effects limit their clinical utility. By studying the structure of human and fungal FKBP12-FK506 or FK520 complexes with CaN, we have designed and synthesized modified FK520 derivatives with strong antifungal activity and reduced immunosuppressive effects. These new derivatives are expected to have significantly improved therapeutic profiles, offering hope for more effective and safer antifungal treatments.

## Introduction

Fungal infections impact over one billion people worldwide, leading to more than 2.5 million deaths annually.^1^ Invasive fungal infections, particularly those caused by *Aspergillus, Candida*, and *Cryptococcus*, significantly contribute to morbidity and mortality globally, especially in immunocompromised patients. *Candida* species rank as the fourth most common isolate in all bloodstream infections,^2^ and *Cryptococcus neoformans* is responsible for cryptococcal meningoencephalitis, accounting for approximately 15% of HIV/AIDS-related deaths.^3^ Over the past decade, both the incidence and mortality rates of invasive aspergillosis have tripled.^4,5^ In 2019, fungal diseases were estimated to cost over $11.5 billion in the US,^6^ with antifungals constituting the largest proportion of anti-infective expenditures in many major medical centers.^7^

Given the substantial global health burden posed by deadly fungal infections, the diversity among antifungal agents remains limited, with only four major classes currently in clinical use.^8^ Current therapies for invasive candidiasis have an approximate success rate of 50–70%, primarily in the healthiest patients.^9^ Recommended treatments for invasive aspergillosis have a success rate of 30–50%.^10,11^ Additionally, these clinically used antifungal agents are constrained by toxicity, adverse drug interactions, or limited scope.^8,12^ At the same time, widespread antifungal resistance is steadily eroding the effectiveness of existing treatment options.

Despite the urgent need, ibrexafungerp, the first antifungal in a novel class to be developed in over 20 years,^13^ is currently only approved for the treatment of vulvovaginal candidiasis. Like other existing antifungal treatments, it still targets the fungal cell wall and the same enzyme as the echinocandin class of antifungals.^14^ Consequently, the growing immunocompromised patient population has outpaced antifungal development, and antifungal resistance continues to hamper effective treatment. Therefore, innovative approaches to exploring broad, new antifungal targets with novel mechanisms of action are urgently needed.

The phosphatase calcineurin (CaN) has been proposed as a promising target for antifungal development. CaN is a highly conserved serine/threonine-specific Ca^2+^-calmodulin-dependent phosphatase important in fungal pathogenesis,^15^ rendering it an ideal target for antifungal development. It is a heterodimer composed of a catalytic (CnA) and a regulatory (CnB) subunit, and it is the target of the immunophilin-immunosuppressant complexes (FKBP12-FK506 and cyclophilin A-CsA), which inhibit its activity.^16–18^ Our comprehensive iterative genetic and pharmacologic approaches have established the importance of CaN for growth, survival in serum, and virulence in a range of fungal pathogens, including *A. fumigatus*, *C. albicans*, *C. neoformans,* and *M. circinelloides*.^19^ In fungal systems, CaN binds to Ca^2+^ and calmodulin, and the combined calmodulin-CaN complex then dephosphorylates Crz1, leading to its nuclear translocation and regulation of gene expression.^20^ We have demonstrated that CaN inhibitors are effective against azole- and echinocandin-resistant *A. fumigatus* strains.^21^ Additionally, when used in combination with echinocandins, they exhibit fungicidal properties.^22,23^ CaN inhibitors combined with fluconazole are fungicidal against *C. albicans,* and effective against fluconazole-resistant strains of *C. krusei* and *C. glabrata.*^24^ Synergism of CaN inhibitors with echinocandins and azoles was also observed against clinical isolates of *C. neoformans* and *R. oryzae*.^25,26^ These findings underscore CaN as a prime target for antifungal development. However, CaN is also present in humans. Like its fungal counterpart, human CaN binds to calmodulin, which dephosphorylates NFAT, allowing its nuclear translocation to upregulate the expression of IL-2, a cytokine essential for the T-cell-mediated immune response.^16,27,28^ Inhibition of human CaN suppresses the immune system.^29–31^ Consequently, developing fungal-specific CaN inhibitors with reduced host immunosuppressive activity has been challenging.^32–35^

FK506 (**1**, **Figure 1**), also known as tacrolimus, is a natural product inhibitor of CaN. FK506 binds to the *cis-trans* peptidyl-prolyl isomerase FKBP12, and the FKBP12-FK506 complex then binds to CaN.^36^ FK506 has robust *in vitro* antifungal activity against critical pathogens such as *A. fumigatus, C. albicans,* and *C. neoformans.*^37^ However, FK506 is also a potent immunosuppressant and is FDA-approved for preventing post-transplant organ rejection.^29^ Due to its long-standing application for organ transplants, numerous chemically and biologically derived analogs of FK506 have been developed, many of which aim to modulate the immunosuppressive activity of FK506.^38–41^

**Figure 1.**
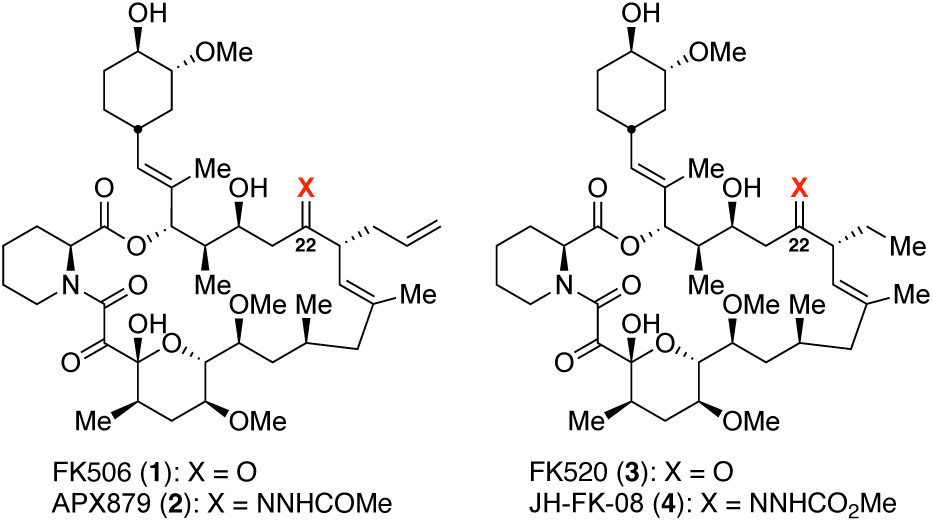
The structures of FK506, APX879, FK520, and JH-FK-08.

In 2019, our group examined the crystal structures of the human and *A. fumigatus* fungal FKBP12-FK506-CaN complexes and discovered a critical difference:^42^ the fungal FKBP12 residue F88 versus the human FKBP12 residue H88 located on the binding surface of CaN with FK506. Modification of FK506 near its interface with F88/H88 was identified as a potential strategy for developing a selective inhibitor targeting fungal CaN. We applied this strategy to design APX879 (**2**), a C22-acetylhydrazone derivative of FK506. We demonstrated that APX879 possesses reduced immunosuppressive activity (IL-2 IC50: APX879=13.48 nM vs. FK506=0.19 nM) while also maintaining antifungal activity (MIC against *C*. *neoformans*: APX879=1 μg/mL vs. FK506=0.05 μg/mL).^42^ Building on this promising result, we recently reported a series of C22- modified derivatives of FK520 (**3**), a natural analog of FK506 in which the C21 allyl group is replaced with ethyl group.^43,44^ This effort yielded JH-FK-08 (**4**) which has reduced immunosuppressive activity and is effective in an *in vivo* murine *C. neoformans* infection model. JH-FK-08 showed strong activity against *C. neoformans* with MIC of 0.8 μg/mL while being markedly less immunosuppressive than FK506 with IC50 of 42.6 nM, an approximately 470-fold reduction from FK506. Additionally, JH-FK-08 acts synergistically with fluconazole in the *in vivo*

### C. neoformans infection model

The development of C22-modified FK520 analogs with an improved therapeutic index (TI) score, which represents the ratio of the immunosuppressive to the antifungal fold change (*vide infra*), offers an excellent starting point for further development. Here, we report the application of *in silico* docking in the design of a new generation of C22-modified FK520 derivatives. Our *in silico* docking study also identified an additional site for modification, leading to the development of new C32-modified FK520 derivatives with *in vitro* activity against both *C. neoformans* and *C. albicans.* Our work strongly advances the feasibility of developing CaN inhibitors as a novel class of antifungals.

## Results

### Docking studies identify the potential role of water molecules in the binding of C22-modified FK506/FK520 derivatives to a FKBP12-CaN complex

Our previous chemical, structural, biophysical, and biological studies on the C22 modification of FK506 and FK520 identified C22 as a strategic site for developing non-immunosuppressive derivatives with antifungal properties.^42–44^ After completing the initial biological testing of C22 derivatives, we shifted our focus to refining the approach to identify potential non-immunosuppressive, antifungal C22 derivatives. Given the potential of *in silico* docking for antifungal drug development, particularly in modifying established antifungal classes,^45–48^ it was employed to investigate the binding of C22-modified FK520 derivatives to the FKBP12-CaN complex. The goal was to improve the selectivity (antifungal vs. immunosuppressive) of FK506/FK520 derivatives.

Docking models for C22-modified FK520 derivatives bound to FKBP12-CaN complexes were constructed for both the human and *C. neoformans* complexes based on available crystal structures (PDB: 7U0T and 6TZ8, respectively). Analysis of docking models for C22 derivatives in the human and fungal binding pockets suggested that the amino acid residue 88 of FKBP12 in both crystal structures might be pivotal in distinguishing the two complexes, in accord with previous analyses of crystal structures.^42^ As illustrated in **Figure 2A** and **2B**, the C22-carbonyl group of FK506/FK520 engages in hydrogen bonding within an extensive network that connects FKBP12, FK520/FK506, and CaN through co-crystallized water molecules, interacting differently with human H88 and fungal F88 residues. In the human enzyme, modification at C22 is expected to disrupt the native hydrogen bonding with H88, potentially weakening the binding of human CaN to the FKBP12-FK506/FK520 complex while having minimal impact on the fungal system. This makes C22 a promising modification site for introducing fungal selectivity. Our hypothesis for C22 modifications to achieve fungal selectivity was supported by the improvement in TI score of several C22-modified FK520 derivatives reported by Rivera *et al*.^43^ Comparison of APX879 (**2**, **Figure 1**), a C22-acetylhydrazone derivative, bound to human FKBP12 (PDB: 6VCU) with both human and fungal FKBP12-FK506/FK520-CaN complexes also revealed that the carbonyl group of APX879’s acetylhydrazone moiety may replace the co-crystallized water and interact with F88/H88 of fungal and human complexes, respectively (**Figure 2C** and **2D**).

**Figure 2.**
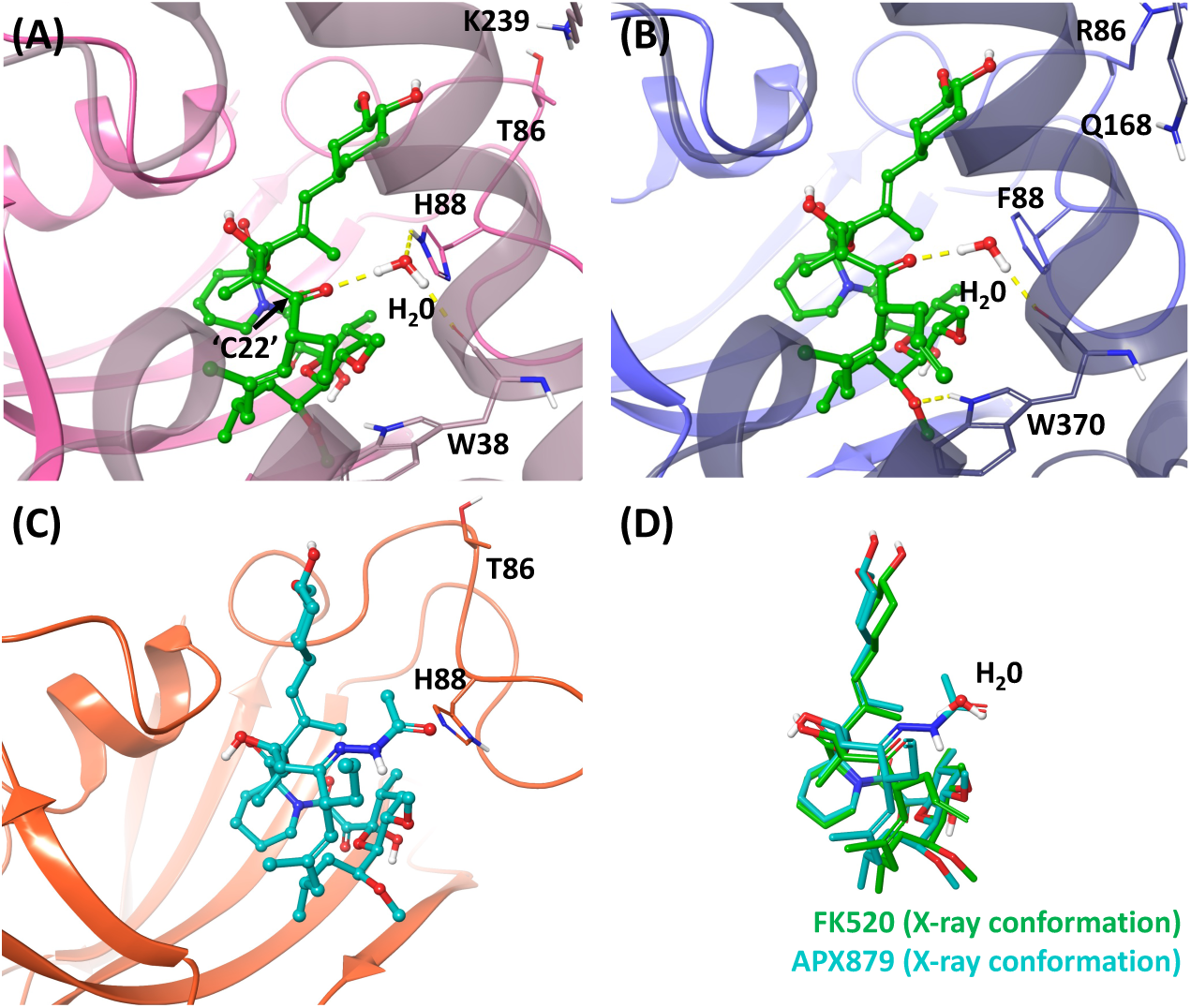
(A) Crystal structure of human FKBP12-FK520-CaN complex (PDB 7U0T). FKBP12 and CaN are colored in pink and dark pink, respectively. FK520 is colored in green. (B) Crystal structure of *C. neoformans* FKBP12-FK506-CaN complex (PDB 6TZ8). FKBP12 (blue), FK506 (green), CaN (dark blue), and a co-crystallized water molecule are shown. Hydrogen bonds are represented by yellow dashed lines. (C) Crystal structure of human FKBP12-APX879 complex (PDB 6VCU). FKBP12 (orange) and APX879 (teal) are shown. (D) Superimposition of FK506 (6TZ8), co-crystallized water, and APX879 (6VCU).

Analysis of docking models of C22 derivatives in the human and fungal binding pockets suggested that the human FKBP12-FK520-CaN pocket is generally more tolerant of bulky and hydrophobic substitution than the fungal system. Conversely, the C22 binding pocket of *C. neoformans* shows a preference for smaller and less hydrophobic substituents at the C22 position compared to its human counterpart. Our *in silico* docking model also suggested that C22 hydrazones or oximes, without the hydrazide carbonyl group, may exhibit reduced immunosuppressive activity due to a reduced interaction with H88 of human FKBP12. In summary, the docking model proposed two potential avenues for optimization: the synthesis of C22-hydrazide derivatives incorporating small, hydrophilic substituents, and the inclusion of hydrazones and oximes over hydrazides at C22.

### Design and synthesis of C22-modified FK520 derivatives aided by molecular docking

Based on our *in silico* docking study, we selected seven commercially available hydrazides and subjected them to condensation reactions with FK520 via the acidic conditions previously developed for FK520 modification to yield JH-FK-29, 30–34, and 36 in a single step (**Supplementary Table S1**).^43^ To further explore the effect of steric repulsion by the C22 hydrazide carbonyl group, we additionally synthesized seven oximes and hydrazones (JH-FK-35, 37, 38, 40, 43, 45, and 46) according to aforementioned condensation reactions or via an alternative sodium acetate-facilitated procedure reported by Yadav *et al*.^49^

### C22-modified FK520 derivatives maintain antifungal activity

The antifungal activity of the new C22-modified FK520 derivatives (JH-FK-29–38, 40, 43, 45, and 46) was evaluated *in vitro* against *C. neoformans* and *A. fumigatus* (**Table 1**). Of the fourteen C22-derivatives we tested, eleven compounds showed more potent antifungal activity than 2.5 µg/mL against *C. neoformans*, and seven of these compounds had a lower minimum inhibitory concentration (MIC80) than the lead compound JH-FK-08 (0.8 µg/mL). Although all fourteen compounds showed less potent antifungal activity than FK506, the C22 derivatization guided by *in silico* docking overall yielded potent antifungal compounds. Immunosuppressive activity was assessed by measuring the reduction of interleukin 2 (IL-2) expression from CD4^+^ T cells cultured in the presence of C22- derivatives (**Table 1**; **Supplementary Figure S1 and S2**). Since CaN is essential for IL-2 expression, a reduction in IL-2 expression serves as an indirect measure of mammalian CaN inhibition. All fourteen derivatives displayed reduced immunosuppressive activity compared to FK506, with five of them being less immunosuppressive than JH-FK-08 (42.6 nM).

**Table 1.** Antifungal and immunosuppressive activity and therapeutic index (TI and TI*) scores of C22-modified FK520 derivatives.

| Compd. | MIC<sub>80</sub><sup>a</sup><br>(Cn<sup>b</sup>) | MEC<sup>a</sup><br>(Af<sup>c</sup>) | IL-2<br>IC<sub>50</sub><sup>d</sup> | TI<sup>e</sup><br>(Cn<sup>b</sup>) | TI*<sup>f</sup><br>(Cn<sup>b</sup>, ×10<sup>-3</sup>) | TI<sup>e</sup><br>(Af<sup>c</sup>) | TI*<sup>f</sup><br>(Af<sup>c</sup>, ×10<sup>-3</sup>) |
| --- | --- | --- | --- | --- | --- | --- | --- |
| FK520<sup>g</sup> | 0.05 | 0.16 | 0.30 | 3.3 | 4.8 | 0.8 | 1.5 |
| JH-FK-08<sup>g</sup> | 0.80 | 1.3 | 43 | 30 | 46 | 15 | 28 |
| FK506 | 0.04–0.06 | 0.039 | 0.04–0.08 | 1.0 | 1.5 | 1.0 | 1.8 |
| JH-FK-29 | 0.60 | 1.3 | 29 | 24 | 42 | 11 | 20 |
| JH-FK-30 | 0.60 | 2.5 | 5.7 | 4.8 | 8.3 | 1.1 | 2.0 |
| JH-FK-31 | 0.20 | 0.31 | 2.1 | 5.3 | 9.2 | 3.3 | 5.9 |
| JH-FK-32 | 1.3 | 5.0 | 48 | 19 | 33 | 4.7 | 8.2 |
| JH-FK-33 | 1.3 | 2.5 | 55 | 22 | 38 | 11 | 19 |
| JH-FK-34 | 1.3 | 2.5 | 34 | 14 | 24 | 6.6 | 12 |
| JH-FK-35 | 0.30 | 0.63 | 4.3 | 7.2 | 12 | 3.3 | 5.5 |
| JH-FK-36 | 0.30 | 0.63 | 0.49 | 0.8 | 1.5 | 0.4 | 0.74 |
| JH-FK-37 | 5.0 | NI<sup>h</sup> | 79 | 7.9 | 13 | ND<sup>i</sup> | ND<sup>i</sup> |
| JH-FK-38 | 2.5 | NI<sup>h</sup> | 214 | 43 | 70 | ND<sup>i</sup> | ND<sup>i</sup> |
| JH-FK-40 | 0.30 | 0.31 | 15 | 26 | 43 | 24 | 42 |
| JH-FK-43 | 2.0 | >10 | 31 | 22 | 14 | ND<sup>i</sup> | ND<sup>i</sup> |
| JH-FK-45 | 11 | NI<sup>h</sup> | 398 | 52 | 31 | ND<sup>i</sup> | ND<sup>i</sup> |
| JH-FK-46 | 0.23 | 0.31 | 0.49 | 2.9 | 1.7 | 1.5 | 1.3 |

<sup>a</sup> µg/mL
<sup>b</sup> *C. neoformans*
<sup>c</sup> *A. fumigatus*
<sup>d</sup> nM
<sup>g</sup> Values taken from Rivera, *et al.* mBio 2023
<sup>h</sup> No inhibition
<sup>i</sup> Not determined

Examination of only antifungal or immunosuppressive activity, however, is insufficient to evaluate the therapeutic potential of these derivatives. Previously, we have employed the therapeutic index (TI) score, a ratio of the immunosuppressive and antifungal activity relative to FK506, given by the equation 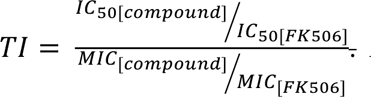. A larger TI suggests that an inhibitor is more biased towards antifungal activity than FK506 (TI=1). Of the fourteen derivatives we tested, thirteen of them showed a TI greater than 1, suggesting an improvement in the ratio of antifungal to immunosuppressive activity (**Table 1**; **Figure 3**). Only one C22-derivative, JH-FK-36 with a TI of 0.8, was biased more strongly towards immunosuppressive activity than antifungal activity. While JH-FK-38 and JH-FK-45 displayed improved TI (43 and 52, respectively) relative to JH-FK-08 (30), they showed only weak antifungal activity against *C. neoformans* (MIC80: 2.5 and 11 µg/mL, respectively). All fourteen compounds maintained some level of activity against *C. neoformans*, with activities ranging from 0.2 to 11 μg/mL. Seven of them (JH-FK-29, 30, 31, 35, 36, 40, and 46), demonstrated greater antifungal potency than JH-FK-08. They also displayed a strong retention of immunosuppressive activity, leading to TI of 0.8–52, where two of them (JH-FK-38 and 45) outperformed JH-FK-08 (TI: JH-FK-38=43, JH-FK-45=52 vs. JH-FK-08=30). The seven hydrazides selected for their size and hydrophilic character (JH-FK-29, 30, 31, 32, 33, 34, and 36) were strongly antifungal with all MIC80 values at or below 1.3 μg/mL, with four of them having stronger antifungal activity than JH-FK-08. The strong antifungal activity observed in assays supports our hypothesis that compounds with a small, hydrophilic substituent at C22, such as JH-FK-31 and JH-FK-36, would potently inhibit *C. neoformans*. However, they also retain strong immunosuppressive activities, resulting in modest TI scores. Although these small, hydrophilic substitutions might be well tolerated by *C. neoformans*, they may not sufficiently perturb human FKBP12-CaN binding to limit immunosuppressive activity.

**Figure 3.**
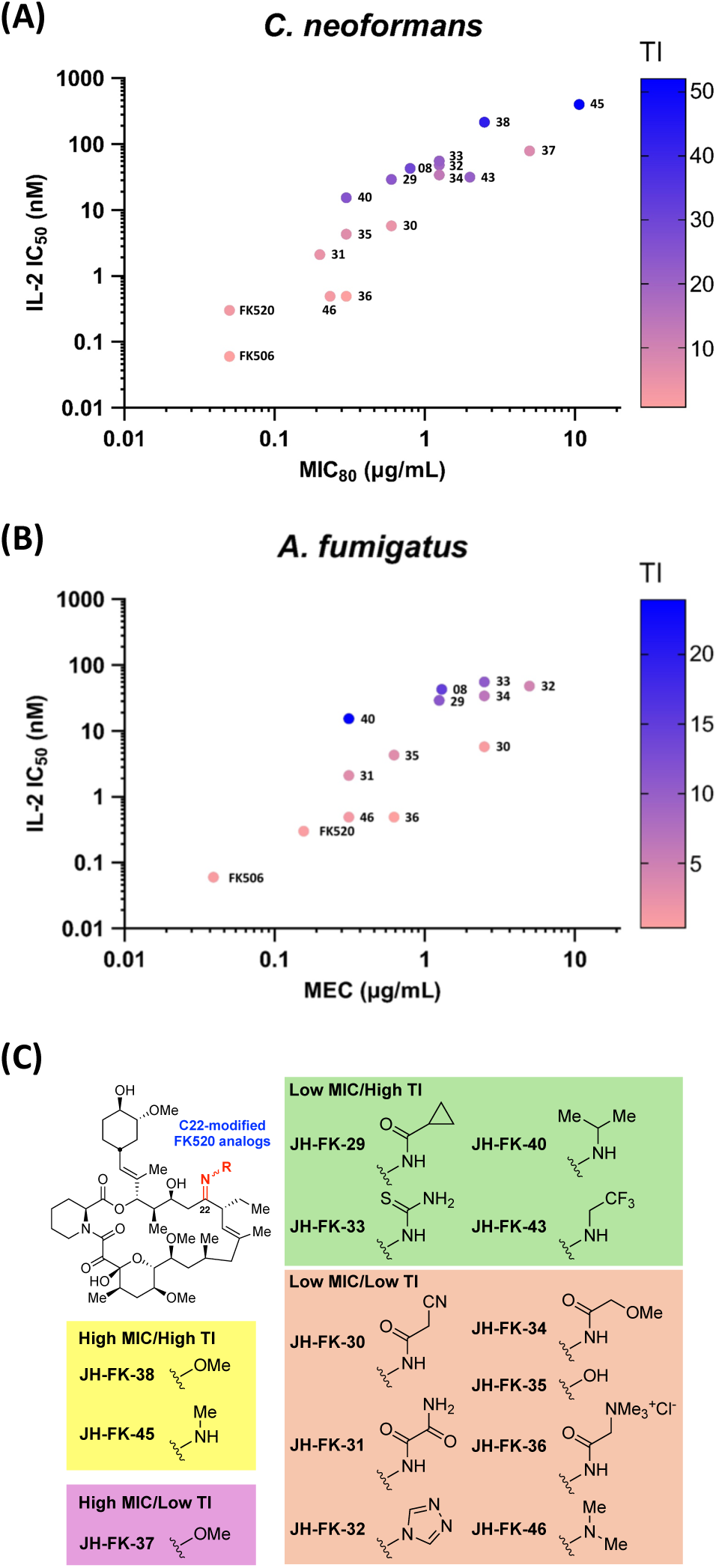
Scatter plots for the antifungal and immunosuppressive activity of C22-modified FK520 derivatives tested against *C. neoformans* (A) and *A. fumigatus* (B). (C) Summary of the impact of structural modifications on antifungal activity and TI. MIC: ≤ 2.5 µg/mL (Low) and > 2.5 µg/mL (High) against *C. neoformans*. TI: < 20 (Low) and ≥ 20 (High) against *C. neoformans*. For clarity, only the numerical identifiers of the compound codes are shown in (A) and (B). JH-FK-37, 38, 43, and 45 are excluded from (B) due to their lack of antifungal activity against *A. fumigatus*.

The seven oxime and hydrazone substitutions were designed to avoid potential steric clashes of C22 substitution with F88/H88. Although these compounds were designed to minimize deleterious interactions, oximes and hydrazones broadly mitigated antifungal activity, with only two of the seven oxime and hydrazone compounds having an MIC80 <1 μg/mL. Immunosuppressive activity was similarly mitigated, though one compound (JH-FK-46) maintained potent antifungal and immunosuppressive activity. Among the oximes and hydrazones tested for antifungal and immunosuppressive activity, two compounds (JH-FK-38 and JH-FK-45) demonstrated improved TI scores against *C. neoformans*. While both had limited antifungal activity (2.5 and 11 µg/mL, respectively), immunosuppressive activity was strongly reduced, leading to excellent TI scores. The docking model of JH-FK-45 on the human FKBP12-FK520- CaN complex revealed steric clashes between the methyl oxime group of JH-FK-45 and both the co-crystalized water and H88 (**Figure 4**). While the synthesized oximes and hydrazones were designed to mitigate steric clashes with target residues F88/H88, they appear to broadly reduce binding to both the human and *Cryptococcus* systems.

**Figure 4.**
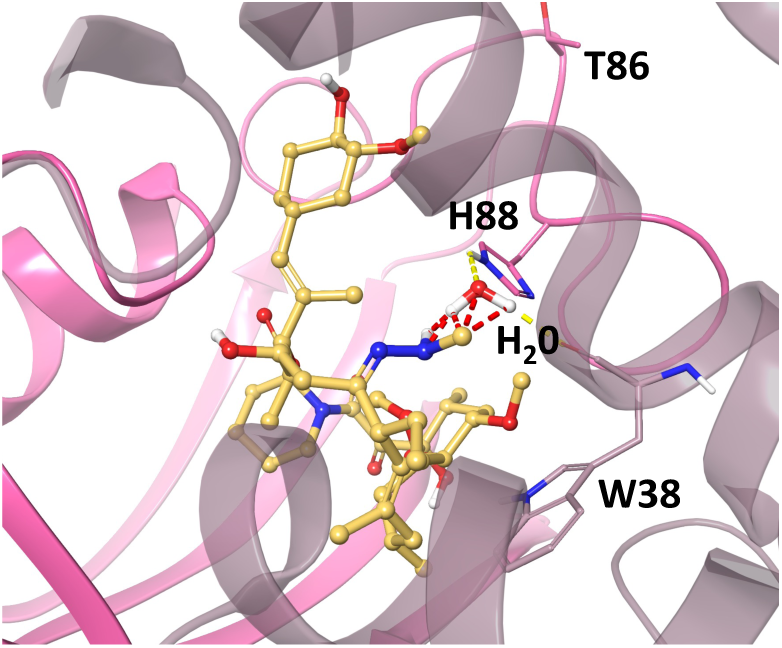
Docking model of JH-FK-45 (gold) on human FKBP12-FK520-CaN complex (PDB 7U0T). FKBP12 and CaN are colored in pink and dark pink, respectively. The co-crystallized water in the initial structure is represented. Hydrogen bonds and steric clashes are represented by yellow and red dashed lines, respectively.

Comparison with *Aspergillus* noted similar trends. All substitutions improved the therapeutic index for *A. fumigatus,* excluding JH-FK-36. Derivatives with limited antifungal activity against *C. neoformans* (JH-FK-37, 38, 43, and 45) similarly displayed poor or no antifungal activity against *A. fumigatus*. While both JH-FK-38 and JH-FK-45 displayed excellent TI scores against *C. neoformans*, neither was active against *A. fumigatus.* JH-FK-40 exhibited a good TI score against *C. neoformans*, but inferior to JH-FK-08. However, JH-FK-40 outperformed JH-FK-08 against *A. fumigatus.* The activity against *Cryptococcus* and *Aspergillus* strains appears broadly similar, but the two compounds with the best TI scores against *C. neoformans* (JH-FK-38 and 45) exhibited no observable antifungal activity against *A. fumigatus*.

*Additional in silico docking studies further implicate C32 as part of a wider hydrogen bonding network*.

Having explored *in silico* docking to design the new C22 derivatives, we employed the *in silico* approach towards other potential sites for modification. Comparison of crystal structures of human, *A. fumigatus*, *C. neoformans,* and *C. albicans* CaN complexed with FK506 or FK520 and FKBP12 noted potential differences near the C32 hydroxyl group (C32-OH) of FK506/FK520. The C32-OH has long been recognized to occupy a solvent-exposed region at the interface of CaN and FKBP12.^50^ Additionally, molecular dynamics simulations demonstrated that the C32-OH and nearby atoms of FK506 and APX879 have different interactions with human FKBP12 compared to fungal FKBP12.^51^ Examination of human CaN bound to FK520 and FKBP12 (**Figure 5A**) suggested that the C32-OH is part of an extensive hydrogen bonding network, functioning as a hydrogen bond acceptor for water. This water molecule, in turn, forms a hydrogen bond with H88, the residue that is also targeted for modification at C22. *A. fumigatus*, *C. neoformans,* and *C. albicans* complexes with FK506, however, interact differently with the C32-OH of FK506. In both *A. fumigatus* and *C. neoformans* (**Figure 5B** and **Figure 5C**, respectively), H88 is substituted with F88, which does not participate in the hydrogen bonding with the C32-OH of FK506. In *C. albicans*, I102 replaces H88, which also does not participate in hydrogen bonding connected to the C32-OH (**Figure 5D**). The presence of a hydrogen bonding network connecting the C32-OH of FK506/FK520 to H88 appears to be unique to the human system, as it does not appear in the fungal complexes.

**Figure 5.**
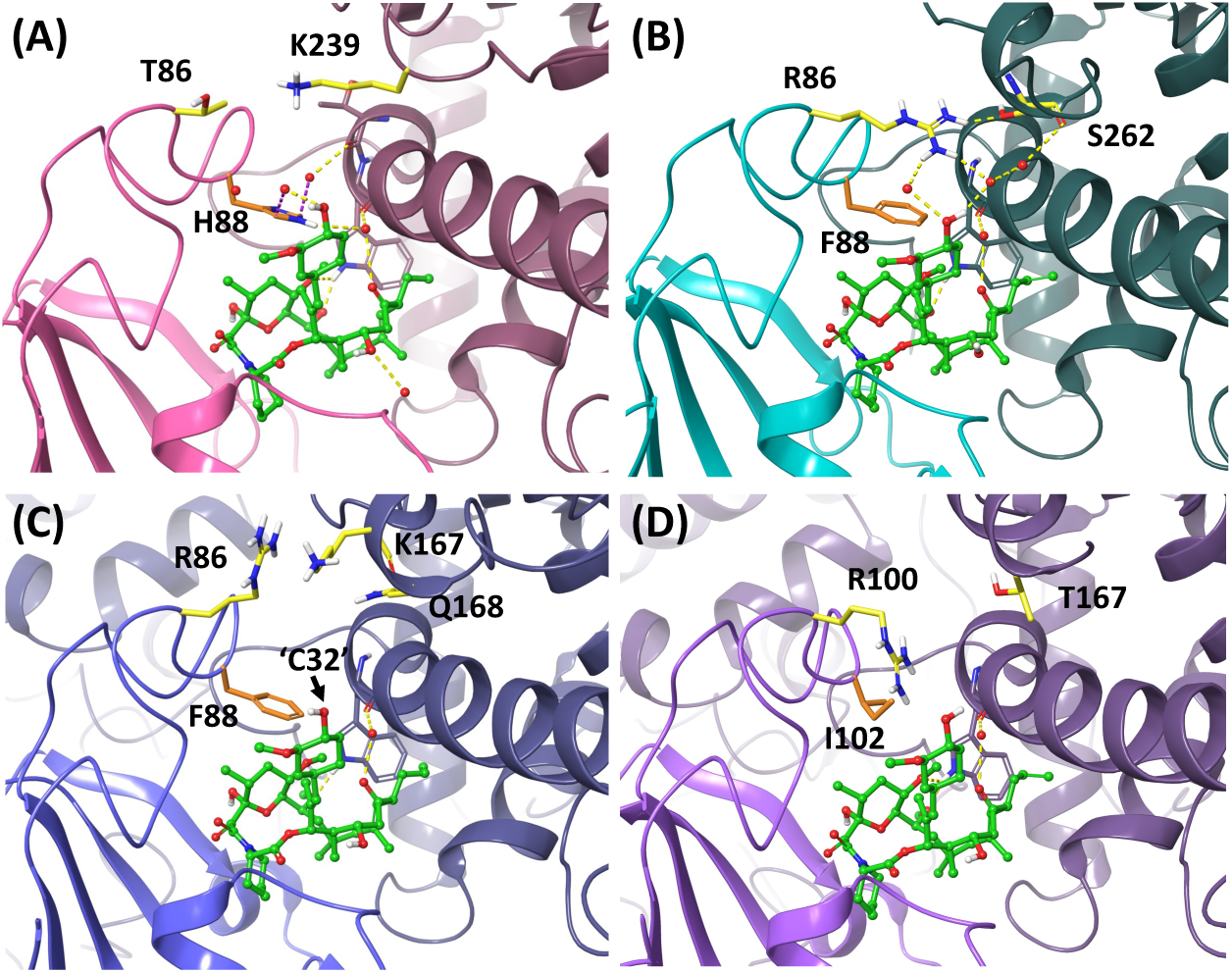
Binding-site regions surrounding the C32 hydroxyl group (C32-OH) of FK506 and FK520. The complex structures of (A) human (PDB 7U0T), (B) *A. fumigatus* (PDB 7U0U), (C) *C. neoformans* (PDB 6TZ8), and (D) *C. albicans* (PDB 6TZ6) are depicted in cartoon representation, respectively. FK506 and FK520 are colored in green. Co-crystallized water molecules are shown as red spheres. Hydrogen bonds and aromatic hydrogen bonds are represented by yellow and purple dashed lines, respectively. Brighter ribbon represents FKBP12, and darker chains represent CaN. The H88/F88/I102 residues are shown in orange. The key residues near the C32 position are shown in yellow and highlighted.

While the human complex has a hydrogen bond network connecting C32-OH-H88-CaN, fungi have alternative unique interactions from FKBP12 R86 with CaN. R86 is expected to interact with CaN S262 (PDB: 7U0U for *A. fumigatus*; **Figure 5B**). Alternatively, fungal R86 may interact with CaN via water-mediated hydrogen bonding. The C32-OH of FK506, however, engages in alternate hydrogen bonding modes in *A. fumigatus*, either hydrogen bonding with CaN P41 (PDB: 7U0U) through water bridging or direct hydrogen bonding through CaN E375 (PDB: 6TZ7). The residue T86 in human FKBP12 occupies the same space as fungal R86 (*A. fumigatus* and *C. neoformans*) or R100 (*C. albicans*). T86 is relatively small and positioned far away from the C32- OH of FK506/FK520. No interactions between the C32-OH and T86 are observed in the human FKB12-FK506-CaN complex. Therefore, the C32-OH of FK506/FK520 is a potentially exploitable site for developing non-immunosuppressive antifungal FK520 derivatives.

### Design and synthesis of C32-modified FK520 derivatives

Having identified the potential role of the C32-OH in human FKBP12-FK506/FK520-CaN binding, we decided to target the C32 position of FK520 in the development of non-immunosuppressive antifungal compounds. Because disrupting hydrogen bonding to the C32-OH may be significantly deleterious to human CaN binding while having minimal impact on fungal CaN binding, we proposed methylation of C32 in FK520 to generate the C32-methoxy derivative JH-FK-41 (**Figure 6A**). Expanding on the idea of interrupting hydrogen bonding, the C32- benzyloxy derivative JH-FK-42 could simultaneously disrupt the hydrogen bonding in the human system and introduce beneficial cation–π interactions with the fungal R86 residue. In doing so, C32-benzylation could reduce the immunosuppressive activity of FK520 while maintaining antifungal activity. Lastly, esterification at C32 with phthalic acid (JH-FK-44) could achieve similar disruption of hydrogen bonding in the human system as JH-FK-42, while also capturing additional cation–π interactions with the R86 in *C. neoformans* FKBP12 (**Figure 6B** and **6D**). As shown in **Figure 6C** and **6E**, the strong ionic interactions between the C32 phthalic acid group of JH-FK-44 and human CaN K239 (1.89 Å from the C32 phthalic acid group) are expected and may further destabilize the FKBP12-ligand-CaN complex, contributing to a loss of immunosuppressive activity. In contrast, *C. neoformans* Q168 (5.32 Å from the C32 phthalic acid group), which aligns with human CaN K239 in sequence, and K167 (6.54 Å from the C32 phthalic acid group) are not expected to form strong interactions with the phthalic acid group (**Figure 6B**). Therefore, we propose that these C32-modified compounds could provide a foundation for exploring C32 modifications in the development of antifungal, non-immunosuppressive FK520 derivatives.

**Figure 6.**
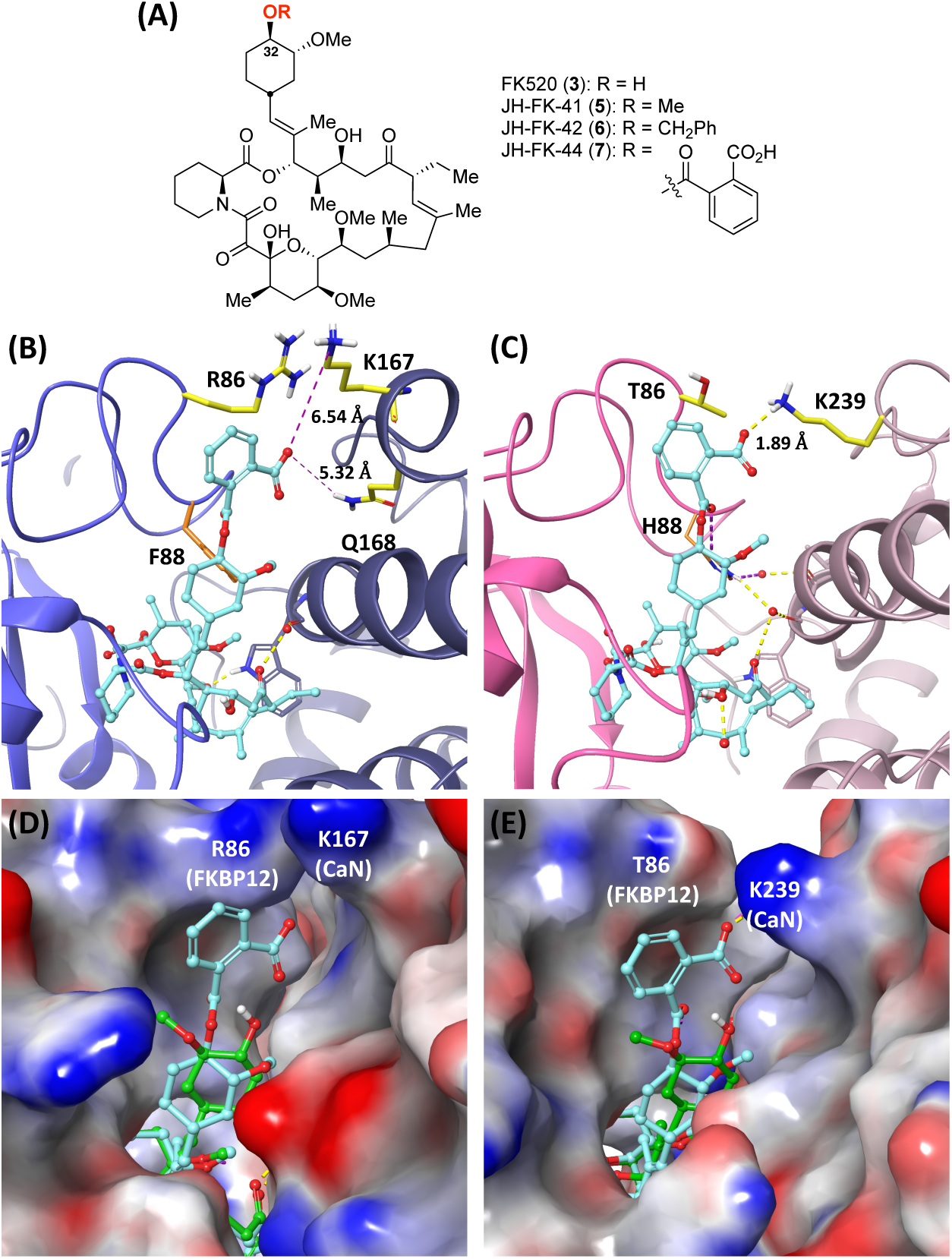
(A) Structures of proposed C32-modified FK520 derivatives JH-FK-41, 42, and 44. (B) Docking model of JH-FK-44 (**7**) bound to the *C. neoformans* CaN and FKBP12 complex (PDB 6TZ8). Brighter ribbon represents FKBP12, and darker chains represent CaN. The F88/H88 residues are shown in orange. The key residues close to the C32 position are shown in yellow and highlighted. Distances between the phthalic acid group and adjacent residues are shown. (C) Docking model of JH-FK-44 bound to human CaN and FKBP12 complex (PDB 7U0T). Hydrogen bonds and aromatic hydrogen bonds are represented by yellow and purple dashed lines, respectively. (D) Electrostatic surface representation of the docking model of JH-FK-44 on the *C. neoformans* complex and (E) the human complex. JH-FK-44 is shown in cyan. Co-crystalized FK506 and FK520 are shown in green.

To evaluate the antifungal activity of the proposed C32 modifications, we embarked on the synthesis of three C32-modified FK520 derivatives (**Figure 7**). Drawing from an earlier synthesis of C32-derivatized FK506,^52^ FK520 (**3**) was protected with TBSOTf to yield the C24/C32 di-TBS ether **8**. Selective deprotection of the C32 TBS group under acidic conditions afforded the C24 mono-TBS ether **9**, which was used as the common synthetic intermediate for all three derivatives. Methylation of **9** with trimethyloxonium tetrafluoroborate yielded the C24-OTBS/C32-OMe ether **10**. Benzylation of **9** with benzyl 2,2,2-trichloroacetimidate gave the C24-OTBS/C32-OBn ether **11**. Esterification of **6** with phthalic anhydride proceeded smoothly to afford the C24-OTBS/C32 *ortho-*benzoic ester **12**. TBS deprotection of **10**–**12** completed the synthesis of target compounds JH-FK-41 (**5**), JH-FK-42 (**6**), and JH-FK-44 (**7**).

**Figure 7.**
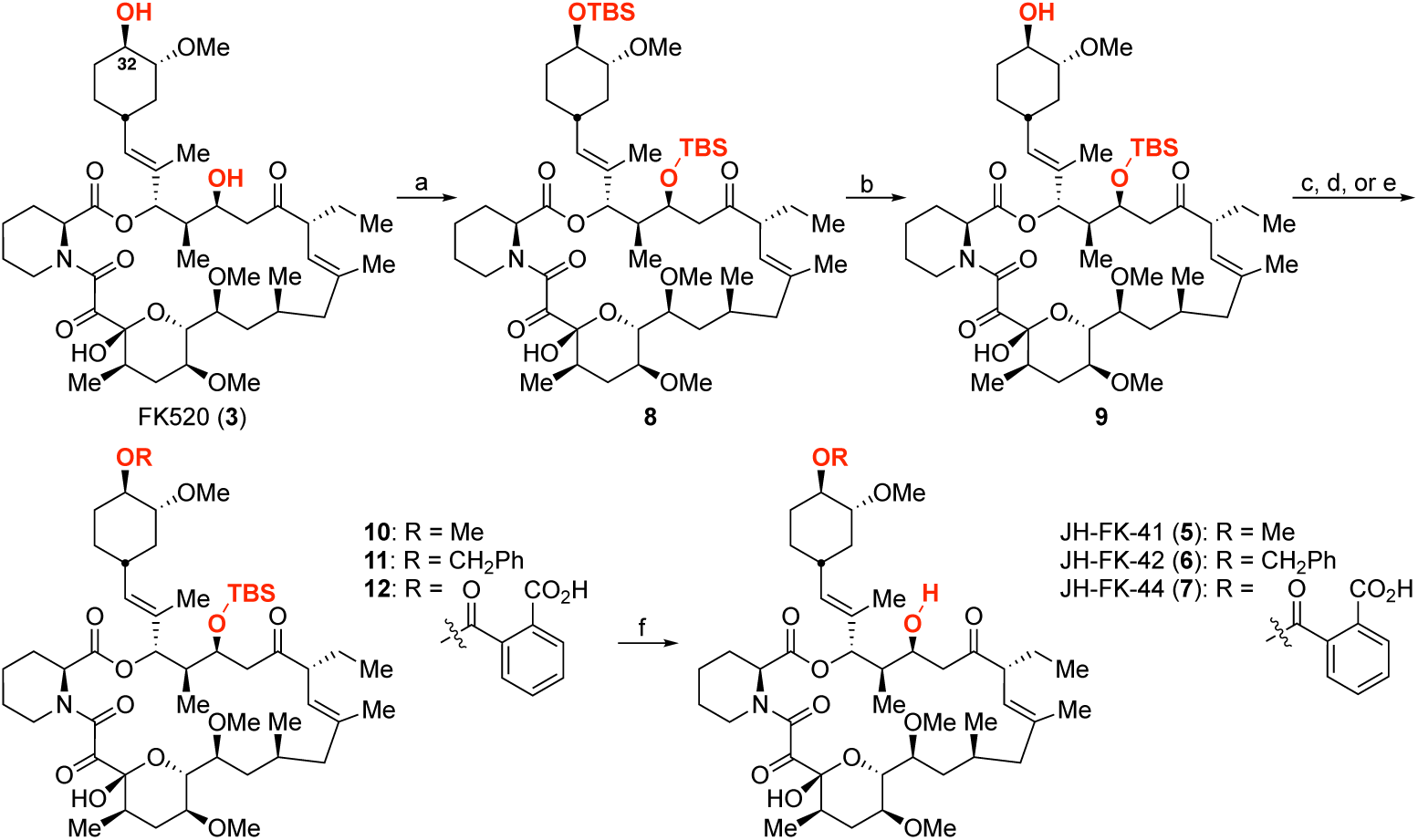
The synthesis of C32-modified FK520 derivatives. (a) TBSOTf, 2,6-lutidine, CH2Cl2, 0 °C, 1 h, 72%; (b) *p*-TsOH·H2O, MeOH/CH2Cl2, 25 °C, 5 h, 75%; (c) proton sponge, 4Å MS, Me3OBF4, CH2Cl2, 25 °C, 18 h, 76%; (d) benzyl 2,2,2-trichloroacetimidate, TfOH, CH2Cl2, 25 °C, 5 h, 24%; (e) phthalic anhydride, DMAP, pyridine, 25 °C, 24 h, 38%; (f) HF·pyridine, MeCN, 25 °C, 20 h, JH-FK-41 (**5**): 45%, JH-FK-42 (**6**): 35%, JH-FK-44 (**7**): 22%.

### Biological characterization of C32-modified FK520

The antifungal activity of the C32-modified FK520 derivatives JH-FK-41, 42, and 44 was assessed via fungal growth in a series of MIC assays against *C. neoformans, C. albicans*, and *A. fumigatus*. Immunosuppressive activities and TI scores were calculated according to the IL-2 assay and equation previously used for C22-modified derivatives (**Table 2**; **Supplementary Figure S2**). Both JH-FK-41 and JH-FK-42 showed potent antifungal activity (0.052–0.50 μg/mL) against *C. neoformans* and *C. albicans*, although they were strongly immunosuppressive as well. JH-FK-41 also demonstrated strong antifungal activity against *A. fumigatus* (0.16 μg/mL), indicating it possesses broad efficacy. JH-FK-42 has much weaker antifungal activity against *A. fumigatus* than *C. neoformans* or *C. albicans* (MEC 1.3 μg/mL vs. 0.50 and 0.13 μg/mL, respectively). Immunosuppressive activity was more strongly impacted than antifungal activity in all three species, in line with our proposed strategy. JH-FK-44 showed weaker antifungal activity than either JH-FK-41 or JH-FK-42, with MICs of 3.2 and 1.1 μg/mL against *C. neoformans* and *C. albicans*, respectively, and no activity against *A. fumigatus.* However, as predicted by the docking model predicted (**Figure 6**), JH-FK-44 exhibited only very weak immunosuppressive activity, with an IL-2 IC50 of 129 nM, compared to FK506 (IC50: 0.04–0.08 nM). Consequently, JH-FK-44 exhibited the most favorable TI score (57 against *C. neoformans* and 93 against *C. albicans*) thus far, surpassing lead compound JH-FK-08 (30). The significantly weak immunosuppressive activity of JH-FK-44 compared to FK506 or FK520 suggests that C32 modification warrants further studies and may offer a great opportunity for the development of non-immunosuppressive antifungal agents.

**Table 2.**
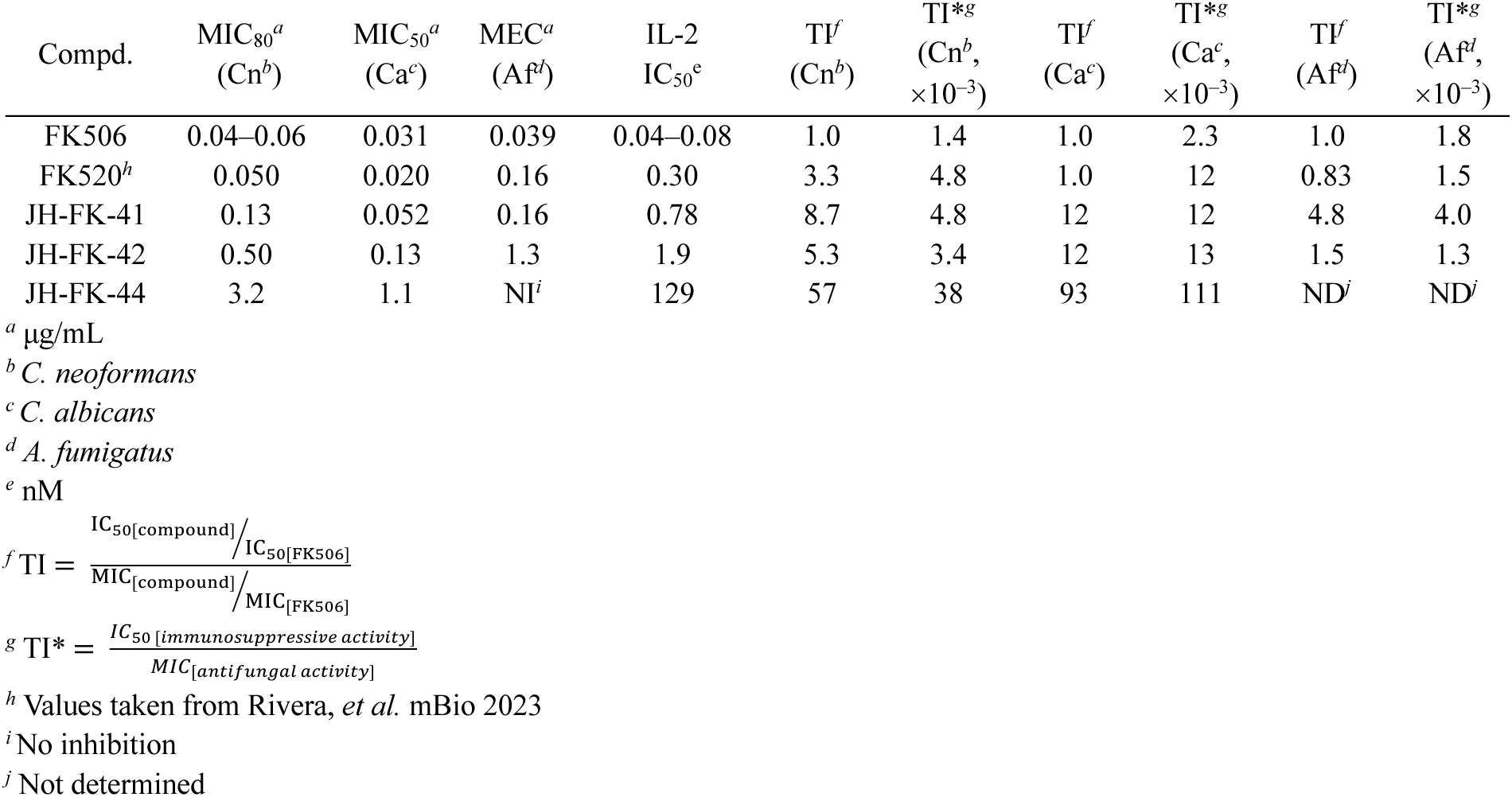
Antifungal and immunosuppressive activity and therapeutic index (TI and TI*) scores of C32- modified FK520 derivatives.

| Compd. | MIC<sub>80</sub><sup>a</sup><br>(Cn<sup>b</sup>) | MIC<sub>50</sub><sup>a</sup><br>(Ca<sup>c</sup>) | MEC<sup>a</sup><br>(Af<sup>d</sup>) | IL-2<br>IC<sub>50</sub><sup>e</sup> | TI<sup>f</sup><br>(Cn<sup>b</sup>) | TI*<sup>g</sup><br>(Cn<sup>b</sup>,<br>×10<sup>-3</sup>) | TI<sup>f</sup><br>(Ca<sup>c</sup>) | TI*<sup>g</sup><br>(Ca<sup>c</sup>,<br>×10<sup>-3</sup>) | TI<sup>f</sup><br>(Af<sup>d</sup>) | TI*<sup>g</sup><br>(Af<sup>d</sup>,<br>×10<sup>-3</sup>) |
| --- | --- | --- | --- | --- | --- | --- | --- | --- | --- | --- |
| FK506 | 0.04–0.06 | 0.031 | 0.039 | 0.04–0.08 | 1.0 | 1.4 | 1.0 | 2.3 | 1.0 | 1.8 |
| FK520<sup>h</sup> | 0.050 | 0.020 | 0.16 | 0.30 | 3.3 | 4.8 | 1.0 | 12 | 0.83 | 1.5 |
| JH-FK-41 | 0.13 | 0.052 | 0.16 | 0.78 | 8.7 | 4.8 | 12 | 12 | 4.8 | 4.0 |
| JH-FK-42 | 0.50 | 0.13 | 1.3 | 1.9 | 5.3 | 3.4 | 12 | 13 | 1.5 | 1.3 |
| JH-FK-44 | 3.2 | 1.1 | NI<sup>i</sup> | 129 | 57 | 38 | 93 | 111 | ND<sup>j</sup> | ND<sup>j</sup> |

<sup>a</sup> µg/mL
<sup>b</sup> *C. neoformans*
<sup>c</sup> *C. albicans*
<sup>d</sup> *A. fumigatus*
<sup>e</sup> nM
<sup>h</sup> Values taken from Rivera, *et al.* mBio 2023
<sup>i</sup> No inhibition
<sup>j</sup> Not determined

### NMR binding studies confirm the suggested bonding mode of C32-derivatives

Previously reported NMR titrations of FK506 into human and *A. fumigatus* FKBP12 proteins, monitored by ^1^H-^15^N HSQC experiments, enabled residue level characterization of the protein responses to ligand binding in solution.^51,53^ FK520 binding to the apo proteins induced chemical shift perturbations similar to those observed with FK506 binding. Specifically, in *A. fumigatus*, perturbations at least one standard deviation above the average were observed for residues I25, H26, Y27, T49, V56, I57, V102, and E103, while a different set of residues in human FKBP12–namely, V25, S40, R43, G52, I57, E62, A65, and F100–exhibited the largest chemical shift variations between the apo and FK520 bound forms. These results reinforce the observation that human and fungal FKBP12 proteins display altered electronic environments around these residues upon ligand binding, indicating differential binding determinants.

To validate our *in silico* models of C32 modifications, we compared these FK520 binding results with a series of ^1^H-^15^N HSQC NMR binding studies of human and *A. fumigatus* FKBP12 with three C32-modified compounds: JH-FK-41, JH-FK-42, and JH-FK-44 (**Figure 8**). Overall, JH-FK-41, JH-FK-42, and JH-FK-44 binding to the apo human and *A. fumigatus* FKBP12 proteins induced chemical shift perturbations similar to those seen with FK520 binding, with the same residues showing the largest perturbations. These data suggest that all of these inhibitors bind in the same overall orientation as FK520. Subtracting the chemical shift perturbations of each residue observed for FK520 binding from the corresponding perturbations measured in the C32-modified compounds revealed that, as expected, the major differences occur primarily at residues T86 and H88 in human and R86 and F88 in *A. fumigatus*. These findings support the hypothesis that C32 modifications disrupt the hydrogen bonding network in the human protein while introducing favorable cation–π interactions with the fungal R86 residue. Additionally, the C32-methoxy modification resulted in a measurable difference at H88 in human FKBP12 that was not observed at *A. fumigatus* F88.

**Figure 8.**
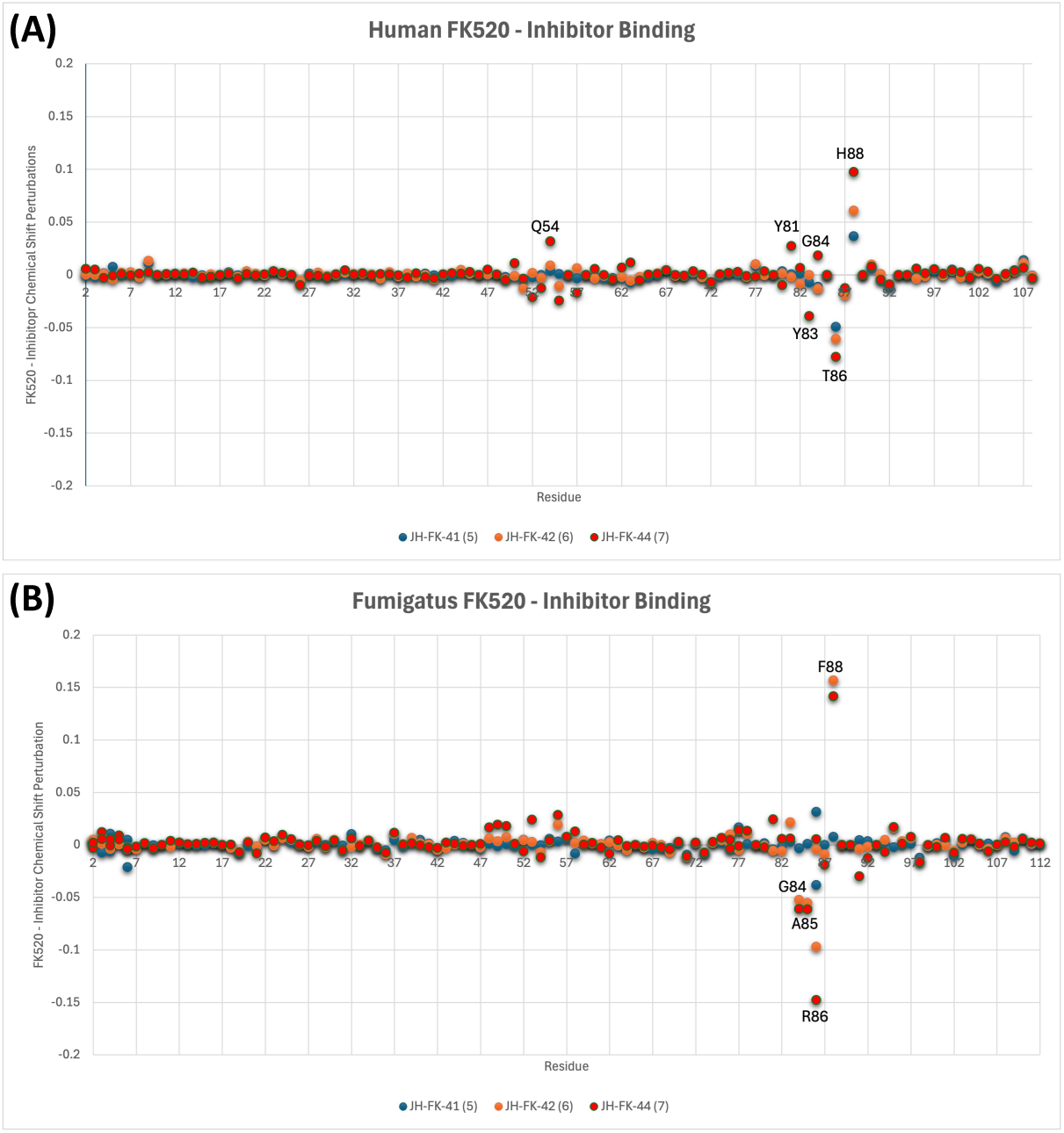
^1^H-^15^N HSQC NMR binding of selected C32-modified FK520 derivatives (JH-FK-41, JH-FK-42, and JH-FK-44) with (A) human and (B) *A. fumigatus* FKBP12. Chemical shifts perturbations upon binding to JH-FK-41, JH-FK-42, and JH-FK-44 that are significantly different from FK520 perturbations are labeled.

### Cytotoxicity of selective C22 and C32 derivatives of FK520

We have generated a series of C22 and C32 derivatives of FK520 with improved TI scores. Therapeutic application, however, is also dependent on the level of general cytotoxicity. Weak antifungal activity may require higher concentrations for efficacy, potentially at concentrations toxic to humans. To assess the cytotoxicity of C22 and C32 derivatives of FK520, we incubated mouse fibroblast NIH3T3 cells with C22 derivatives (JH-FK-43, 45, and 46) and C32 derivatives (JH-FK-41, 42, and 44). After 24 h-incubation, the cell viability was assessed using CellTiter-Glo luminescent cell viability assay (**Supplementary Figure S3**). Two of the six compounds we tested, JH-FK-43 and JH-FK-44, were more cytotoxic than FK520. JH-FK-46 displayed comparable cytotoxicity to FK520, while JH-FK-41, 42, and 45 were less cytotoxic. JH-FK-41, 42, and 45 displayed cell viability >80 or 90% at 10 µM. The significantly low cytotoxicity of these C22 and C32 derivatized compounds is promising for further development, as it suggests the potential for non-cytotoxic FK520-based antifungals.

## Conclusions

In this study, we have leveraged *in silico* models of FK506/FK520 binding to human and fungal FKBP12 and calcineurin to expand the chemical space around the C22 carbonyl group of FK506/FK520 and design a novel series of C32-derivatives aimed at developing non-immunosuppressive antifungal agents. Our design strategy for the C22 position focused on replacing the co-crystallized water molecules near C22, which are part of a larger hydrogen bonding network connecting FKBP12, FK520, and CaN. In parallel, we explored the use of carbonyl bioisosteres, such as hydrazones and oximes, to test the hypothesis that C22 hydrazones or oximes could enhance antifungal activity by improving hydrophobic interactions with F88 in fungal FKBP12. Additionally, examination of human and fungal FKBP12-CaN crystal structures bound to FK506 or FK520 revealed exploitable differences around the C32-hydroxyl group. This led to the synthesis of three C32 derivatives and the discovery of JH-FK-44, which exhibited the highest therapeutic index against *C. neoformans*. The promising *in vitro* results of JH-FK-44 suggest that C32 modifications could play a key role in the future development of non-immunosuppressive antifungal calcineurin inhibitors.

## Materials and Methods

### In silico docking

Ligand docking was performed using Schrödinger Maestro, release 2024-1, operated on Windows 10 (Schrödinger, LLC: New York, NY, USA, 2024). Ligands were either sketched using the 2D sketcher module or downloaded as .sdf files from the PubChem database (https://pubchem.ncbi.nlm.nih.gov/) and prepared using the LigPrep module. All possible ionization states, including the neutral form, were generated under physiological pH conditions for use in docking. The X-ray crystal structures of human FKBP12-FK520-CaN complex (PDB ID: 7U0T), *A. fumigatus* FKBP12-FK506-CaN complex (PDB ID: 7U0U), *C. neoformans* FKBP12- FK506-CaN complex (PDB 6TZ8), *C. albicans* FKBP12-FK506-CaN complex (PDB 6TZ6), and human FKBP12-APX879 complex (PDB 6VCU) were downloaded from the RCSB Protein Data Bank (https://www.rcsb.org/). The downloaded structures were prepared using the Protein Preparation module, with missing side chains and loops filled using the Prime module. All unnecessary molecules were removed. The grid boxes were defined based on the complexed inhibitors, with grid box kept at default. All grid boxes were set to defaults. Docking was performed using XP-Glide in the Ligand Docking module under default settings, with the ‘sampling macrocycles’ option selected. A maximum of 20 poses were generated with post-docking minimization, and the results were manually reviewed and selected. The residue numbering for FKBP12 follows the UniProt sequence, while the numbering for the calcineurin segment is based on the residue numbers from the PDB.

### Synthesis of C22- and C32-modified FK520 derivatives

Detailed synthesis and characterization of C22- and C32-modified FK520 derivatives is described in Supporting Information.

### Antifungal susceptibility assay

Strains were grown overnight in YPD broth at 30°C and pelleted by centrifugation. Pellets were washed two times with sterile water and then resuspended in RPMI-1640 medium (Sigma-Aldrich). OD600 of the cell suspension was measured using a BioRad SmartSpec 3000, and the used to inoculate the 96-well plate at a final density of OD600 = 0.0005. Microdilution broth assays were conducted as previously described in 96-well plates using modified CLSI protocols M27-A3 for antifungal susceptibility testing in yeasts.^42^ Antifungal assays of *C. albicans* strain SC5314 were carried out at 30 °C for 48 h, in the presence of 2 µg/ml fluconazole in order to sensitize the strain to FK506. Antifungal assays of *C. neoformans* strain H99 were incubated at 37 °C for 72 h. At the end of incubation, plates were read with a Molecular Devices iD3 Spectra plate reader in addition to visual scoring.

### *A. fumigatus* strains, culture conditions and *in vitro* drug susceptibility assays

The wild-type *A. fumigatus* (*akuB*^KU80^) and the immunophilin AfFKBP12 deletion (Δ*Affkbp12*) strains were utilized for FK506/FK520 analog susceptibility assays. Strains were cultured on glucose minimal medium (GMM) agar for 5 days at 37 °C to collect conidia for the assays. Broth dilution antifungal susceptibility testing was performed following the Clinical Laboratory Standards Institute (CLSI) guidelines with slight modifications, using RPMI-1640 medium. Conidia were harvested in 0.05% Tween 80, counted to obtain a spore concentration of 10^4^/ml, and further diluted to obtain 5×10^2^/mL in each well (200 µL) of a 96-well plate. The effects of FK506/FK520 and various analogs on the growth of different strains were tested at concentrations ranging from 0 to 20 µg/mL. Growth inhibition was assessed by microscopic observation after 2- day growth at 37 °C, and minimum effective concentration (MEC) results were interpreted. Bright- field photomicrographs were captured using a VWR inverted microscope equipped with a Canon T5i digital camera.

### *In vitro* immunosuppressive activity

Spleens and lymph nodes were collected from C57B/L6 mice and filtered through a 40 µm cell strainer. Red blood cells were lysed using ACK lysis buffer. Pan-CD4^+^ T cells were enriched using a MagniSort mouse CD4 T-cell kit (eBioscience) according to the manufacturer’s protocol. Thereafter, CD4^+^ CD25^−^ CD44^lo^ CD62L^hi^ naive T cells were sorted by FACS using a MoFlo Astrios or MoFlo XDP cell sorter (Beckman Coulter). Naïve CD4^+^ T cells were cultured in Iscove’s Modified Dulbecco’s medium supplemented with 10% FBS, penicillin (10 U/mL), streptomycin (10 µg/mL), glutamine (2 mM), gentamycin (50 µg/mL), and *β*-mercaptoethanol (55 µM). Cells were cultivated with serially diluted drugs in DMSO. Cells were cultured on anti-hamster IgG-coated plates in the presence of hamster anti-CD3 epsilon and anti-CD28 antibodies (BD Biosciences), neutralizing anti-IL-4 antibody (eBioscience), recombinant IL-12 (10 ng/mL), and recombinant IL-2 (50 U/mL) for 72 h. Cells were cultured with phorbol 12-myristate 13-acetate, ionomycin, and GolgiStop for the final 4 h. Cells were surface stained with viability dye eFluor 506 and FITC conjugated anti-CD4 antibody in PBS. Cells were fixed using an eBioscience Fix/Perm kit. Intracellular staining was conducted with PE-conjugated anti-IL-2 antibody. Cells were resuspended in a MACS buffer and examined via flow cytometry with a BD FACSCantoII and FlowJo (BD Biosciences).

### 1H-15N HSQC NMR binding studies

Procedure adapted from Gobeil *et al., 2021*.^51^*A. fumigatus* and *H. sapiens* FKBP12 constructs were obtained from GenScript (Piscataway, NJ) in the pET-15b vector. The plasmids, containing the proteins with a His6-tag at the *N*-terminus and a thrombin cleavage site, were transformed into *E. coli* BL21(DE3) cells and plated on LB-agar containing ampicillin (100 µg/mL). For NMR inhibitor titrations, uniformly [^15^N]-labeled proteins were overexpressed at 25 °C in modified M9 minimal medium containing 1 g/L ^15^NH4Cl (Cambridge Isotopes, Tewksbury, MA). Cells were propagated to an OD600 of 0.6 and induced with 1 mM Isopropyl *β*-D-1-thiogalactopyranoside (IPTG) for 4 h. Cells were then harvested by centrifugation at 4 °C for 15 min at 6,000 g and the pellet stored at −20 °C until purification. The frozen pellets were resuspended in 30 mL of lysis buffer (50 mM Na2HPO4, 500 mM NaCl, pH 8.0) supplemented with 1 mL of protease inhibitor cocktail (Sigma, St. Louis, MO) and 1 mM phenylmethanesulfonyl fluoride (PMSF). Lysis was performed using three cycles of 30 s of sonication at 12 W power, with a 2-min rest interval on ice. Lysate was clarified by centrifugation at 20,000 g for 15 min at 4 °C, and the supernatant was loaded onto a Ni-NTA column. Protein was eluted using a gradient of 0 to 1 M imidazole. Fractions containing FKBP12 were identified by SDS-PAGE and stained with Coomassie Blue staining, then pooled and dialyzed four times for 1 h each into 50 mM Na2HPO4, 500 mM NaCl, pH 8.0 buffer to remove imidazole. The 6×-His tag was cleaved for 16 h at 4 °C using 1U of thrombin per 100 µg of total protein (GE Healthcare). The cleaved proteins were then passed through a Ni-NTA column to remove the cleaved 6×-His tag and any uncleaved protein. Pure protein was obtained via size-exclusion chromatography using a Sephacyrl S100HR XK26/60 FPLC column. Typical yields were 20 mg/L of > 98% pure protein, as determined by SDS-PAGE with Coomassie Blue staining. [^15^N]-labeled samples were concentrated to approximately 0.4–0.7 mM and buffer exchanged into NMR buffer (20 mM Na2HPO4, 100 mM NaCl, 0.02% NaN3 and 5% D2O at pH 6.0) using a 3,000 MWCO Amicon concentrator. All NMR experiments were performed at 25 °C on an 800MHz Agilent NMR spectrometer equipped with a triple resonance cryoprobe. ^15^N, ^13^C, and ^1^H backbone and sidechain resonances for the FKBP12 proteins from *A. fumigatus* and human were assigned for comparative evaluation in both the apo, FK520, JH-FK-41, JH-FK-42, and JH-FK-44 bound forms. Inhibitors were titrated into ^15^N stable isotope-labelled FKBP12s (ratio FKBP12:inhibitor of 1:0, 1:0.25, 1:0.5, 1:0.75, 1:1, 1:2) to investigate conformational changes upon complex formation. The ^1^H-^15^N HSQC NMR spectrum imparts a cross peak for each amide bond in the protein. Cross peaks were compared to determine the impact of interactions.

### Cytotoxicity assay

Mouse fibroblast NIH3T3 cells were purchased from Duke Cell Culture Facility (Durham, NC, USA). Cells were maintained in a humidified incubator at 37 °C with 5% CO2 in DMEM (GIBCO- 11995) supplemented with 10% heat-inactivated fetal bovine serum (#10082147, ThermoFisher) and antibiotics (streptomycin, 10,000 UI/ml and penicillin, 10,000 UI/ml, #15140122, ThermoFisher). Cells were seeded at 3,000–4,000 cell/well in 96 well plate (Corning®, cat# 3903) overnight. The indicated compounds were then added to the cells, and the viability was measured 24 or 48 h after compound treatment using CellTiter-Glo luminescent cell viability assay (Promega) according to the manufacturer’s instructions. Briefly, 15 µL of CellTiter-Glo substrate was added to cells cultured in a 96-well plate with 100 µl of media, followed by 10-min shaking. The signal intensity was measured using a chemiluminescence plate reader.

## Data, Materials, and Software Availability

All study data are included in the article and/or *SI Appendix*.

## Acknowledgments

We thank Dr. Benjamin Bobay (Duke University NMR Center and Department of Radiology, Duke University) for his helpful comments on the manuscript. This research is supported by NIH/NIAID R01 grant AI172451-02 and R56 grant AI112595-05 and the Gilhuly Accelerator Fund. M.C. acknowledges support from R01 GM115474. This material is based upon work supported by the National Science Foundation Graduate Research Fellowship under Grant No. DGF 2139754. Joseph Heitman is co-director and fellow of the CIFAR Program Fungal Kingdom: Threats & Opportunities.

## Author contributions

J.H., J.H., and P.R.J. designed research; P.A.D., P.J., G.N., H.J., A.R., A.F.A., E.P., T.-C.L. J.W., R.A.V., and P.R.J. performed research; P.A.D., P.J., G.N., H.J., A.R., A.F.A., E.P., T.-C.L. M.C., J.W., J.-T.A.C., R.A.V., H.-J.P., W.J.S., P.R.J., J.H., and J.H. analyzed data; P.A.D, P.J., G.N., J.H., and J.H. wrote the paper with contributions from all authors.

