## Supplementary Information for "Structure-guided design and synthesis of C22- and C32-modified FK520 analogs with enhanced activity against human pathogenic fungi"

**Table of Contents**

1. General methods for chemical synthesis
2. General synthetic procedures A, B, and C
3. Synthetic procedures and analytical data
4. Supplementary Table S1. Synthesis and structure of C22-modified FK520 derivatives
5. Supplementary Figure S1. Immunosuppressive activity of JH-FK-29, 30, 31, 32, 33, 34, 35, 36, 37, 38, and 40
6. Supplementary Figure S2. Immunosuppressive activity of JH-FK-41, 42, 43, 44, 45, and 46
7. Supplementary Figure S3. Cytotoxicity data for C22- or C32-modified FK502 derivatives
8. Copy of $^{1}H$ and $^{13}C$ NMR spectra
9. References

### 1. General methods for chemical synthesis

FK520 was purchased from APIChem Technology. Unless otherwise stated, all reagents were purchased from commercial suppliers (Sigma–Aldrich, Acros, Fisher, Alfa Aesar, TCI, or Ambeed) and used without further purification. All solvents were American Chemical Society (ACS) grade or better and used without further purification. Analytical thin layer chromatography (TLC) was performed with glass-backed silica gel (60 Å) plates with fluorescent indication (Whatman). Visualization was accomplished by UV irradiation at 254 nm and/or by staining with *p*-anisaldehyde solution or potassium permanganate solution followed by heating. Flash column chromatography was performed by using silica gel (particle size 230−400 mesh, 60 Å) purchased from Silicycle. All $^{1}H$ NMR spectra were recorded with a Bruker 500 (500 MHz) spectrometer in acetone-$d_6$. All NMR δ values are given in parts per million (ppm) and are referenced to the residual isotopomer solvent signals: acetone-$d_6$: δ = 2.05 ppm for $^{1}H$ NMR spectra and δ = 29.84 ppm for $^{13}C$ NMR spectra. Coupling constants ($J$) are given in Hertz (Hz) and multiplicities are indicated using the conventional abbreviation (s = singlet, d = doublet, t = triplet, q = quartet, m = multiplet or overlap of non-equivalent resonances, br = broad). Electrospray ionization (ESI) mass spectrometry (MS) was recorded with an Agilent 6224 series (LC/MS–TOF) spectrometer to obtain the molecular masses of the compounds.

**2. General synthetic procedures A, B, and C**

(1) *General Procedure A.* To a solution of FK520 in MeOH was added hydrazide or hydrazine (6 eq) and a catalytic amount of TFA (0.1 eq). The reaction was stirred at 25 °C for 48 h. The reaction mixture was concentrated *in vacuo* and reconstituted in EtOAc. The reaction mixture was washed with $H_2O$, brine, and 10% aqueous $NaHCO_3$ solution. The organic fraction was dried over anhydrous $Na_2SO_4$ and concentrated *in vacuo*. The residue was purified by column chromatography (silica gel, $CH_2Cl_2$/MeOH or hexanes/acetone) to afford the desired product.

(2) *General Procedure B.* To a solution of FK520 in EtOH was added hydroxylamine hydrochloride or *O*-methylhydroxylamine hydrochloride (1.2 eq) under inert atmosphere. The resulting mixture was stirred for 10 min and treated with NaOAc (1.5 eq). The reaction mixture was stirred at 25 °C for 24 h. The reaction mixture was concentrated *in vacuo*, reconstituted in EtOAc, and washed with $H_2O$. Combined organic phases were dried over anhydrous $Na_2SO_4$ and concentrated *in vacuo*. The residue was purified by column chromatography (silica gel, hexanes/acetone, 2/1) to afford the desired hydrazone.

(3) *General Procedure C.* To a solution of C24-TBS protected FK520 analog in acetonitrile was added HF·pyridine. The reaction mixture was stirred at 25 °C for 20 h. The reaction was quenched by an addition of saturated aqueous $NaHCO_3$ and the reaction mixture was extracted with $CH_2Cl_2$. The organic phase was dried over anhydrous $Na_2SO_4$ and condensed *in vacuo*. The residue was purified by column chromatography (silica gel, hexanes/EtOAc) to afford JH-FK-41 (**5**), JH-FK-42 (**6**), and JH-FK-44 (**7**).

### 3. Synthetic procedures and analytical data

**JH-FK-29.** Following *General Procedure A*, FK520 (40.0 mg, 0.051 mmol) was reacted with cyclopropanecarbohydrazide (30.6 mg, 0.306 mmol) and TFA (0.4 µL, 0.005 mmol) in MeOH (0.5 mL). The residue was purified by column chromatography (silica gel, hexanes/acetone, 2/1 to 3/2) to yield JH-FK-29 (33.9 mg, 76%) as a white solid: $^1$H NMR (500 MHz, acetone-$d_6$) δ 10.00 (s, 1H) 5.53 (d, $J$ = 9.3 Hz, 1H), 5.38 (d, $J$ = 15.4 Hz, 1H), 5.16 (s, 2H), 4.89 (s, 1H), 4.40–4.25 (m, 3H), 4.04 (d, $J$ = 17.7 Hz, 1H), 3.64 (s, 1H), 3.53 (t, $J$ = 10.1 Hz, 1H), 3.36 (br s, 6H), 3.33 (s, 3H), 3.23 (q, $J$ = 8.7 Hz, 1H), 3.03–2.95 (m, 2H), 2.41 (d, $J$ = 9.9 Hz, 1H), 2.40–2.21 (m, 4H), 2.19–2.09 (m, 5 Hz), 2.02–1.95 (m, 3H), 1.90–1.85 (m, 2H), 1.81–1.76 (m, 3H), 1.67 (s, 3H), 1.64 (s, 3H), 1.60–1.48 (m, 6H), 1.39–1.27 (m, 6H), 1.15–1.06 (m, 3H), 0.98 (d, $J$ = 7.2 Hz, 5H), 0.90–0.83 (m, 6H), 0.80–0.76 (m, 1H); HRMS (ESI) $m/z$ 874.5423 [(M + H)$^+$ calcd for $C_{47}H_{75}N_3O_{12}$ 874.5424].

**JH-FK-30.** Following *General Procedure A*, FK520 (40.0 mg, 0.051 mmol) was reacted with cyanoacetohydrazide (30.3 mg, 0.306 mmol) and TFA (0.4 µL, 0.005 mmol) in MeOH (0.5 mL) The residue was purified via column chromatography (hexanes/acetone, 2/1 to 1/1) to yield JH-FK-30 (37.2 mg, 84%) as a white solid: $^1$H NMR (500 MHz, acetone-$d_6$) δ 10.30 (s, 1H), 5.47 (d, $J$ = 3.9 Hz, 1H), 5.17 (t, $J$ = 8.0 Hz, 2H), 5.12–5.07 (m, 1H), 4.44 (s, 1H), 4.35 (s, 1H), 4.33 (d, $J$ = 14.5 Hz, 1H), 4.03 (d, $J$ = 12.1 Hz, 1H), 3.96 (s, 1H), 3.86 (s, 1H), 3.65 (s, 1H), 3.53 (d, $J$ = 9.9 Hz, 2H), 3.43–3.38 (m, 2H), 3.37 (s, 3H), 3.36 (s, 4H), 3.34–3.30 (m, 4H), 3.23 (q, $J$ = 8.9 Hz, 1H), 3.06 (t, $J$ = 13.1 Hz, 1H), 3.01–2.95 (m, 1H), 2.44 (d, $J$ = 11.0 Hz, 1H) 2.40–2.32 (m, 2H), 2.14–2.08 (m, 6H), 2.01–1.94 (m, 3H), 1.90–1.84 (m, 2H), 1.79–1.74 (m, 3H), 1.66 (s, 3H), 1.65 (s, 3H), 1.61–1.52 (m, 5H), 1.39–1.29 (m, 5H), 1.16–1.06 (m, 3H), 0.97 (d, $J$ = 7.2 Hz, 6H), 0.90–0.84 (m, 7H); HRMS (ESI) $m/z$ 873.5221 [(M + H)$^+$ calcd for $C_{46}H_{72}N_4O_{12}$ 873.5220].

**JH-FK-31.** Following *General Procedure A*, FK520 (40.0 mg, 0.051 mmol) was reacted with 2-hydrazinyl-2-oxoacetamide (31.5 mg, 0.306 mmol) and TFA (0.4 µL, 0.005 mmol) in MeOH (0.5 mL). The residue was purified via column chromatography (hexanes/acetone, 1/1) to yield JH-FK-31 (22.5 mg, 50%) as a white solid: $^1$H NMR (500 MHz, acetone-$d_6$) δ 11.77 (s, 1H), 7.49 (s, 1H), 6.89 (s, 1H), 5.47 (s, 1H), 5.25 (s, 1H), 5.17 (d, $J$ = 10.5 Hz, 1H), 5.12 (d, $J$ = 7.8 Hz, 1H), 4.35 (s, 1H), 4.33 (d, $J$ = 13.9 Hz, 1H), 4.22 (d, $J$ = 4.1 Hz, 1H), 4.08 (s, 1H), 3.44 (dd, $J$ = 15.4, 10.1

Hz, 2H), 3.36 (s, 3H), 3.35 (s, 3H), 3.32 (s, 3H), 3.00–2.89 (m, 3H), 2.42 (d, *J* = 11.6 Hz, 1H), 2.38–2.32 (m, 1H), 2.28–2.17 (m, 3H), 2.15–2.10 (m, 3H), 2.03–1.98 (m, 1H), 1.96 (s, 1H), 1.93–1.84 (m, 2H), 1.81–1.71 (m, 3H), 1.67 (s, 3H), 1.66 (s, 3H), 1.63–1.52 (m, 5H), 1.38–1.29 (m, 4H), 1.15–1.04 (m, 2H), 1.00 (t, *J* = 6.5 Hz, 7H), 0.95 (d, *J* = 11.1 Hz, 1H), 0.89 (t, *J* = 7.3 Hz, 4H), 0.83 (d, *J* = 6.6 Hz, 3H); HRMS (ESI) *m/z* 877.5169 $[(M + H)^+$ calcd for $C_{44}H_{72}N_4O_{13}$ 877.5169].

**JH-FK-32.** To a solution of FK520 (40.0 mg, 0.051 mmol) in EtOH (0.5 mL) was added 4*H*-1,2,4-triazol-4-amine (25.7 mg, 0.306 mmol) and TFA (1 µL, 0.013 mmol)) in a sealed vial. The reaction mixture was stirred at 90 °C for 48 hr. After cooling to room temperature, the reaction vessel was unsealed and concentrated *in vacuo*. The crude mixture was reconstituted in EtOAc and washed with $H_2O$ and brine. The organic layer was dried over anhydrous $Na_2SO_4$ and condensed *in vacuo*. The residue was purified by column chromatography (silica gel, hexanes/acetone, 2/1) to afford JH-FK-32 (8.1 mg, 19%) as a white solid**:** $^1$H NMR (500 MHz, acetone-*d₆*) δ 8.54 (s, 2H), 5.30 (d, *J* = 9.6 Hz, 1H), 5.26 (s, 1H), 5.17–5.10 (m, 2H), 4.73–4.69 (m, 1H), 4.62 (d, *J* = 3.8 Hz, 1H), 4.04 (s, 1H), 3.63 (s, 1H), 3.62 (d, *J* = 6.9 Hz, 2H), 3.54 (dt, *J* = 9.3, 6.0 Hz, 1H), 3.45–3.40 (m, 1H), 3.37 (s, 3H), 3.35 (s, 3H), 3.34 (s, 3H), 3.32–3.27 (m, 2H), 3.03–2.93 (m, 2H), 2.41–2.38 (m, 2H), 2.37–2.27 (m, 2H), 2.23–2.17 (m, 2H), 2.13–2.09 (m, 2H), 1.99–1.85 (m, 6H), 1.70 (s, 3H), 1.65 (s, 3H), 1.62–1.48 (m, 4H), 1.40–1.28 (m, 7H), 1.16–1.04 (m, 3H), 0.99–0.94 (m, 7H), 0.93–0.87 (m, 3H); HRMS (ESI) *m/z* 858.5222 $[(M + H)^+$ calcd for $C_{45}H_{71}N_5O_{11}$ 858.5223].

**JH-FK-33**. Following *General Procedure A*, FK520 (40.0 mg, 0.051 mmol) was reacted with thiosemicarbazide (27.9 mg, 0.306 mmol) and TFA (0.4 µL, 0.005 mmol) in MeOH (0.5 mL). The residue was purified via column chromatography (hexanes/acetone, 2/1 to 1/1) to yield JH-FK-33 (34.8 mg, 79%) as a white solid: $^1$H NMR (500 MHz, acetone-*d₆*) δ 10.22 (s, 1H), 7.52 (s, 1H), 7.20 (s, 1H), 5.34 (d, *J* = 5.0 Hz, 1H), 5.21 (d, *J* = 9.2 Hz, 1H), 5.17 (d, *J* = 10.7 Hz, 1H), 4.92 (s, 1H), 4.62 (s, 1H), 4.47 (t, *J* = 3.8 Hz, 1H), 4.34 (d, *J* = 15.6 Hz, 1H), 4.06–4.00 (m, 1H), 3.63 (d, *J* = 3.1 Hz, 1H), 3.59 (d, *J* = 9.6 Hz, 1H) 3.53 (d, *J* = 10.7 Hz, 1H), 3.43 (td, *J* = 11, 4.4 Hz, 1H), 3.37 (s, 3H), 3.56 (s, 3H), 3.34–3.32 (m, 3H), 3.30 (s, 1H), 3.24 (dd, *J* = 15.4, 9.2 Hz, 1H), 3.10 (t, *J* = 13.0 Hz, 1H), 3.01–2.95 (s, 1H), 2.60 (s, 1H), 2.45 (td, *J* = 14.2, 4.1 Hz, 1H), 2.39–2.31 (m, 2H), 2.25–2.19 (m, 1H), 2.18–2.11 (m, 4H), 2.02–1.95 (m, 3H), 1.90–1.85 (m, 2H), 1.77 (d, *J*

= 8.5 Hz, 2H) 1.71 (s, 1H), 1.66 (s, 3H), 1.63 (s, 3H), 1.60–1.46 (m, 2H), 1.40–1.27 (m, 5H), 1.20 (s, 3H), 1.14–1.04 (m, 3H), 0.97–0.93 (m, 4H), 0.88–0.85 (m, 4H); HRMS (ESI) *m*/*z* 865.4991 [(M + H)$^+$ calcd for $C_{44}H_{72}N_4O_{11}S$ 865.4989].

**JH-FK-34.** Following *General Procedure A*, FK520 (40.0 mg, 0.051 mmol) was reacted with 2-methoxyacetohdyrazide (31.9 mg, 0.306 mmol) and TFA (0.4 µL, 0.005 mmol) in MeOH (0.5 mL). The residue was purified via column chromatography (hexanes/acetone, 3/1 to 1/1) to yield JH-FK-34 (42.2 mg, 94%) as a white solid: $^1$H NMR (500 MHz, acetone-*d$_6$*) δ 11.02 (s, 1H), 5.51 (s, 1H), 5.29 (s, 1H), 5.16 (d, *J* = 9.0 Hz, 1H), 5.11 (d, *J* = 9.5 Hz, 1H), 4.50 (s, 1H), 4.33 (d, *J* = 14.5 Hz, 1H), 4.22 (d, *J* = 6.0 Hz, 1H), 4.06 (d, *J* = 10.7 Hz, 1H), 3.96 (q, *J* = 17.4 Hz, 2H), 3.66 (s, 1H), 3.47 (S, 2H), 3.36 (s, 6H), 3.33 (s, 4H), 3.31–3.27 (m, 2H), 3.01–2.95 (m, 1H), 2.41–2.31 (m, 2H), 2.27 (m, 4H), 2.03–1.95 (m, 2H), 1.92–1.84 (m, 3H), 1.82–1.72 (m, 3H), 1.68 (s, 3H), 1.66 (s, 3H), 1.63–1.50 (m, 7H), 1.40–1.28 (m, 5H), 1.16–1.04 (m, 2H), 1.01 (t, *J* = 6.9 Hz, 5H), 0.98–0.91 (m, 3H), 0.88 (t, *J* = 7.2 Hz, 4H), 0.83 (d, *J* = 6.6 Hz, 2H); HRMS (ESI) *m*/*z* 878.5374 [(M + H)$^+$ calcd for $C_{46}H_{75}N_3O_{13}$ 878.5373].

**JH-FK-35.** Following *General Procedure B*, FK520 (50.0 mg, 0.051 mmol) was reacted with hydroxylamine hydrochloride (5.3 mg, 0.076 mmol) in EtOH (0.3 mL) to yield JH-FK-35 (19.8 mg, 39%) as a white solid: $^1$H NMR (500 MHz, acetone-*d$_6$*) δ 9.73 (s, 1H), 5.23 (s, 1H), 5.02 (d, *J* = 3.1 Hz, 1H), 4.94 (d, *J* = 9.2 Hz, 1H), 4.40 (s, 1H), 4.19 (d, *J* = 8.5 Hz, 1H), 4.11 (s, 1H), 4.05 (dt, *J* = 9.2, 5.8 Hz, 1H), 3.69 (d, *J* = 6.7 Hz, 1H), 3.34 (d, *J* = 10.4 Hz, 1H), 3.27 (t, *J* = 10.6 Hz, 1H), 3.24 (s, 3H), 3.23 (s, 3H), 3.21 (s, 3H), 3.19–3.15 (m, 2H), 2.91–2.82 (m, 2H), 2.33 (d, *J* = 14.0 Hz, 1H), 2.26–2.19 (m, 1H), 2.10–2.00 (m, 6H), 1.76–1.60 (m, 7H), 1.53 (s, 3H), 1.48 (s, 3H), 1.45–1.39 (m, 2H), 1.38–1.25 (m, 4H), 1.22–1.13 (m, 4H), 1.02–0.93 (s, 1H), 0.88 (d, *J* = 6.4 Hz, 3H), 0.83–0.80 (m, 4H), 0.78–0.72 (m, 7H); HRMS (ESI) *m*/*z* 807.4984 [(M + H)$^+$ calcd for $C_{43}H_{70}N_2O_{12}$ 807.5002].

**JH-FK-36.** Following *General Procedure A*, FK520 (40.0 mg, 0.051 mmol) was reacted with Girard's reagent T ((carboxymethyl)trimethylammonium chloride hydrazide) (51.3 mg, 0.306 mmol) and TFA (0.4 µL, 0.005 mmol) in MeOH (0.5 mL). The residue was purified via column chromatography ($CH_2Cl_2$/MeOH, 5/1) to yield JH-FK-36 (41.3 mg, 86%) as a white solid: $^1$H

NMR (500 MHz, acetone-$d_6$) δ 11.72 (s, 1H), 5.35 (s, 1H), 5.19 (d, *J* = 8.9 Hz, 1H), 5.16 (d, *J* = 9.5 Hz, 1H), 5.12 (d, *J* = 9.5 Hz, 1H), 4.82 (q, *J* = 17.0 Hz, 1H), 4.73 (s, 2H), 4.53 (s, 1H), 4.34 (d, *J* = 18.3 Hz, 2H), 3.82 (s, 1H), 3.65 (s, 1H), 3.59 (s, 3H), 3.50 (s, 9H), 3.37 (s, 3H), 3.35 (s, 3H), 3.33 (s, 3H), 3.22–3.14 (m, 2H), 2.43–2.36 (m, 2H), 2.29–2.10 (m, 6H), 2.01–1.93 (m, 4H), 1.91–1.85 (m, 2H), 1.81–1.73 (s, 3H), 1.69 (s, 1H), 1.67 (s, 2H), 1.57 (s, 3H), 1.55–1.47 (m, 2H), 1.41–1.28 (m, 4H), 1.17–1.04 (m, 2H), 0.98 (d, *J* = 6.4 Hz, 1H), 0.95–0.91 (m, 6H), 0.89–0.84 (m, 7H); HRMS (ESI) *m/z* 905.8540 $[(M)^+$ calcd for $C_{48}H_{81}N_4O_{12}$ 905.8546].

**JH-FK-37** and **JH-FK-38.** Following *General Procedure B*, FK520 (50.0 mg, 0.051 mmol) was reacted with hydroxylamine hydrochloride (5.3 mg, 0.076 mmol) in EtOH (0.3 mL) to yield JH-FK-37 (25.2 mg, 49%) and JH-FK-38 (11.7 mg, 22%) as white solids: For JH-FK-37: $^1$H NMR (500 MHz, acetone-$d_6$) δ 5.37 (s, 1H), 5.15 (d, *J* = 11.0 Hz, 1H), 5.08 (d, *J* = 9.2 Hz, 1H), 4.60 (d, *J* = 8.4 Hz, 1H), 4.34 (d, *J* = 13.6 Hz, 1H), 4.29 (s, 1H), 3.85 (s, 3H), 3.62 (d, *J* = 2.8 Hz, 1H), 3.50 (d, *J* = 10.4 Hz, 1H), 3.45–3.40 (m, 2H), 3.38 (s, 2H), 3.36 (s, 3H), 3.35 (s, 2H), 3.32 (s, 1H), 3.04 (td, *J* = 13.3, 3.4 Hz, 1H), 2.98 (ddd, *J* = 11.2, 8.9, 4.2 Hz, 1H), 2.46 (d, *J* = 14.0 Hz, 1H), 2.40–2.32 (m, 1H), 2.24–2.12 (m, 5H), 2.03–1.98 (m, 2H), 1.96 (s, 2H), 1.89–1.72 (m, 7H), 1.67 (s, 3H), 1.64–1.61 (m, 2H), 1.58 (s, 3H), 1.53–1.29 (m, 8H), 1.16–1.04 (m, 2H), 1.00 (d, *J* = 6.6 Hz, 3H), 0.96 (d, *J* = 7.4 Hz, 3H), 0.91 (d, *J* = 6.4 Hz, 3H), 0.86 (t, *J* = 7.4 Hz, 4H); HRMS (ESI) *m/z* 821.5155 $[(M + H)^+$ calcd for $C_{44}H_{72}N_2O_{12}$ 821.5158]; For JH-FK-38: $^1$H NMR (500 MHz, acetone-d6) δ 5.25 (d, *J* = 6.9 Hz, 1H), 5.23 (d, *J* = 8.7 Hz, 1H), 5.10 (d, *J* = 9.9 Hz, 1H), 4.79 (s, 1H), 4.60 (s, 1H), 4.34 (d, *J* = 11.8 Hz, 1H), 4.01–3.96 (m, 1H), 3.79 (s, 3H), 3.69–3.59 (m, 4H), 3.47–3.39 (m, 1H), 3.38 (s, 3H), 3.35 (s, 3H), 3.33 (s, 3H), 3.33–3.27 (m, 2H), 3.15–3.02 (m, 2H), 3.00–2.94 (m, 1H), 2.65–2.59 (m, 1H), 2.39–2.29 (m, 1H), 2.33–2.10 (m, 5H), 2.03–1.95 (m, 4H), 1.90–1.69 (m, 6H), 1.65 (s, 3H), 1.59 (s, 3H), 1.46–1.25 (m, 8H), 1.16–1.04 (m, 2H), 0.92–-0.89 (m, 6H), 0.86 (t, *J* = 7.4 Hz, 4H); HRMS (ESI) *m/z* 843.4980 $[(M + Na)^+$ calcd for $C_{44}H_{72}N_2O_{12}$ 843.4987].

**JH-FK-40.** Following *General Procedure A*, FK520 (40.0 mg, 0.051 mmol) was reacted with isopropylhydrazine hydrochloride (33.8 mg, 0.306 mmol) and TFA (0.4 µL, 0.005 mmol) in MeOH (0.5 mL). The residue was purified via column chromatography (hexanes/acetone, 2/1) to yield JH-FK-40 (7.2 mg, 17%) as a white solid: $^1$H NMR (500 MHz, acetone-$d_6$) δ 6.97 (s, 1H), 5.37 (s,

1H), 5.13 (d, $J$ = 8.6 Hz, 1H), 5.03 (d, $J$ = 9.0 Hz, 1H), 4.71 (s, 1H), 4.33 (d, $J$ = 4.6 Hz, 1H), 4.29 (d, $J$ = 14.3 Hz, 1H), 3.99 (s, 1H), 3.62 (d, $J$ = 2.9 Hz, 1H), 3.36 (s, 4H), 3.35 (s, 3H), 3.33 (s, 3H), 3.31–3.25 (m, 3H), 3.09 (q, $J$ = 8.3 Hz, 1H), 3.00–2.90 (m, 2H), 2.60 (s, 1H), 2.40–2.32 (m, 2H), 2.26–2.16 (m, 3H), 2.14 (s, 2H), 2.01–1.96 (m, 2H), 1.89–1.71 (m, 6H), 1.65 (s, 3H), 1.60 (s, 3H). 1.58–1.41 (m, 5H), 1.36–1.26 (m, 6H), 1.18 (d, $J$ = 6.4 Hz, 3H), 1.13 (d, $J$ = 6.3 Hz, 3H), 1.03 (d, $J$ = 6.4 Hz, 3H), 0.96 (d, $J$ = 7.2 Hz, 3H), 0.88 (t, $J$ = 7.4 Hz, 4H), 0.83 (d, $J$ = 6.6 Hz, 3H); HRMS (ESI) *m/z* 848.5617 $[(M + H)^+$ calcd for $C_{46}H_{77}N_3O_{11}$ 848.5631].

**JH-FK-43.** Following *General Procedure A*, FK520 (40.0 mg, 0.051 mmol) was reacted with 2,2,2-trifluoroethylhydrazine (70% wt. in $H_2O$) (49.9 mg, 0.306 mmol) and TFA (0.4 μL, 0.005 mmol) in MeOH (0.5 mL). The residue was purified via column chromatography (hexanes/acetone, 3/1 to 1/1) to yield JH-FK-43 (28.2 mg, 62%) as a white solid: $^1$H NMR (500 MHz, acetone-$d_6$) δ 7.20 (d, $J$ = 6.9 Hz, 1H), 5.82 (s, 1H), 5.39 (s, 1H), 5.13 (d, $J$ = 8.9 Hz, 1H), 5.05 (d, $J$ = 9.2 Hz, 1H), 4.84 (s, 1H), 4.33 (s, 1H), 4.31 (d, $J$ = 9.5 Hz, 1H), 4.05–4.00 (m, 1H), 3.80–3.60 (m, 3H), 3.42 (d, $J$ = 8.4 Hz, 1H), 3.36 (s, 6H), 3.34 (s, 3H), 3.33–3.27 (m, 2H), 3.07 (q, $J$ = 7.9 Hz, 1H), 3.00–2.94 (m, 2H), 2.60 (s, 1H), 2.40–2.16 (m, 5H), 2.14 (s, 2H), 2.12–2.06 (m, 4H), 2.03–1.99 (m, 1H), 1.89–1.74 (m, 4H), 1.65 (s, 3H), 1.60 (s, 3H),1.58–1.53 (m, 2H), 1.50–1.27 (m, 8H), 1.08 (qd, $J$ = 13.4, 3.7 Hz, 1H), 1.02 (d, $J$ = 6.4 Hz, 2H), 0.95 (d, $J$ = 7.1 Hz, 3H), 0.87 (t, $J$ = 7.3 Hz, 4H), 0.83 (d, $J$ = 6.7 Hz, 2H); HRMS (ESI) *m/z* 888.5185 $[(M + H)^+$ calcd for $C_{44}H_{72}F_3N_3O_{11}$ 888.5192].

**JH-FK-45.** Following *General Procedure A*, FK520 (80.0 mg, 0.102 mmol) was reacted with methylhydrazine (28.2 mg, 0.612 mmol) and TFA (0.8 μL, 0.01 mmol) in MeOH (1.0 mL). The residue was purified via column chromatography (hexanes/acetone, 3/1 to 1/1) to yield JH-FK-45 (7.8 mg, 9%) as a white solid: $^1$H NMR (500 MHz, acetone-$d_6$) δ 5.39 (d, $J$ = 7.7 Hz, 1H), 5.30 (d, $J$ = 4.7 Hz, 1H), 5.20 (d, $J$ = 10.4 Hz, 1H), 4.79 (d, $J$ = 9.8 Hz, 1H), 4.07 (s, 1H), 3.93 (s, 1H), 3.64–3.58 (m, 2H), 3.46–3.40 (m, 3H), 3.35 (s, 4H), 3.26–3.23 (m, 7H), 3.20–3.15 (m, 1H), 3.00–2.90 (m, 3H), 2.64–2.59 (m, 1H), 2.40–2.25 (m, 4H), 2.03–1.95 (m, 2H), 1.93–1.85 (m, 3H), 1.84 (s, 3H), 1.74–1.66 (m, 4H), 1.64–1.53 (m, 8H), 1.39–1.28 (m, 6H), 1.26–1.06 (m, 5H), 0.99–0.92 (m, 2H), 0.90–0.85 (m, 6H), 0.81 (t, $J$ = 7.5 Hz, 3H); HRMS (ESI) *m/z* 820.5315 $[(M + H)^+$ calcd for $C_{44}H_{73}N_3O_{11}$ 820.5318].

**JH-FK-46**. Following *General Procedure A,* FK520 (80.0 mg, 0.102 mmol) was reacted with 1,1-dimethylhydrazine (36.8 mg, 0.612 mmol) and TFA (0.8 μL, 0.01 mmol) in MeOH (1.0 mL) The residue was purified via column chromatography (hexanes/acetone, 3/1 to 1/1) to yield JH-FK-46 (6.5 mg, 8%) as a white solid: $^{1}$H NMR (500 MHz, acetone-*d$_6$*) δ 5.47 (s, 1H), 5.18 (d, *J* = 9.2 Hz, 1H), 5.06 (d, *J* = 8.6 Hz, 1H), 4.71 (s, 1H), 4.61 (s, 1H), 4.36 (d, *J* = 11.9 Hz, 1H), 3.72 (dt, *J* = 8.7 Hz, 4.0 Hz, 1H), 3.64–3.60 (m, 3H), 3.48–3.40 (m, 2H), 3.38 (s, 3H), 3.35 (s, 3H), 3.34 (s, 3H), 3.32–3.27 (m, 2H), 3.15 (q, *J* = 7.9 Hz, 1H), 3.02–2.95 (m, 2H), 2.40 (s, 6H), 2.37 (s, 1H), 2.30–2.25 (m, 1H), 2.22–2.14 (m, 5H), 1.99–1.85 (m, 4H), 1.82–1.72 (m, 2H), 1.67 (s, 3H), 1.61 (s, 3H),1.59–1.48 (m, 4H), 1.42–1.24 (m, 6H), 1.17–1.10 (m, 2H), 0.94 (d, *J* = 7.0 Hz, 4H), 0.92 (dd, *J* = 6.4, 2.3 Hz, 4H), 0.88–0.83 (m, 4H); HRMS (ESI) *m*/*z* 834.5468 [$(M + H)^+$ calcd for $C_{45}H_{75}N_3O_{11}$ 834.5474].

**Bis-TBS Ether 8.** Compound **8** was synthesized according to a modified procedure from Simmons *et al.*[1] substituting FK506 with FK520. To a solution of FK520 (504 mg, 0.636 mmol) and 2,6-lutidine (0.370 mL, 341 mg, 3.18 mmol) in $CH_2Cl_2$ (13.0 mL) was added *tert*-butyldimethylsilyl triflate (0.511 mL, 588 mg, 2.22 mmol). The reaction mixture was stirred at 0 °C for 45 min and additional *tert*-butyldimethylsilyl triflate (0.146 mL, 168 mg, 0.636 mmol) added. The reaction was then stirred at 0 °C for 15 min. The reaction mixture was diluted by the addition of EtOAc and washed with sat. aq. $NaHCO_3$, $H_2O$, and brine. The organic phase was dried by addition of anhydrous $Na_2SO_4$ and concentrated *in vacuo*. The residue was purified by column chromatography (silica gel, hexanes/EtOAc, 4/1) to afford **8** (685 mg, 105%) as a colorless oil: $^{1}$H NMR (500 MHz, acetone-*d$_6$*) δ 5.30–5.22 (m, 2H), 5.08 (s, 1H), 4.91 (d, *J* = 10.1 Hz, 1H), 4.63 (s, 1H), 4.37 (d, *J* = 11.5 Hz, 1H), 4.15 (s, 1H), 3.75 (d, *J* = 9.7 Hz, 1H), 3.64 (dd, *J* = 11.1, 5.8 Hz, 1H), 3.61–3.54 (m, 1H), 3.51–3.40 (m, 3H), 3.38 (s, 3H), 3.35–3.32 (m, 6H), 3.30 (s, 1H), 3.05 (t, *J* = 11.5 Hz, 1H), 2.98–2.87 (m, 2H), 2.41–2.15 (m, 6H), 2.13–2.06 (m, 1H), 2.03–1.98 (m, 2H), 1.96 (s, 3H), 1.94–1.73 (m, 5H), 1.72 (s, 3H), 1.69–1.60 (m, 4H), 1.58 (s, 3H), 1.56–1.29 (m, 6H), 0.96–0.92 (m, 4H), 0.91–0.82 (m, 18H), 0.87–0.80 (m, 2H), 0.10 (s, 3H), 0.08–0.05 (m, 9H); HRMS (ESI) *m*/*z* 1042.6442 [$(M + H)^+$ calcd for $C_{55}H_{97}NO_{12}Si_2$ 1042.6442].

**Mono-TBS 9.** To a solution of **8** (93.1 mg, 0.0912 mmol) in $CH_2Cl_2$/MeOH (1/1 v/v, 3.0 mL) was added *p*-toluenesulfonic acid monohydrate (4.3 mg, 0.028 mmol). The reaction was stirred at 25

°C for 5 h. Volume was increased with EtOAc and washed with 10% $NaHCO_3$ solution, $H_2O$, and brine. The organic phase was dried over anhydrous $Na_2SO_4$ and concentrated *in vacuo*. The residue was purified by column chromatography (silica gel, hexanes/EtOAc: 1/1 to 1/4) to afford **9** as a colorless foam (61.6 mg, 75%): $^1$H NMR (500 MHz, acetone-$d_6$) δ 5.30 (d, *J* = 8.1 Hz, 1H), 5.27 (d, *J* = 6.3 Hz, 1H), 5.10 (s, 1H), 4.90 (d, *J* = 9.5 Hz, 1H), 4.64 (s, 1H), 4.35 (d, *J* = 13.4 Hz, 1H), 4.15 (s, 1H), 3.75 (d, *J* = 9.4 Hz, 1H), 3.64–3.54 (m, 3H), 3.50–3.43 (m, 2H), 3.38 (s, 3H), 3.35–3.32 (m, 7H), 3.30 (s, 1H), 3.31–2.95 (m, 2H), 2.90 (dd, *J* = 14.8, 6.9 Hz, 1H), 2.40–2.11 (m, 7H), 1.96 (s, 3H), 1.93–1.74 (m, 8H), 1.72 (s, 3H), 1.69–1.62 (m, 3H), 1.58 (s, 3H), 1.43–1.29 (m, 6H), 0.95–0.92 (m, 4H), 0.90 (s, 9H), 0.83–0.80 (m, 4H), 0.10 (s, 3H), 0.07 (s, 3H); HRMS (ESI) *m*/*z* 928.5578 [(M + H)$^+$ calcd for $C_{49}H_{83}NO_{12}Si$ 925.5277].

**Methyl Ether 10**: To a solution of **9** (62.5 mg, 0.069 mmol) in $CH_2Cl_2$ was added proton sponge (118 mg, 0.552 mmol) and freshly-dried 4 Å molecular sieves (80 mg). The solution was stirred for 1 h at 25 °C. Trimethyloxonium tetrafluoroborate (81.6 mg, 0.552 mmol) was added to the reaction. The solution was stirred for 18 h at 25 °C. The reaction mixture was diluted with $CH_2Cl_2$ and passed through a celite filter. The filtrate was washed twice with 1 N HCl and 10% $NaHCO_3$ solution, dried over anhydrous $Na_2SO_4$, and condensed *in vacuo*. The residue was purified by column chromatography (silica gel, hexanes/EtOAc: 9/1 to 4/1) to afford **10** as a white solid (48.5 mg, 76%): $^1$H NMR (125 MHz, acetone-$d_6$) δ 5.29–5.22 (m, 2H), 5.09 (s, 1H), 4.90 (d, *J* = 11.5 Hz, 1H), 4.62 (d, *J* = 3.6 Hz, 1H), 4.35 (d, *J* = 14.8 Hz, 1H), 4.15 (s, 1H), 3.75 (d, *J* = 7.9 Hz, 1H), 3.64 (dd, *J* = 11.1, 4.0 Hz, 1H), 3.60–3.54 (m, 1H), 3.50–3.44 (m, 2H), 3.38 (s, 3H), 3.37 (s, 2H), 3.36 (s, 1H), 3.35 (s, 2H), 3.34 (s, 1H), 3.33 (s, 2H), 3.30 (s, 1H), 3.09–3.04 (m, 2H), 3.00–2.95 (m, 1H), 2.90 (dd, *J* = 15.1 Hz, 7.0 Hz, 1H), 2.40–2.29 (m, 2H), 2.26 (dd, *J* = 17.5*,* 5.5 Hz, 1H), 2.20–2.10 (m, 3H), 2.03–1.97 (m, 3H), 1.82–1.73 (m, 6H), 1.72 (s, 3H), 1.68–1.61 (m, 3H), 1.57 (s, 3H), 1.43–1.29 (m, 7H), 0.94–0.92 (m, 3H), 0.91–0.87 (m, 13H), 0.84–0.80 (m, 4H), 0.10 (s, 3H), 0.07 (s, 3H); $^{13}$C NMR (500 MHz, acetone-d6) δ 210.2, 198.1, 170.9, 169.8, 166.1, 140.0, 139.1, 132.7, 124.5, 99.4, 98.3, 84.4, 84.2, 83.8, 83.7, 76.3, 74.6, 74.4, 73.6, 70.7, 57.7, 57.61, 57.59, 57.4, 56.4, 56.0, 55.5, 49.8, 44.7, 41.1, 39.5, 37.4, 35.6, 33.2, 31.4, 28.7, 26.6, 26.4, 26.3, 24.4, 21.7, 20.8, 20.0, 18.62, 18.58, 16.9, 16.5,16.2, 16.0, 14.5, 12.7, 12.0, 11.9, 11.1, 10.8.

**JH-FK-41 (5).** Following *General Procedure C*, compound **10** (23.3 mg, 0.025 mmol) was reacted with HF·pyridine (110 mg, 0.1 mL) in MeCN (1.0 mL). The residue was purified via column chromatography (silica gel, hexanes/EtOAc, 2/1) to yield **5** (9.2 mg, 45%) as a white solid: $^1$H NMR (500 MHz, acetone-$d_6$) δ 7.26 (d, *J* = 4.9 Hz, 1H), 5.21 (d, *J* = 9.2 Hz, 1H), 5.06 (s, 1H), 4.96 (d, *J* = 11.5 Hz, 1H), 4.65 (s, 1H), 4.35 (d, *J* = 13.5 Hz, 1H), 3.99–3.95 (m, 1H), 3.81 (d, *J* = 4.5 Hz, 1H), 3.72 (d, *J* = 9.7 Hz, 1H), 3.63 (dd, *J* = 15.0, 5.4 Hz, 1H), 3.58–3.40 (m, 3H), 3.38 (s, 2H), 3.36 (s, 5H), 3.35 (s, 1H), 3.34 (s, 2H), 3.28 (s, 1H), 3.10–3.04 (m, 1H), 3.00–2.94 (m, 2H), 2.37–2.31 (m, 2H), 2.21–2.14 (m, 4H), 1.96–1.92 (m, 4H), 1.82–1.73 (m, 5H), 1.69 (s, 3H), 1.61 (s, 3H), 1.57–1.54 (m, 1H), 1.44–1.30 (m, 6H), 1.10–0.99 (m, 3H), 0.95–0.90 (m, 9H), 0.87–0.81 (m, 4H); HRMS (ESI) *m*/*z* 828.4858 [(M + Na)$^+$ calcd for $C_{44}H_{71}NO_{12}$ 828.4869].

**Benzyl Ether 11**. To a solution of **9** (50.9 mg, 0.056 mmol) was added benzyl 2,2,2-trichloroacetimidate (28.3 mg, 0.112 mmol) in $CH_2Cl_2$ (1.0 mL). Trifluoromethanesulfonic acid (1.7 mg, 0.0112 mmol) was added, and the solution was stirred for 5 h at 25 °C. The reaction was quenched with saturated aqueous $NaHCO_3$, diluted with EtOAc, and extracted with EtOAc. The combined organic phases were dried over anhydrous $Na_2SO_4$, and condensed *in vacuo*. The residue was purified by column chromatography (silica gel, hexanes/EtOAc) to afford **11** as a white solid (13.1 mg, 24%): $^1$H NMR (500 MHz, acetone-$d_6$) δ 7.37 (d, *J* = 7.5 Hz, 2H), 7.32 (t, *J* = 7.2 Hz, 2H), 7.24 (t, *J* = 7.2 Hz, 1H), 5.30–5.19 (m, 2H), 5.10 (s, 1H), 4.91–4.89 (d, *J* = 10.1 Hz, 1H), 4.67 (s, 2H), 4.63 (br s, 1H), 4.36 (d, *J* = 12.8 Hz, 1H), 4.15 (s, 1H), 3.76 (d, *J* = 9.6 Hz, 1H), 3.64 (dd, *J* = 11.2, 4.1 Hz, 1H), 3.61–3.44 (m, 3H), 3.41 (s, 1H), 3.40 (s, 2H), 3.38 (s, 2H), 3.34 (s, 1H), 3.33 (s, 2H), 3.30 (s, 1H), 3.29–3.16 (m, 2H), 3.04 (t, *J* = 13.0 Hz, 1H), 2.90 (dd, *J* = 14.8, 6.9 Hz, 1H), 2.41–2.24 (m, 3H), 2.21–2.08 (m, 4H), 2.02–1.96 (m, 3H), 1.80–1.73 (m, 6H), 1.71 (s, 3H), 1.69–1.62 (m, 3H), 1.58 (s, 2H), 1.43–1.31 (m, 7H), 0.94 (d, *J* = 6.6 Hz, 3H), 0.91–0.89 (m, 10H), 0.86–0.81 (m, 3H), 0.10 (s, 3H), 0.07 (s, 3H); HRMS (ESI) *m*/*z* 1018.9046 [(M + Na)$^+$ calcd for $C_{56}H_{89}NO_{12}Si$ 1018.6048].

**JH-FK-42 (6).** Following *General Procedure C*, compound **11** (13.1 mg, 0.0131 mmol) was reacted with HF·pyridine (110 mg, 0.1 mL) in MeCN (1.0 mL). The residue was purified via column chromatography (silica gel, hexanes/EtOAc, 2/1) to yield **5** (4.1 mg, 35%) as a white solid: $^1$H NMR (500 MHz, acetone-$d_6$) δ 7.37 (d, *J* = 7.2 Hz, 2H), 7.33 (t, *J* = 7.3 Hz, 2H), 7.25 (t, *J* =

7.2 Hz, 1H), 5.27 (d, *J* = 4.9 Hz, 1H), 5.23 (d, *J* = 8.7 Hz, 1H), 5.06 (s, 1H), 4.96 (d, *J* = 10.4 Hz, 1H), 4.67 (s, 2H), 4.64 (br s, 1H), 4.35 (d, *J* = 13.7 Hz, 1H), 4.00–3.94 (m, 1H), 3.81 (d, *J* = 4.5 Hz, 1H), 3.72 (d, *J* = 8.0 Hz, 1H), 3.65 (dd, *J* = 9.7, 4.0 Hz, 1H), 3.58–3.50 (m, 1H), 3.44 (qd, *J* = 11.2, 4.3 Hz, 2H), 3.41 (s, 3H), 3.38 (s, 2H), 3.33 (s, 2H), 3.28–3.16 (m, 2H), 2.97 (dd, *J* = 13.1, 2.9 Hz, 1H), 2.41–2.31 (m, 2H), 2.22–2.14 (m, 4H), 2.10–2.07 (m, 2H), 2.01–1.90 (m, 2H), 1.83–1.73 (m, 6H), 1.69 (s, 3H), 1.62 (s, 3H), 1.60–1.51 (m, 2H), 1.44–1.31 (m, 6H), 1.04 (q, *J* = 10.9 Hz, 2H), 0.95–0.91 (m, 8H), 0.89–0.81 (m, 5H); HRMS (ESI) *m/z* 904.5172 [$(M + Na)^+$ calcd for $C_{50}H_{75}NO_{12}$ 904.5182].

**Ester 12.** To a solution of **9** (52.5 mg, 0.0579 mmol) in pyridine (0.5 mL) was added DMAP (6.1 mg, 0.050 mmol) and phthalic anhydride (40.9 mg, 0.276 mmol). The solution was stirred for 24 hr at 25 °C. The reaction was quenched by addition of $H_2O$ and extracted with $CH_2Cl_2$. The aqueous phase was acidified with $NaH_2PO_4$ and extracted with $CH_2Cl_2$. The combined organic phases were washed with saturated aqueous $NaHCO_3$, dried over anhydrous $Na_2SO_4$ and condensed *in vacuo*. The residue was purified by column chromatography (silica gel, hexanes/EtOAc: 1/2) to afford **12** as a white solid (26.6 mg, 44%): $^1H$ NMR (500 MHz, acetone-*d*$_6$) δ 7.84 (s, 1H), 7.67–7.62 (m, 3H), 5.33–5.28 (m, 2H), 5.10 (s, 1H), 4.94–4.85 (m, 2H), 4.62 (d, *J* = 6.0 Hz, 1H), 4.36 (d, *J* = 11.6 Hz, 1H), 4.17 (s, 1H), 3.76 (d, *J* = 9.6 Hz, 1H), 3.66–3.55 (m, 2H), 3.50–3.43 (m, 2H), 3.38 (s, 2H), 3.34 (m, 2H), 3.32–3.30 (m, 3H), 3.30 (s, 1H), 3.05 (t, *J* = 13.8 Hz, 1H), 2.52–2.24 (m, 4H), 2.21–2.12 (m, 4H), 1.87–1.74 (m, 6H), 1.71 (s, 3H), 1.70–1.65 (m, 2H), 1.60 (s, 3H), 1.57–1.29 (m, 9H), 0.95–0.90 (m, 19H), 0.87–0.80 (m, 5H), 0.12 (s, 3H), 0.08 (s, 3H); HRMS (ESI) *m/z* 1076.5720 [$(M + Na)^+$ calcd for $C_{57}H_{87}NO_{15}Si$ 1076.5737].

**JH-FK-44 (7).** Following *General Procedure C*, compound **12** (26.6 mg, 0.0252 mmol) was reacted with HF·pyridine (165 mg, 0.15 mL) in MeCN (2.0 mL). The residue was purified via column chromatography (silica gel, hexanes/EtOAc, 2/1) to yield **7** (5.2 mg, 22%) as a white solid: $^1H$ NMR (500 MHz, acetone-*d*$_6$) δ 7.83 (s, 1H), 7.69 (s, 1H), 7.64 (d, *J* = 3.4 Hz, 1H), 7.63 (d, *J* = 3.6 Hz, 1H), 5.34–5.25 (m, 2H), 5.06 (s, 1H), 4.98 (d, *J* = 10.1 Hz, 1H), 4.86 (td, *J* = 11.3, 4.9 Hz, 1H), 4.66 (s, 1H), 4.36 (d, *J* = 13.2 Hz, 1H), 3.99 (s, 1H), 3.85 (s, 1H), 3.72 (d, *J* = 9.6 Hz, 1H), 3.64 (dd, *J* = 12.7 Hz, 1H), 3.60–3.42 (m, 3H), 3.38 (s, 2H), 3.35 (s, 1H), 3.34 (s, 2H), 3.33 (s, 2H), 3.20 (s, 1H), 2.98 (td, *J* = 13.3, 3.5 Hz, 1H), 2.50–2.26 (s, 2H), 2.22–2.15 (m, 4H), 2.10–

2.07 (m, 2H), 1.97–1.92 (m, 1H), 1.85–1.73 (m, 6H), 1.70 (s, 3H), 1.64 (s, 2H), 1.61–1.29 (m, 8H), 1.25–1.10 (m, 4H), 0.96–0.90 (m, 10H), 0.87–0.81 (m, 4H); HRMS (ESI) *m*/*z* 962.4879 [(M + Na)$^+$ calcd for $C_{51}H_{73}NO_{15}$ 962.4872].

**4. Table S1.** Synthesis and structure of C22-modified FK520 derivatives.

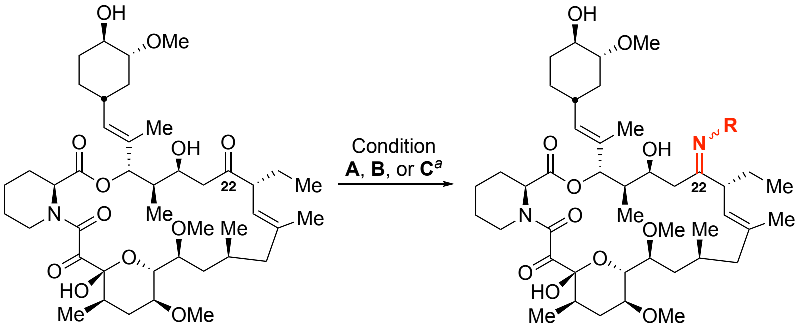


FK520 (**3**)    C22-modified FK520 analogs

| Compound | Reactant | Condition | R in Product | Yield |
|---|---|---|---|---|
| JH-FK-29 | 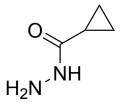 | A | 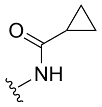 | 76% |
| JH-FK-30 | 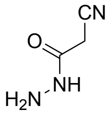 | A | 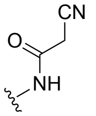 | 84% |
| JH-FK-31 | 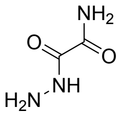 | A | 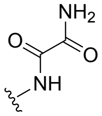 | 50% |
| JH-FK-32 | 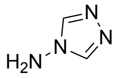 | B | 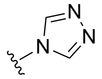 | 19% |
| JH-FK-33 | 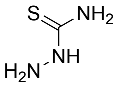 | A | 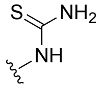 | 79% |
| JH-FK-34 | 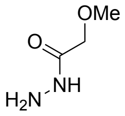 | A | 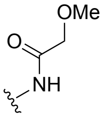 | 94% |
| JH-FK-35 | $NH_2OH\cdot HCl$ | C | OH | 39% |
| JH-FK-36 | 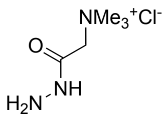 | A | 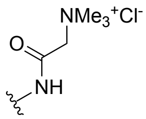 | 86% |
| JH-FK-37[b] | $NH_2OMe\cdot HCl$ | C | (*E*)- or (*Z*)-OMe | 49% |
| JH-FK-38[b] | $NH_2OMe\cdot HCl$ | C | (*E*)- or (*Z*)-OMe | 22% |
| JH-FK-40 | 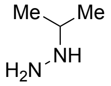 | A | 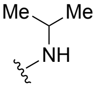 | 17% |
| JH-FK-43 | 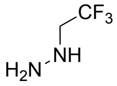 | A | 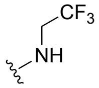 | 62% |
| JH-FK-45 | 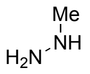 | A | 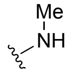 | 9% |

| | | | | |
|---|---|---|---|---|
| JH-FK-46 | 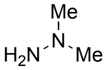 | A | 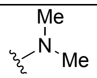 | 8% |

[a] Condition **A**: $RNHNH_2$, TFA, MeOH, 25 °C, 48 h; **B**: 4-amino-4*H*-1,2,4-triazole, TFA, EtOH, 90 °C, 48 h; **C**: $MeOH_2N{\cdot}HCl$, NaOAc, EtOH, 25 °C, 24 h.

[b] Two (*E*/*Z*)-oxime isomers (JH-FK-37 and JH-FK-38) were separated, but the (*E*/*Z*) configuration was not determined.

**5. Supplementary Figure S1.** Immunosuppressive activity, measured as the percentage of IL-2-producing cells, was assessed for JH-FK-29, 30, 31, 32, 33, 34, 35, 36, 37, 38, and 40 across a concentration range of 0.001 to 1,000 nM. FK506 and FK520 are included as positive controls for immunosuppressive activity. All derivatives exhibited lower immunosuppressive *in vitro* compared to FK506 or FK520.

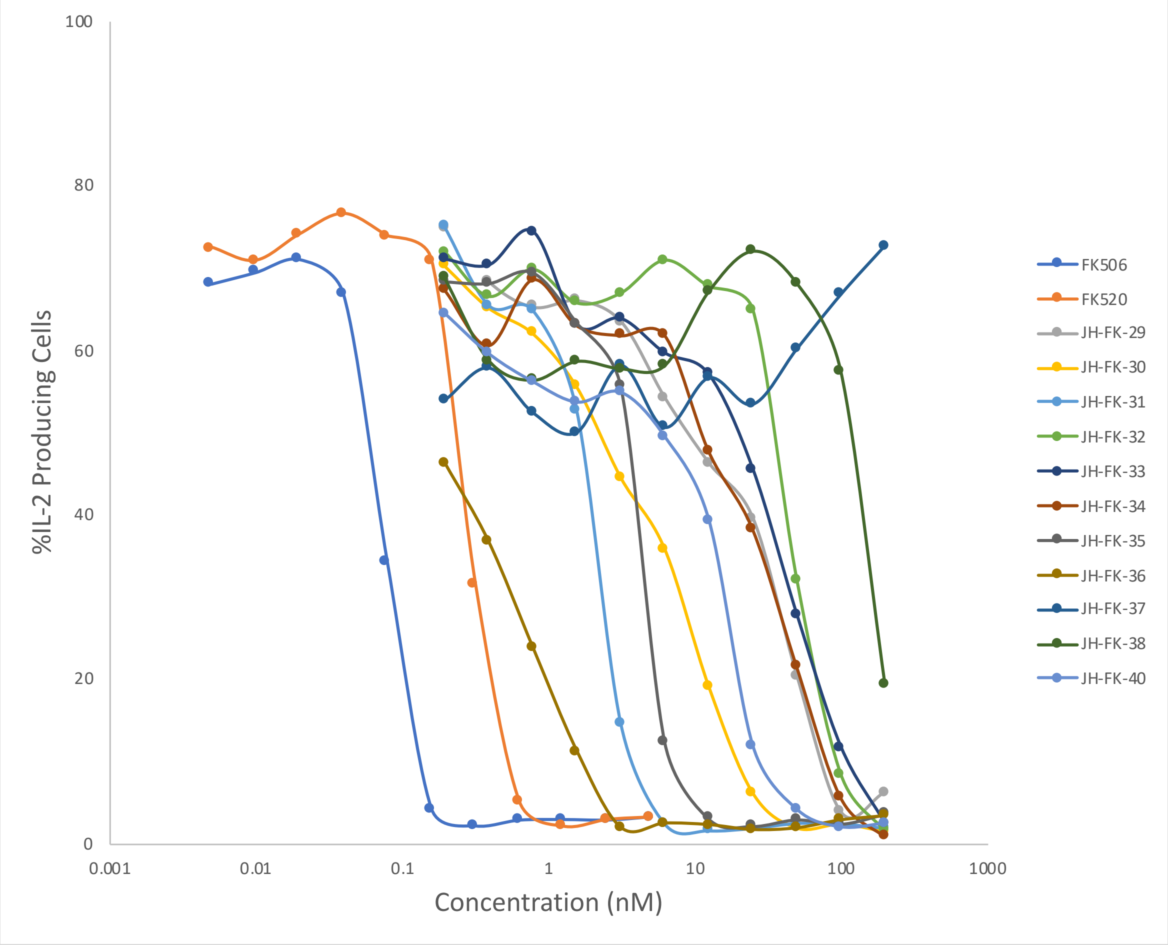

**6. Supplementary Figure S2.** Immunosuppressive activity, measured as the percentage of IL-2-producing cells, was assessed for JH-FK-41, 42, 43, 44, 45, and 46 across a concentration range of 0.001 to 1,000 nM. FK506 and FK520 are included as positive controls for immunosuppressive activity. All derivatives exhibited lower immunosuppressive *in vitro* compared to FK506 or FK520.

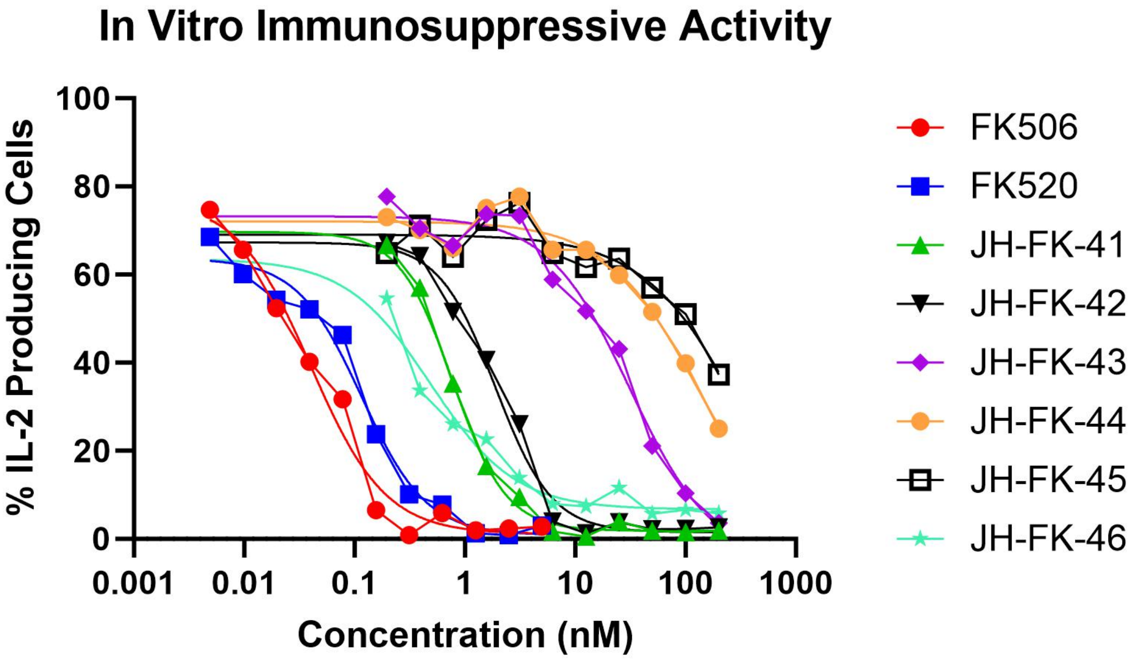

**7. Supplementary Figure S3.** Cytotoxicity data for C22- or C32-modified FK502 derivatives were assessed. The relative cell viability plot of NIH 3T3 cells against FK520 and JH-FK-41 through 46 at concentrations ranging from 0.03 to 300 µM. JH-FK-43 and 44 exhibited greater cytotoxicity than the parent compound FK520, while JH-FK-46 showed comparable cytotoxicity. In contrast, JH-FK-41, JH-FK-42, and JH-FK-45 were less toxic than FK520. Error bars represent one standard deviation.

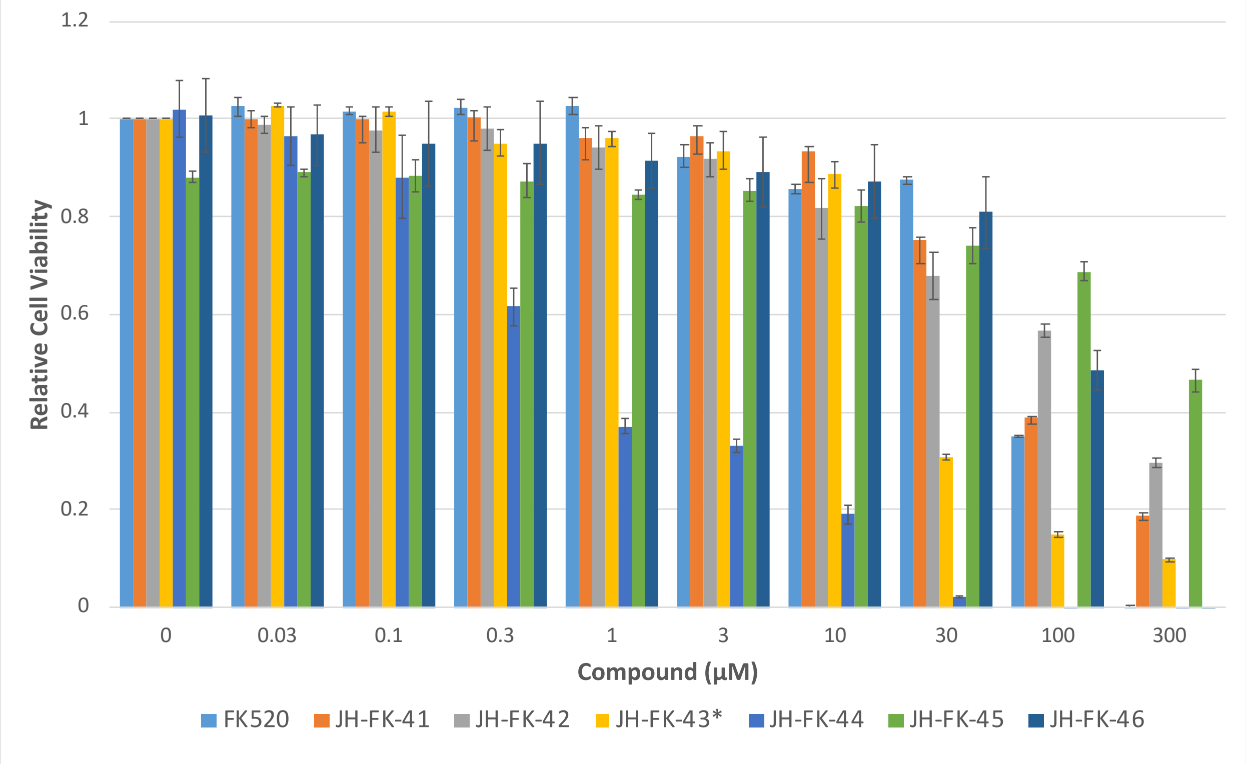


*Precipitation of JH-FK-43 was observed at 100 and 300 µM, contributing to the observed asymptotic behavior in cytotoxicity.

### 8. Copy of $^{1}H$ and $^{13}C$ NMR spectra

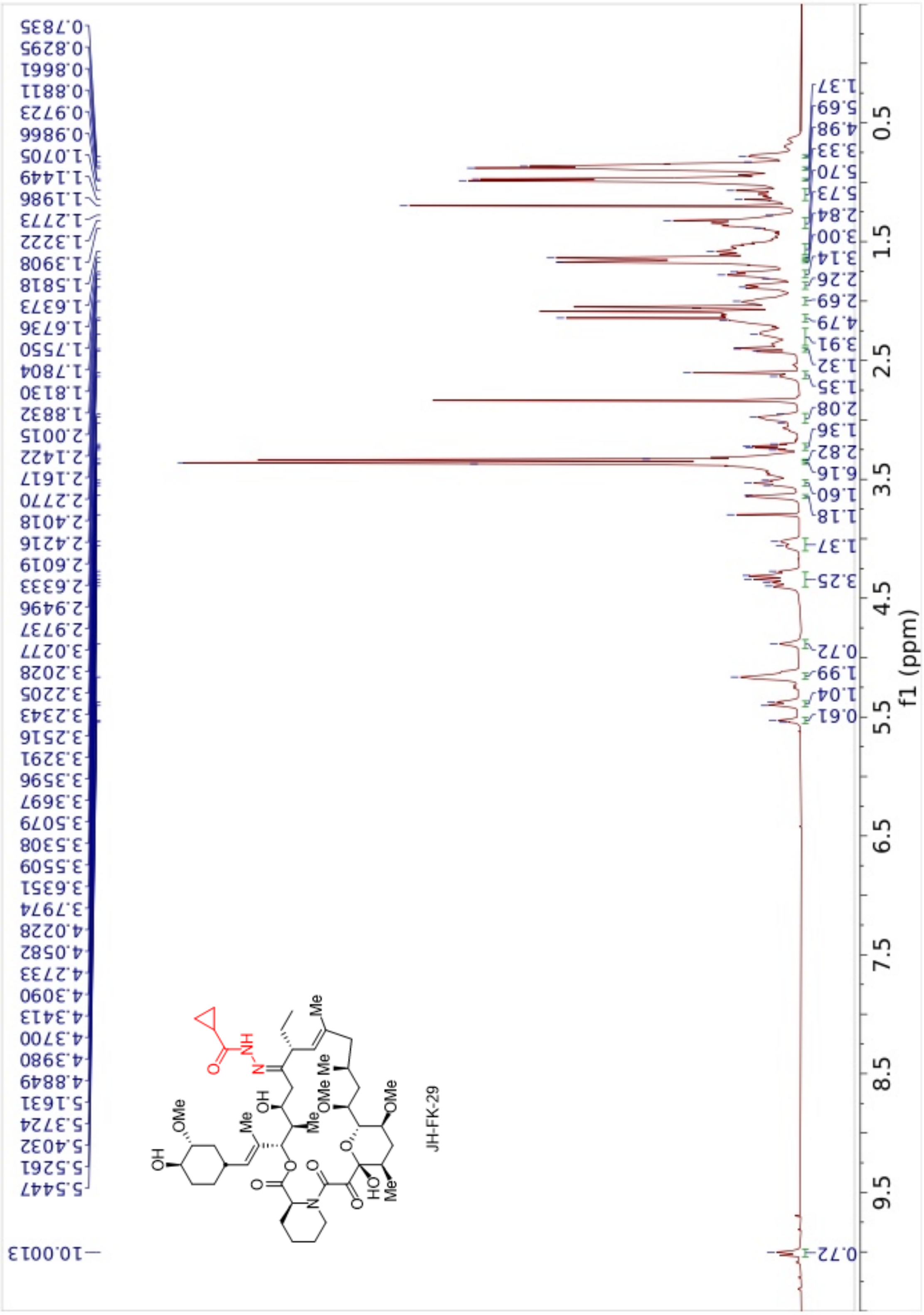

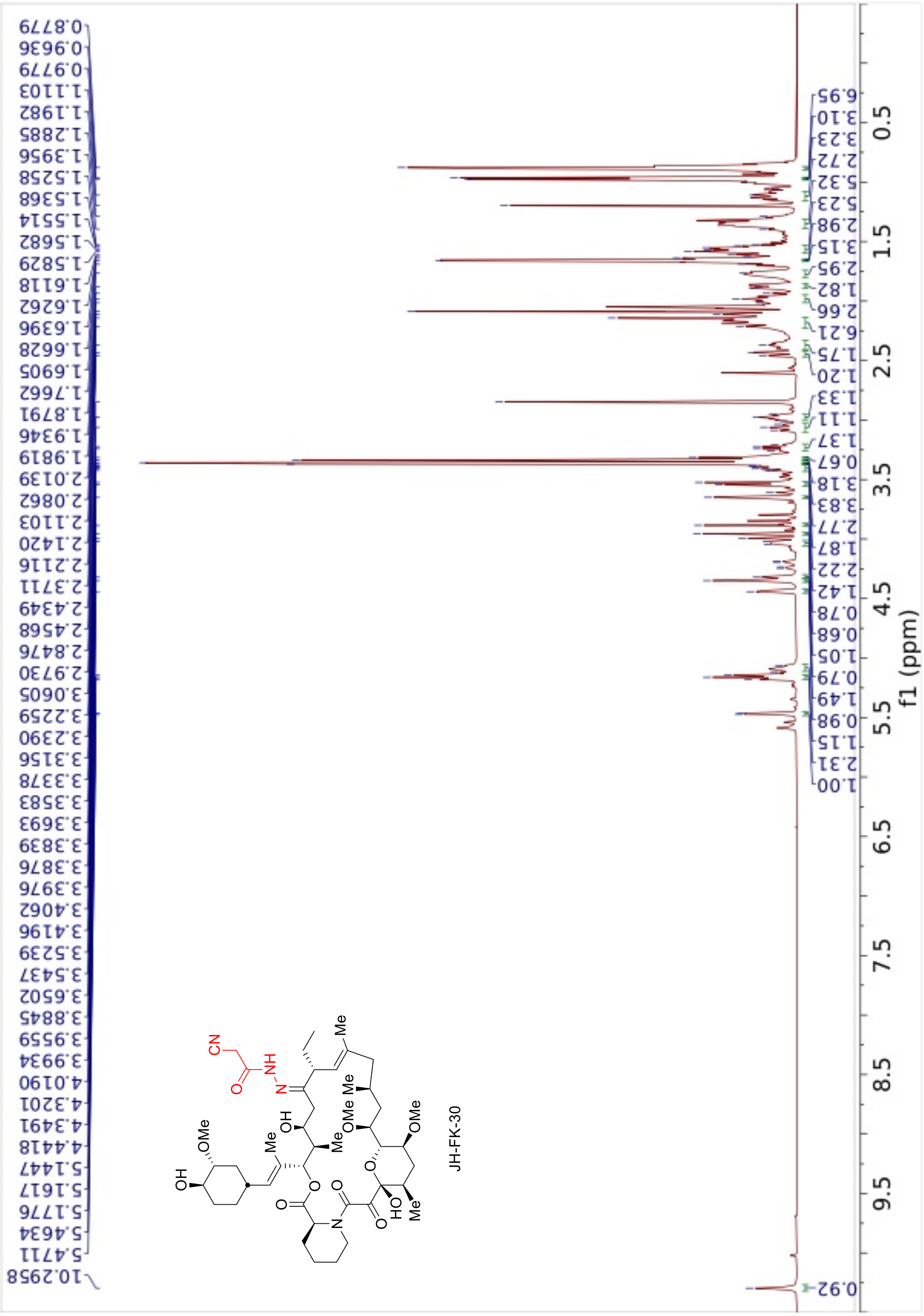
0.8779
0.9636
0.9779
1.1103
1.1982
1.2885
1.3956
1.5258
1.5368
1.5514
1.5682
1.5829
1.6118
1.6262
1.6396
1.6628
1.6905
1.7662
1.8791
1.9346
1.9819
2.0139
2.0862
2.1103
2.1420
2.2116
2.3711
2.4349
2.4568
2.8476
2.9730
3.0605
3.2259
3.2390
3.3156
3.3378
3.3583
3.3693
3.3839
3.3876
3.3976
3.4062
3.4196
3.5239
3.5437
3.6502
3.8845
3.9559
3.9934
4.0190
4.3201
4.3491
4.4418
5.1447
5.1617
5.1776
5.4634
5.4711
10.2958
JH-FK-30
6.95
3.10
3.23
2.72
5.32
5.23
2.98
3.15
2.95
1.82
2.66
6.21
1.75
1.20
1.33
1.11
1.37
0.67
3.18
3.83
2.77
1.87
2.22
1.42
0.78
0.68
1.05
0.79
1.49
0.98
1.15
2.31
1.00
0.92
0.5
1.5
2.5
3.5
4.5
5.5
6.5
7.5
8.5
9.5
f1 (ppm)

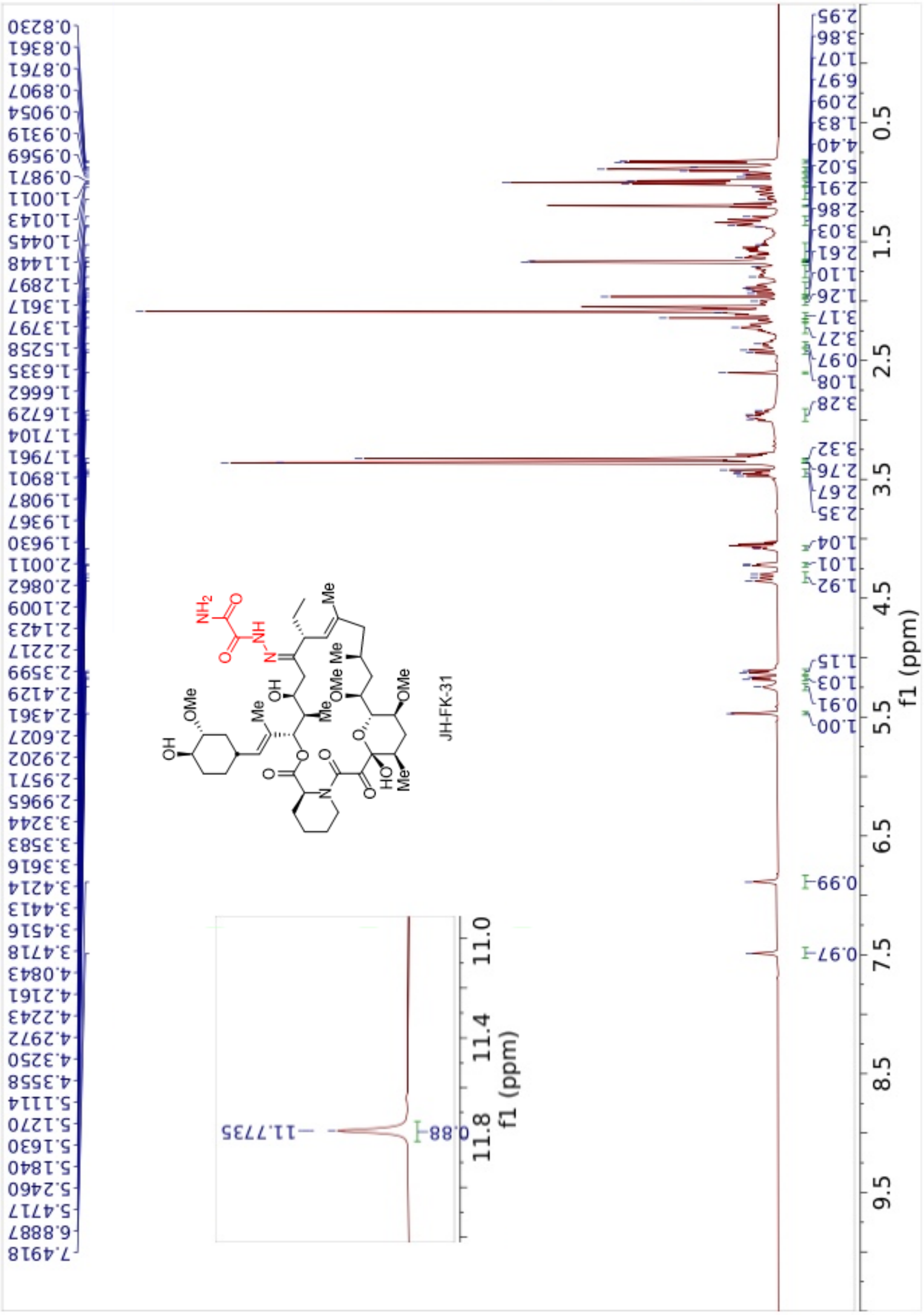
JH-FK-31
f1 (ppm)

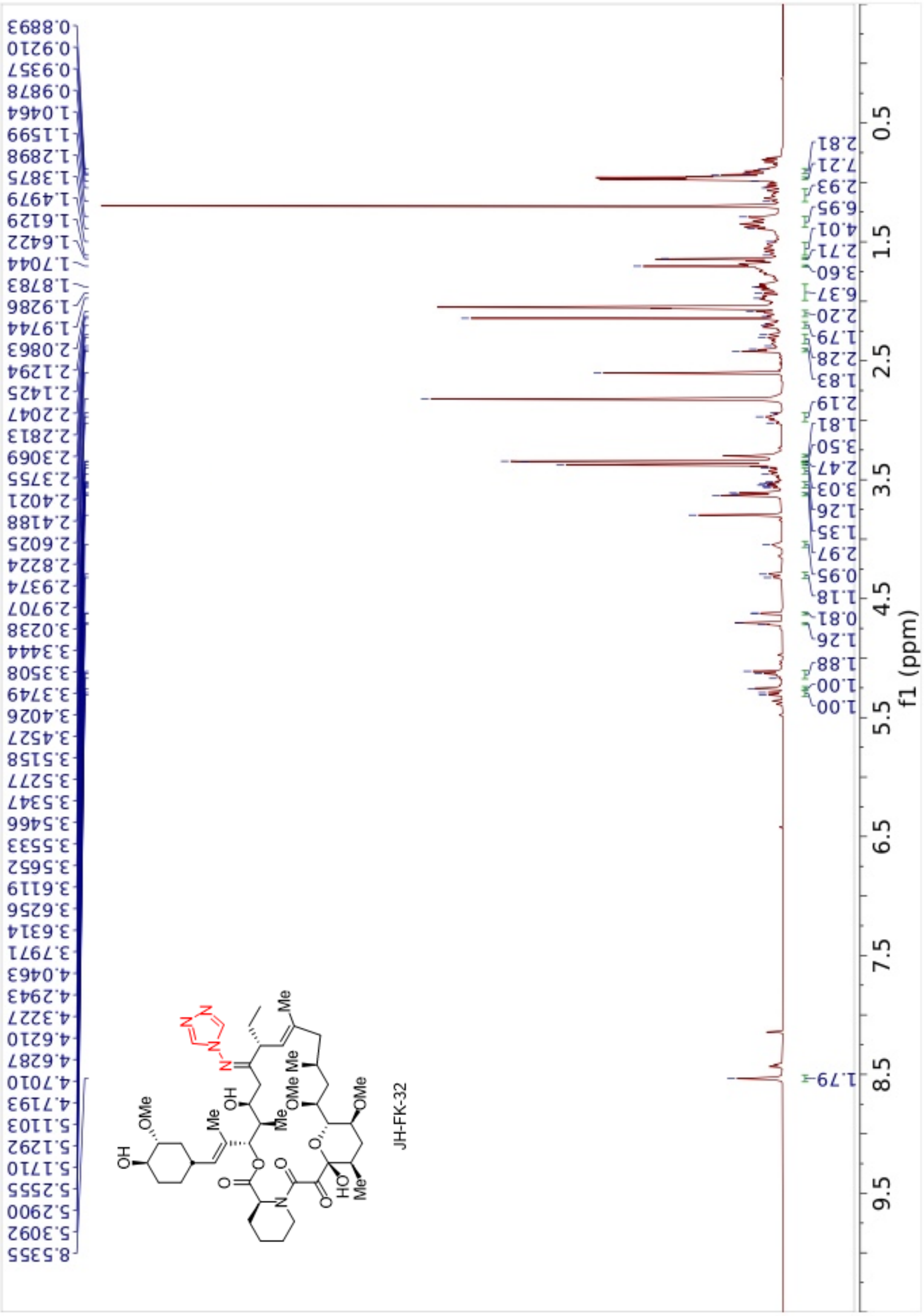
JH-FK-32
f1 (ppm)

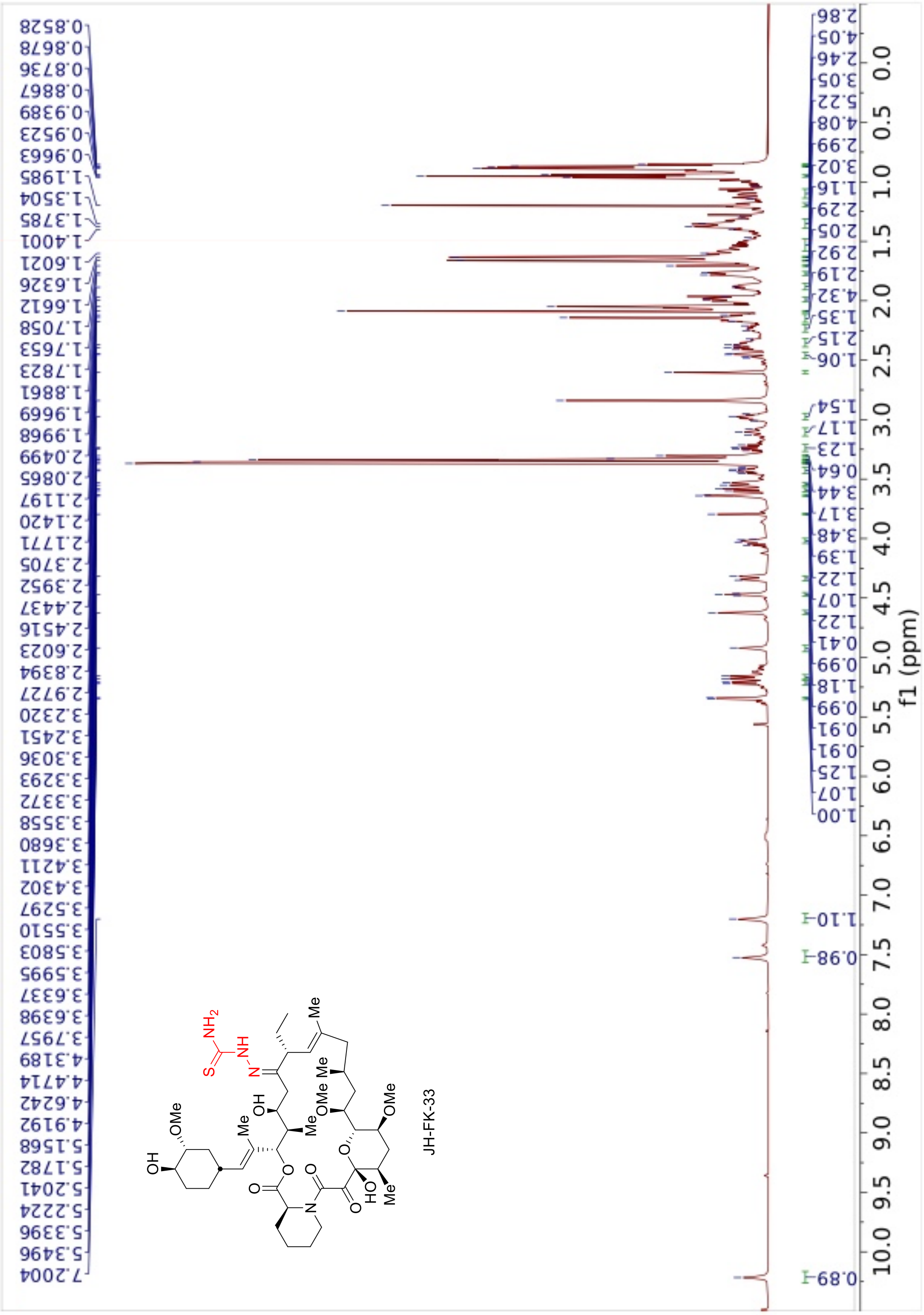
JH-FK-33
f1 (ppm)
0.8528
0.8678
0.8736
0.8867
0.9389
0.9523
0.9663
1.1985
1.3504
1.3785
1.4001
1.6021
1.6326
1.6612
1.7058
1.7653
1.7823
1.8881
1.9669
1.9968
2.0499
2.0865
2.1197
2.1420
2.1771
2.3705
2.3952
2.4437
2.4516
2.6023
2.8394
2.9727
3.2320
3.2451
3.3036
3.3293
3.3372
3.3558
3.3680
3.4211
3.4302
3.5297
3.5510
3.5803
3.5995
3.6337
3.6398
3.7957
4.3189
4.4714
4.6242
4.9192
5.1568
5.1782
5.2041
5.2224
5.3396
5.3496
7.2004

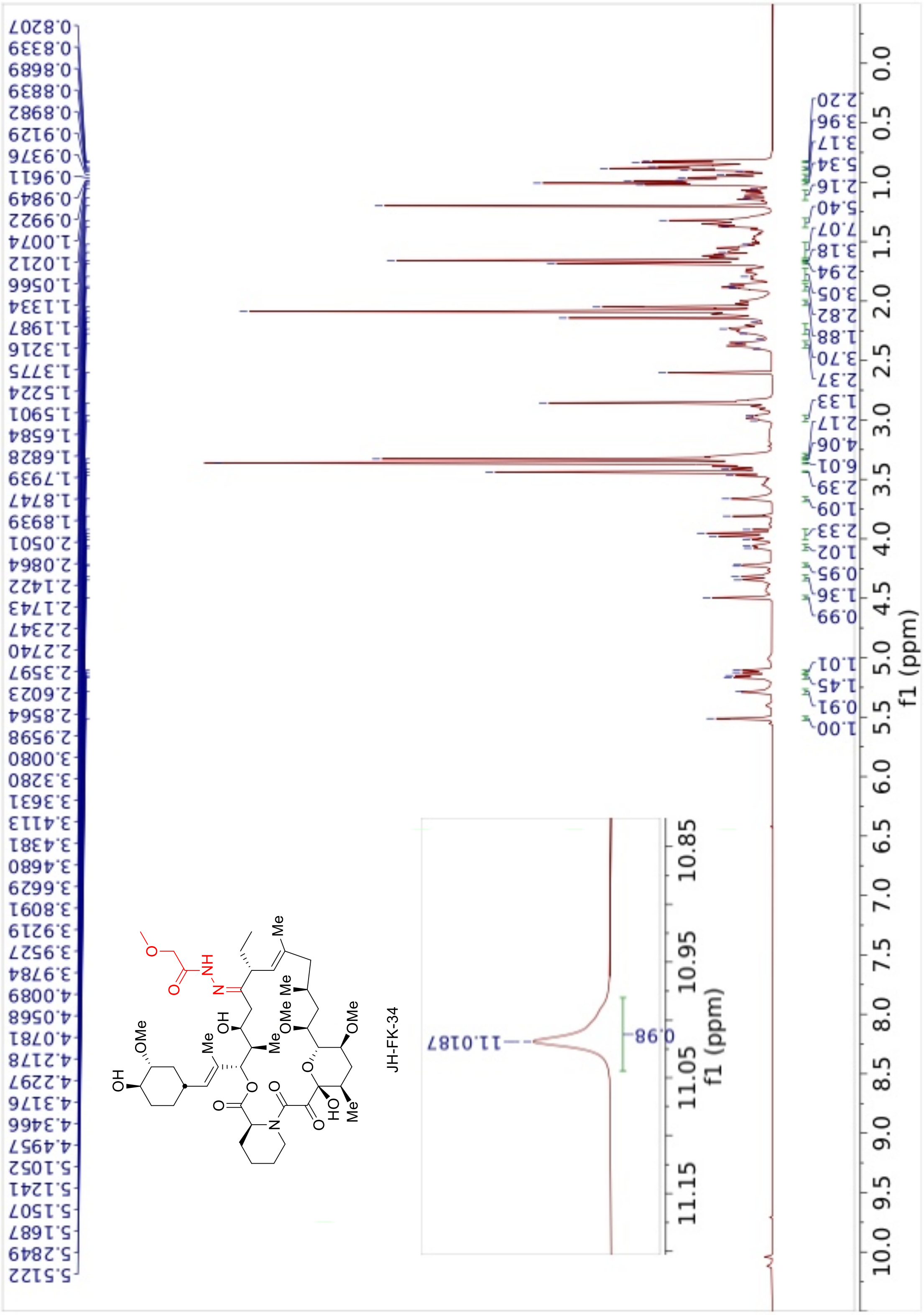
0.8207
0.8339
0.8689
0.8839
0.8982
0.9129
0.9376
0.9611
0.9849
0.9922
1.0074
1.0212
1.0566
1.1334
1.1987
1.3216
1.3775
1.5224
1.5901
1.6584
1.6828
1.7939
1.8747
1.8939
2.0501
2.0864
2.1422
2.1743
2.2347
2.2740
2.3597
2.6023
2.8564
2.9598
3.0080
3.3280
3.3631
3.4113
3.4381
3.4680
3.6629
3.8091
3.9219
3.9527
3.9784
4.0089
4.0568
4.0781
4.2178
4.2297
4.3176
4.3466
4.4957
5.1052
5.1241
5.1507
5.1687
5.2849
5.5122
JH-FK-34
11.0187
0.98
f1 (ppm)
2.20
3.96
3.17
5.34
2.16
5.40
7.07
3.18
2.94
3.05
2.82
1.88
3.70
2.37
1.33
2.17
4.06
6.01
2.39
1.09
2.33
1.02
0.95
1.36
0.99
1.01
1.45
0.91
1.00
f1 (ppm)

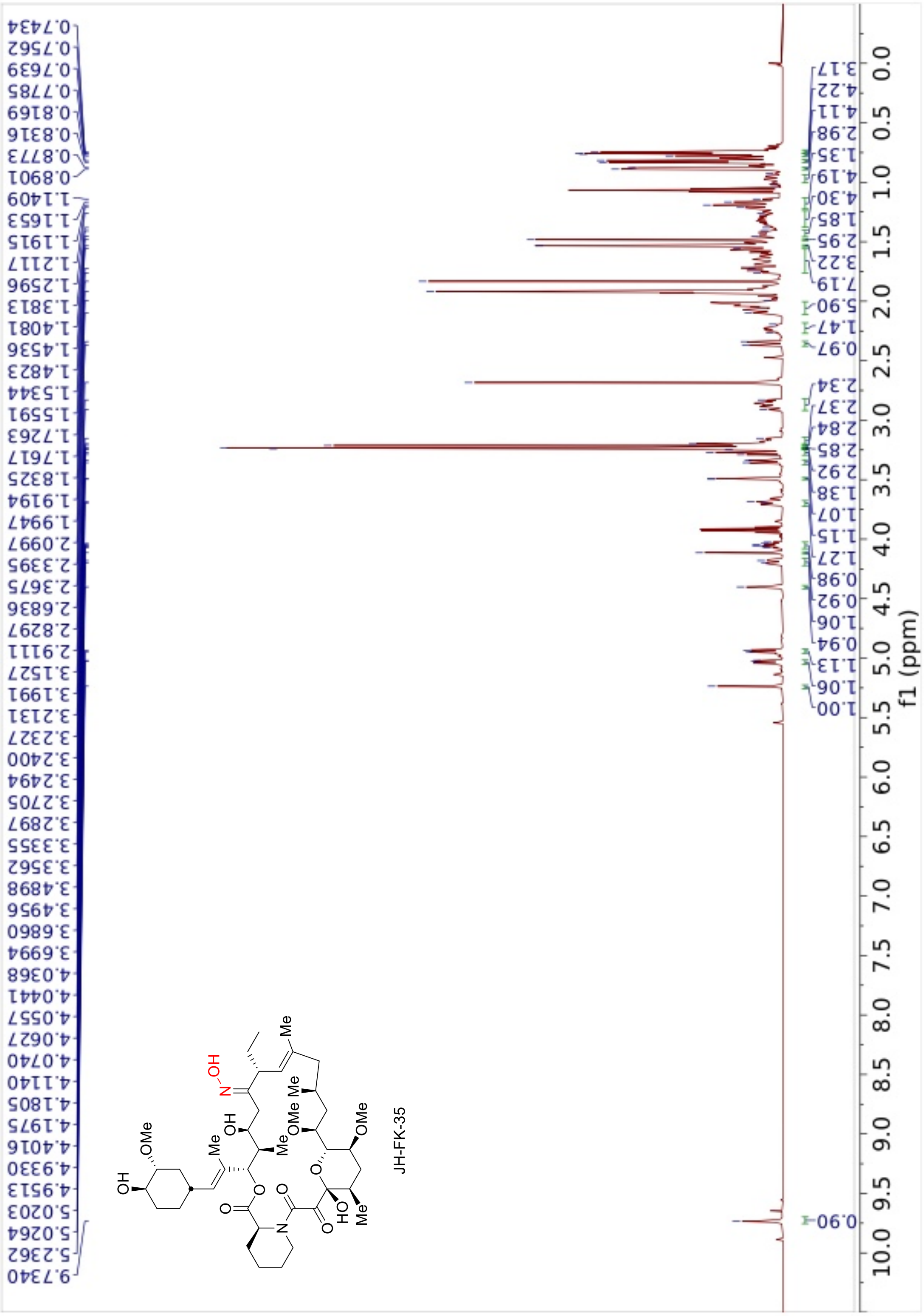
JH-FK-35
f1 (ppm)

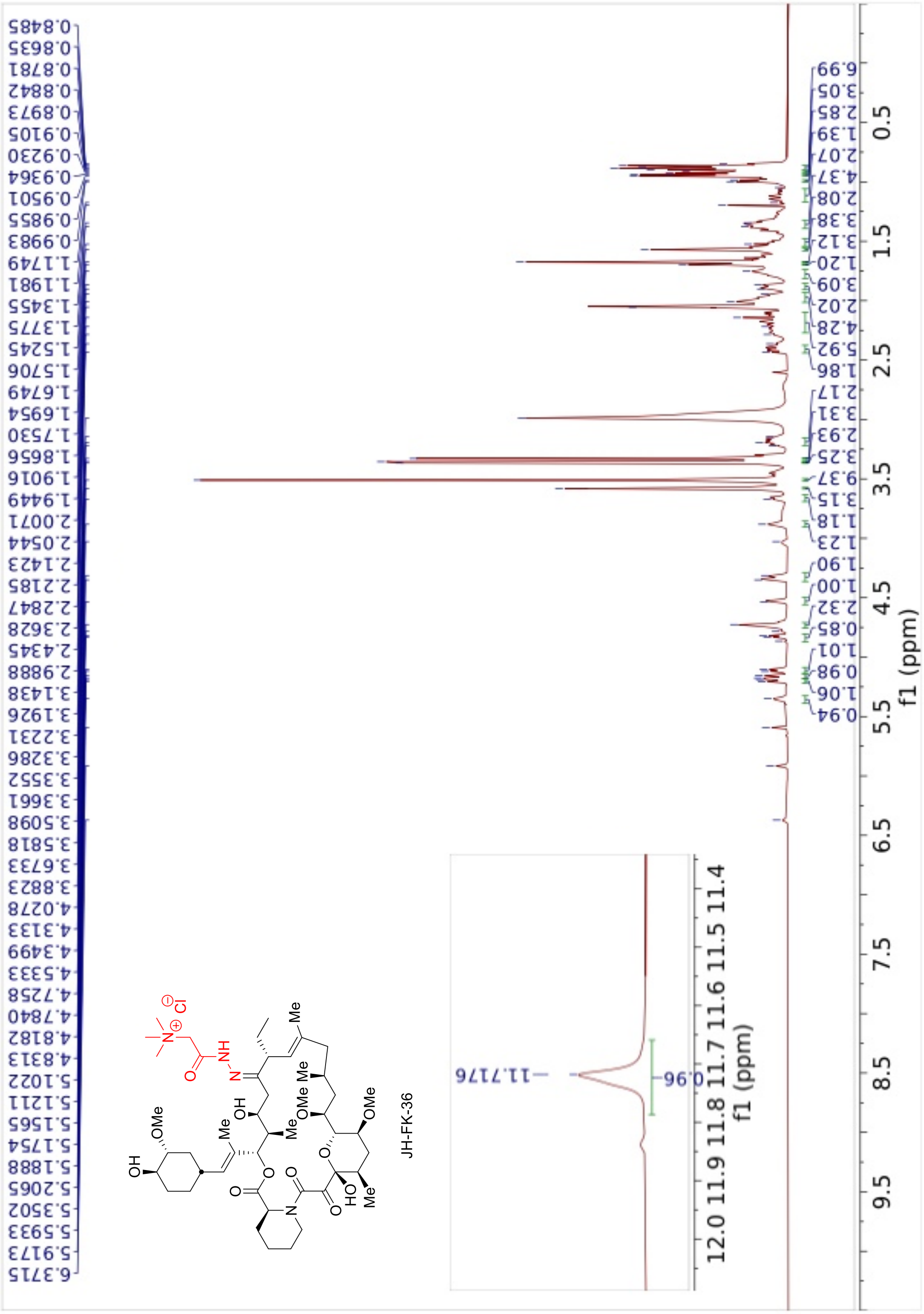
JH-FK-36
f1 (ppm)

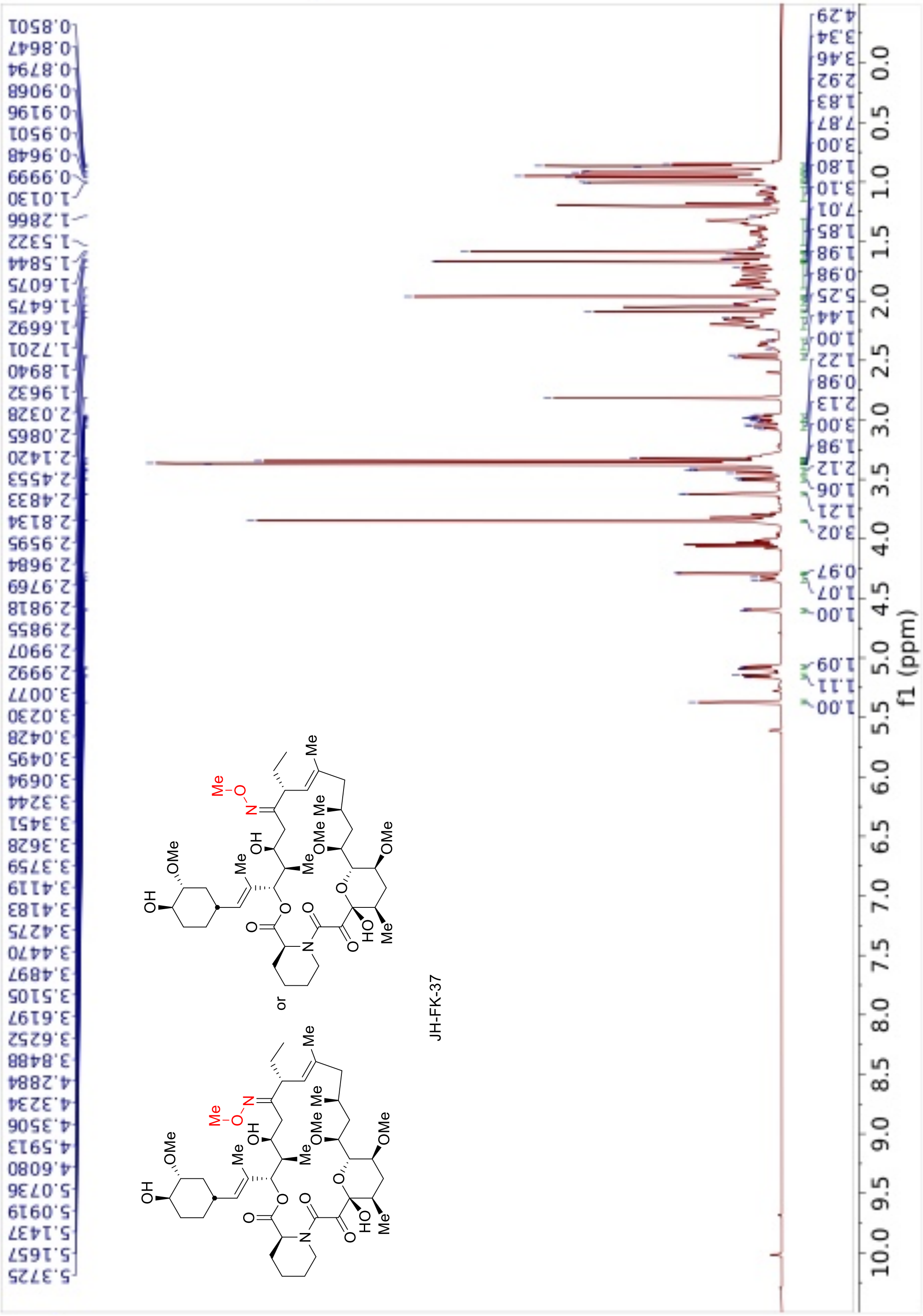

0.8501
0.8647
0.8794
0.9068
0.9196
0.9501
0.9648
0.9999
1.0130
1.2866
1.5322
1.5844
1.6075
1.6475
1.6692
1.7201
1.8940
1.9632
2.0328
2.0865
2.1420
2.4553
2.4833
2.8134
2.9595
2.9684
2.9769
2.9818
2.9855
2.9907
2.9992
3.0077
3.0230
3.0428
3.0495
3.0694
3.3244
3.3451
3.3628
3.3759
3.4119
3.4183
3.4275
3.4470
3.4897
3.5105
3.6197
3.6252
3.8488
4.2884
4.3234
4.3506
4.5913
4.6080
5.0736
5.0919
5.1437
5.1657
5.3725
or
JH-FK-37
f1 (ppm)

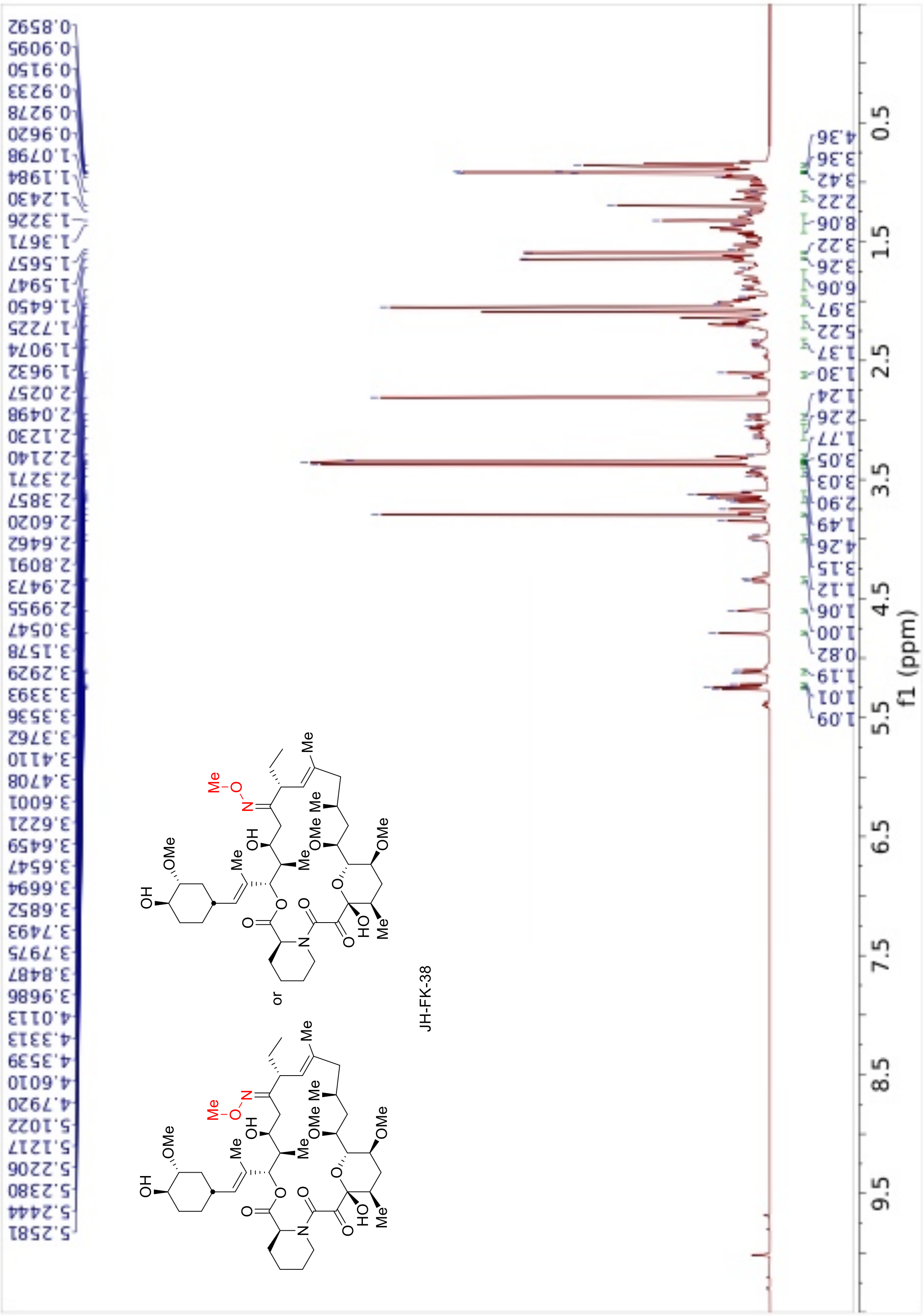
0.8592
0.9095
0.9150
0.9233
0.9278
0.9620
1.0798
1.1984
1.2430
1.3226
1.3671
1.5657
1.5947
1.6450
1.7225
1.9074
1.9632
2.0257
2.0498
2.1230
2.2140
2.3271
2.3857
2.6020
2.6462
2.8091
2.9473
2.9955
3.0547
3.1578
3.2929
3.3393
3.3536
3.3762
3.4110
3.4708
3.6001
3.6221
3.6459
3.6547
3.6694
3.6852
3.7493
3.7975
3.8487
3.9686
4.0113
4.3313
4.3539
4.6010
4.7920
5.1022
5.1217
5.2206
5.2380
5.2444
5.2581
or
JH-FK-38
4.36
3.36
3.42
2.22
8.06
3.22
3.26
6.06
3.97
5.22
1.37
1.30
1.24
2.26
1.77
3.05
3.03
2.90
1.49
4.26
3.15
1.12
1.06
1.00
0.82
1.19
1.01
1.09
f1 (ppm)

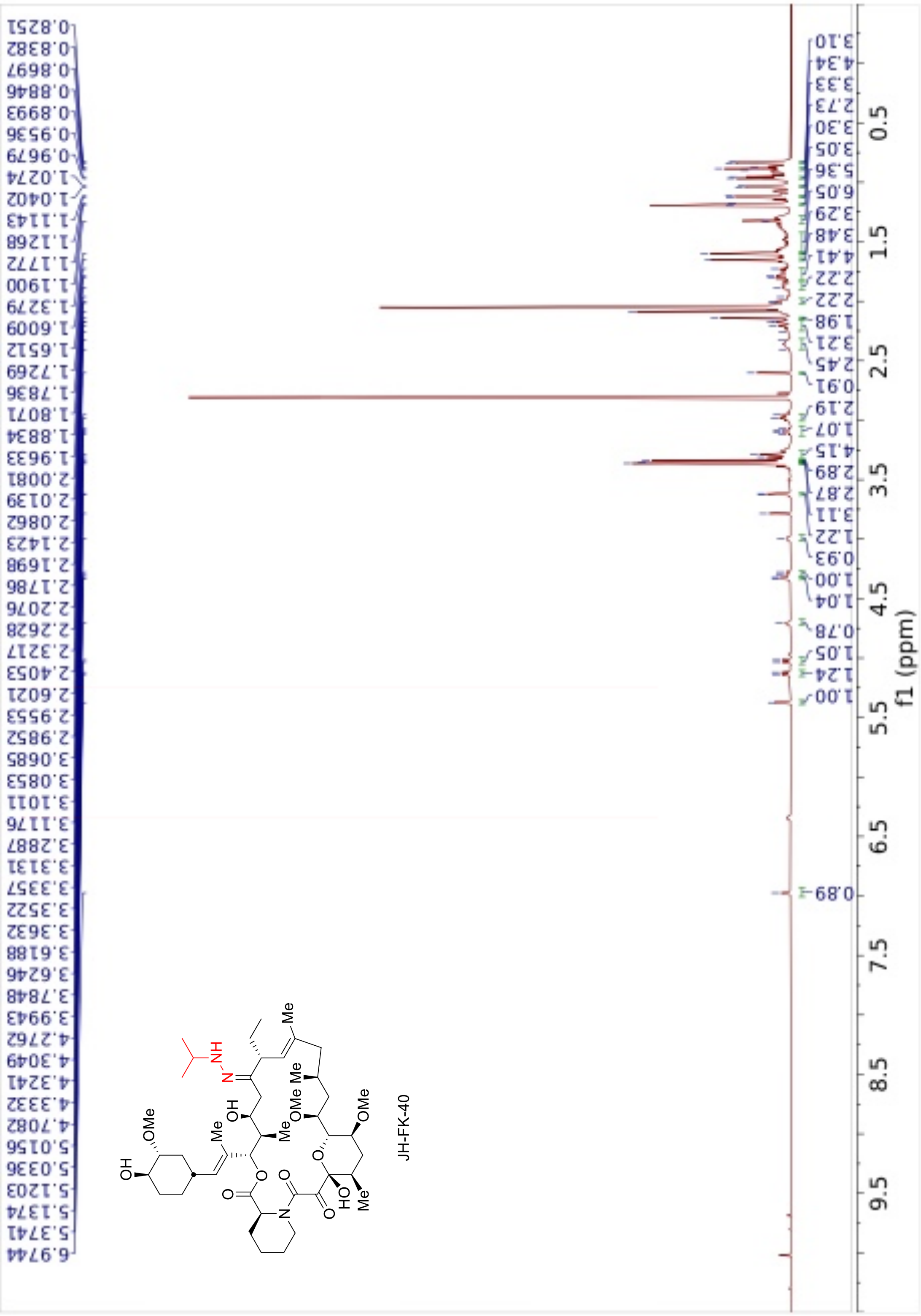

0.8251
0.8382
0.8697
0.8846
0.8993
0.9536
0.9679
1.0274
1.0402
1.1143
1.1268
1.1772
1.1900
1.3279
1.6009
1.6512
1.7269
1.7836
1.8071
1.8834
1.9633
2.0081
2.0139
2.0862
2.1423
2.1698
2.1786
2.2076
2.2628
2.3217
2.4053
2.6021
2.9553
2.9852
3.0685
3.0853
3.1011
3.1176
3.2887
3.3131
3.3357
3.3522
3.3632
3.6188
3.6246
3.7848
3.9943
4.2762
4.3049
4.3241
4.3332
4.7082
5.0156
5.0336
5.1203
5.1374
5.3741
6.9744
JH-FK-40
f1 (ppm)

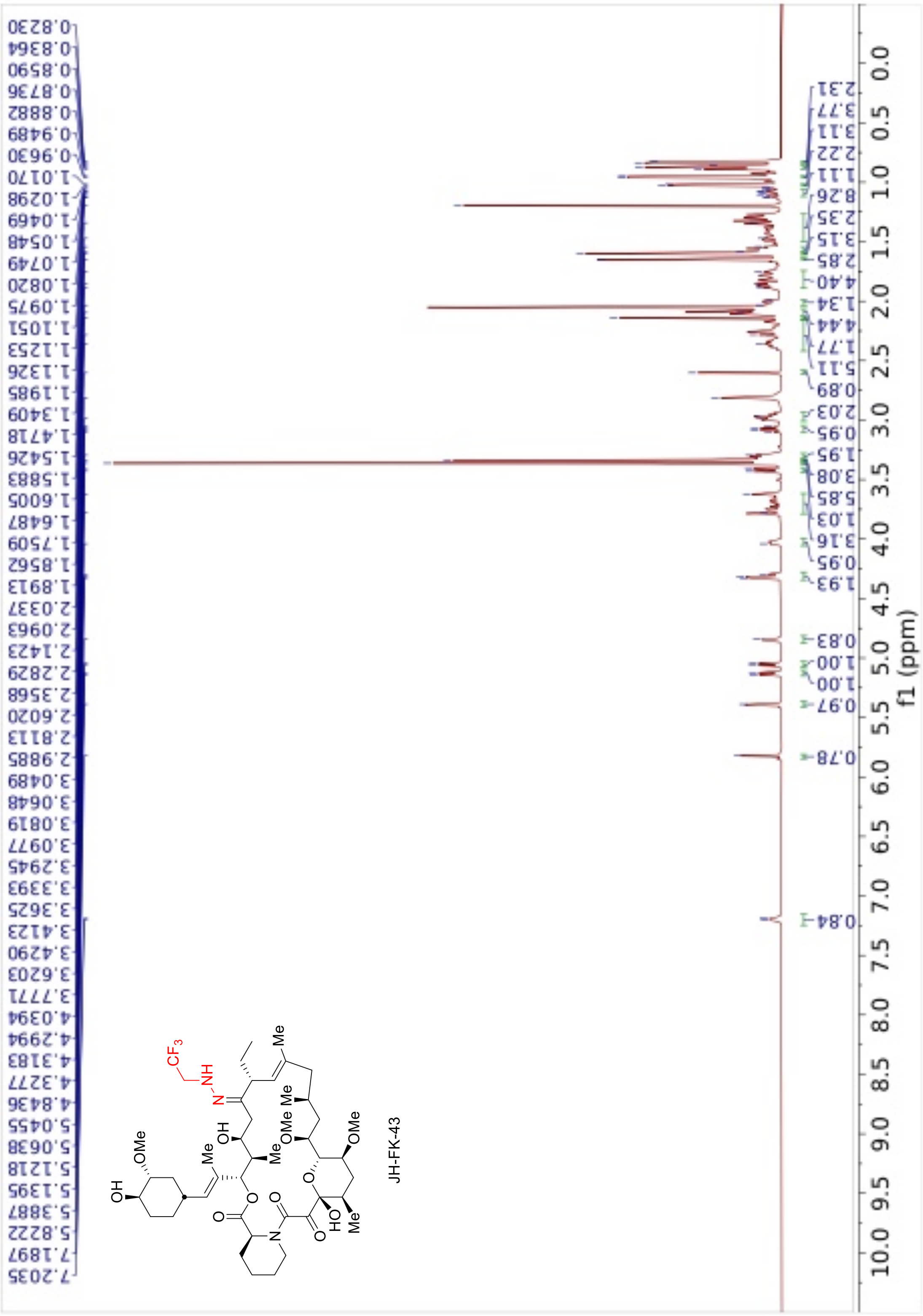
JH-FK-43
f1 (ppm)

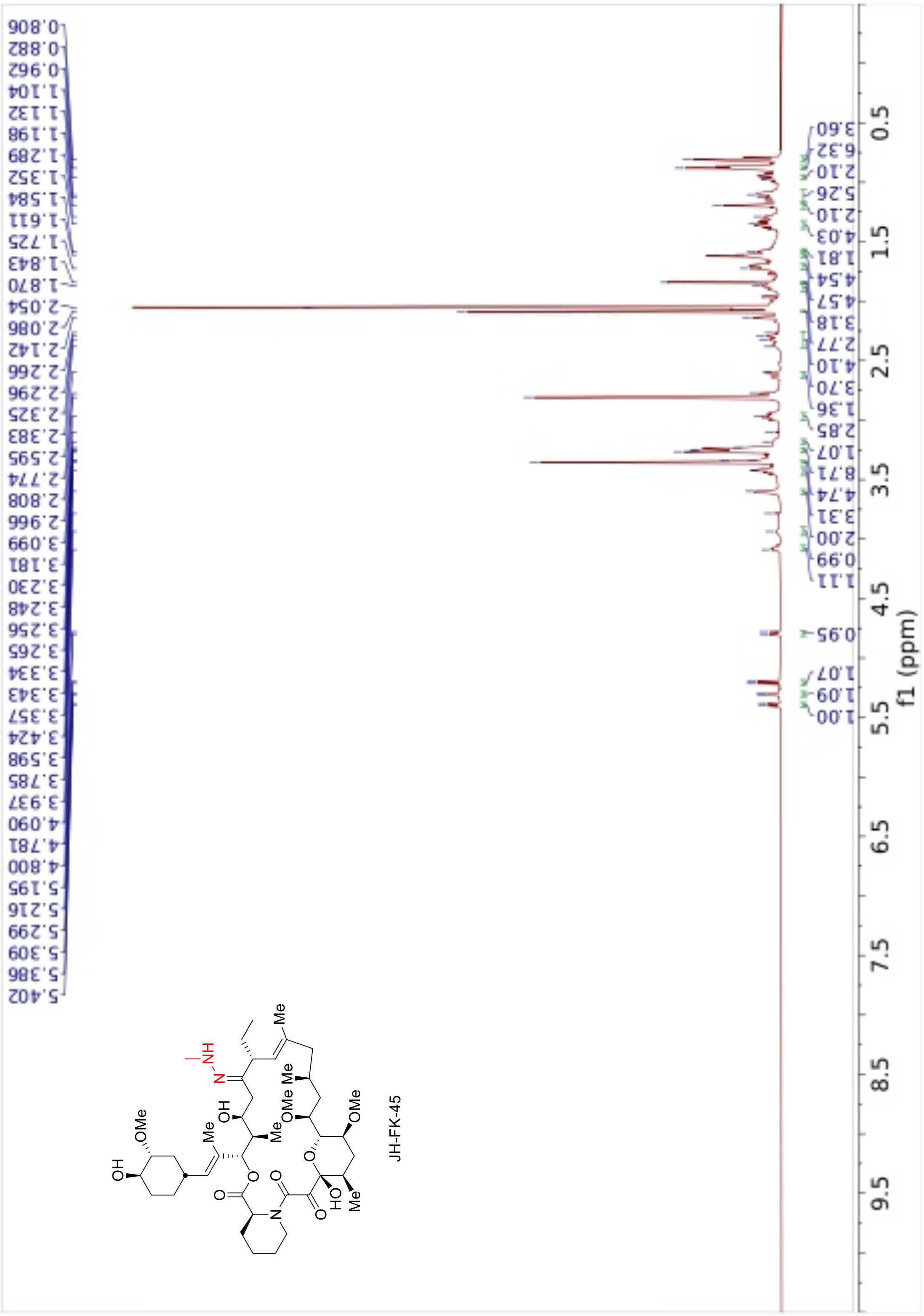
JH-FK-45
f1 (ppm)

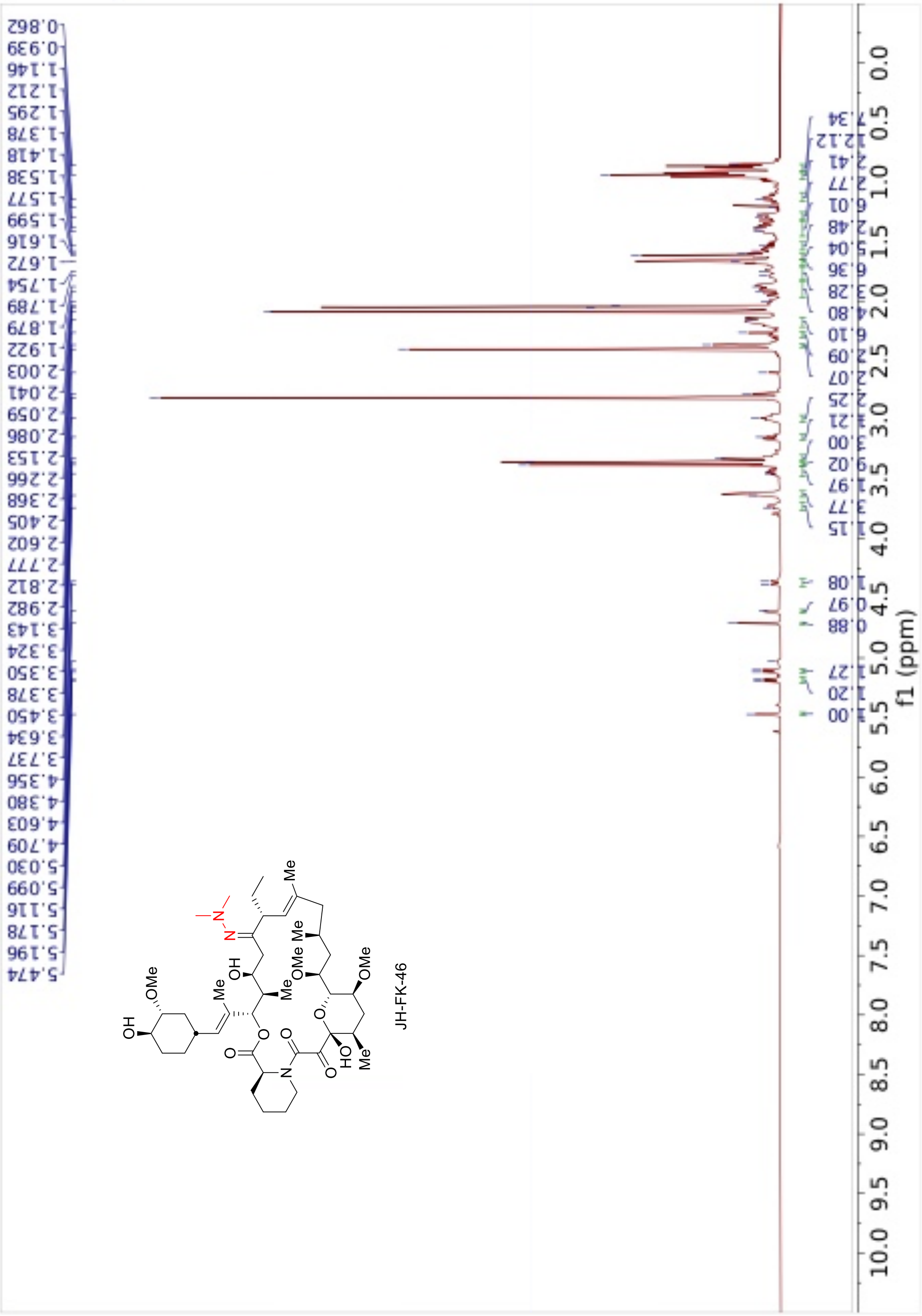

0.862
0.939
1.146
1.212
1.295
1.378
1.418
1.538
1.577
1.599
1.616
1.672
1.754
1.789
1.879
1.922
2.003
2.041
2.059
2.086
2.153
2.266
2.368
2.405
2.602
2.777
2.812
2.982
3.143
3.324
3.350
3.378
3.450
3.634
3.737
4.356
4.380
4.603
4.709
5.030
5.099
5.116
5.178
5.196
5.474
JH-FK-46
f1 (ppm)

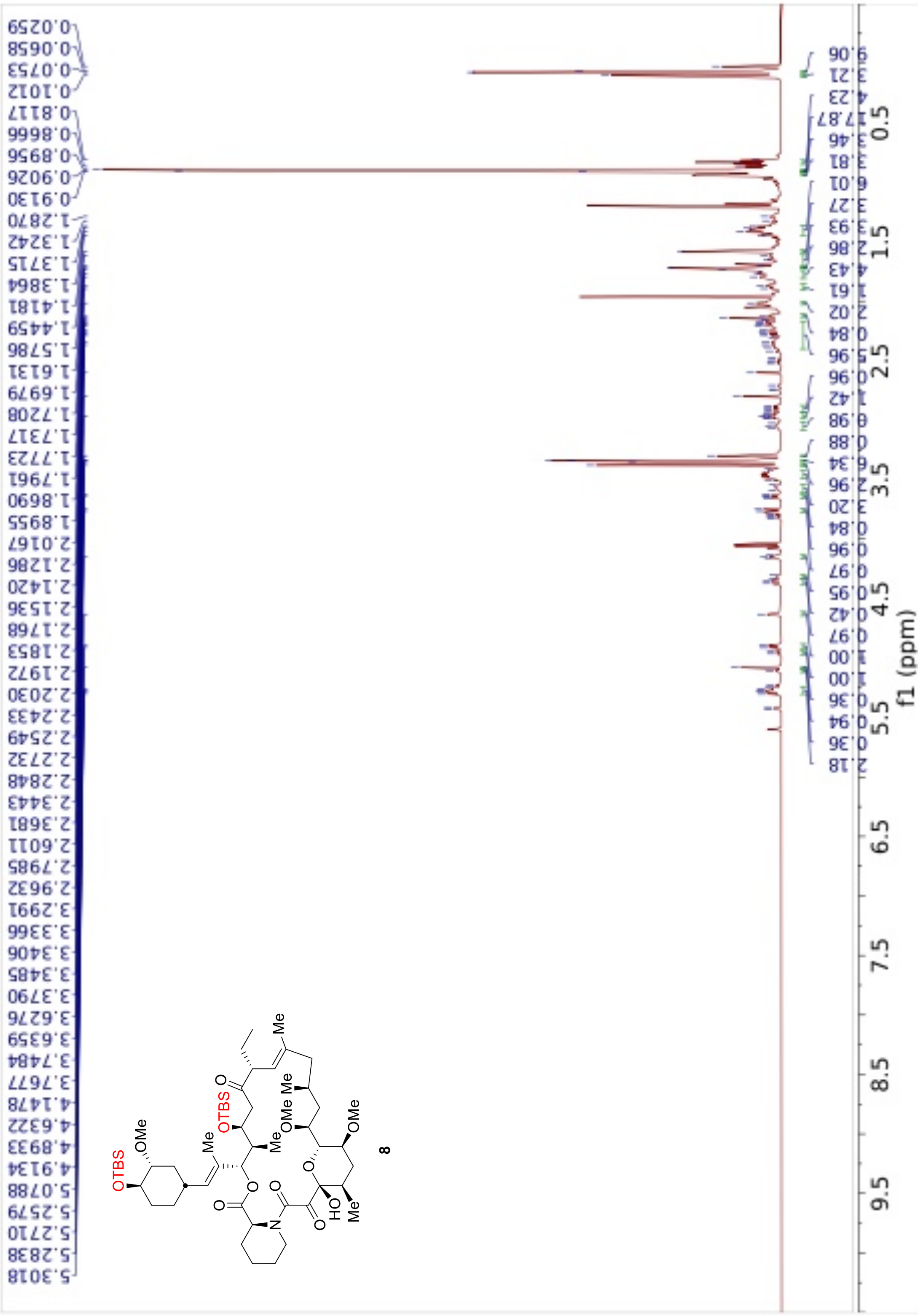
f1 (ppm)
OTBS
OMe
8

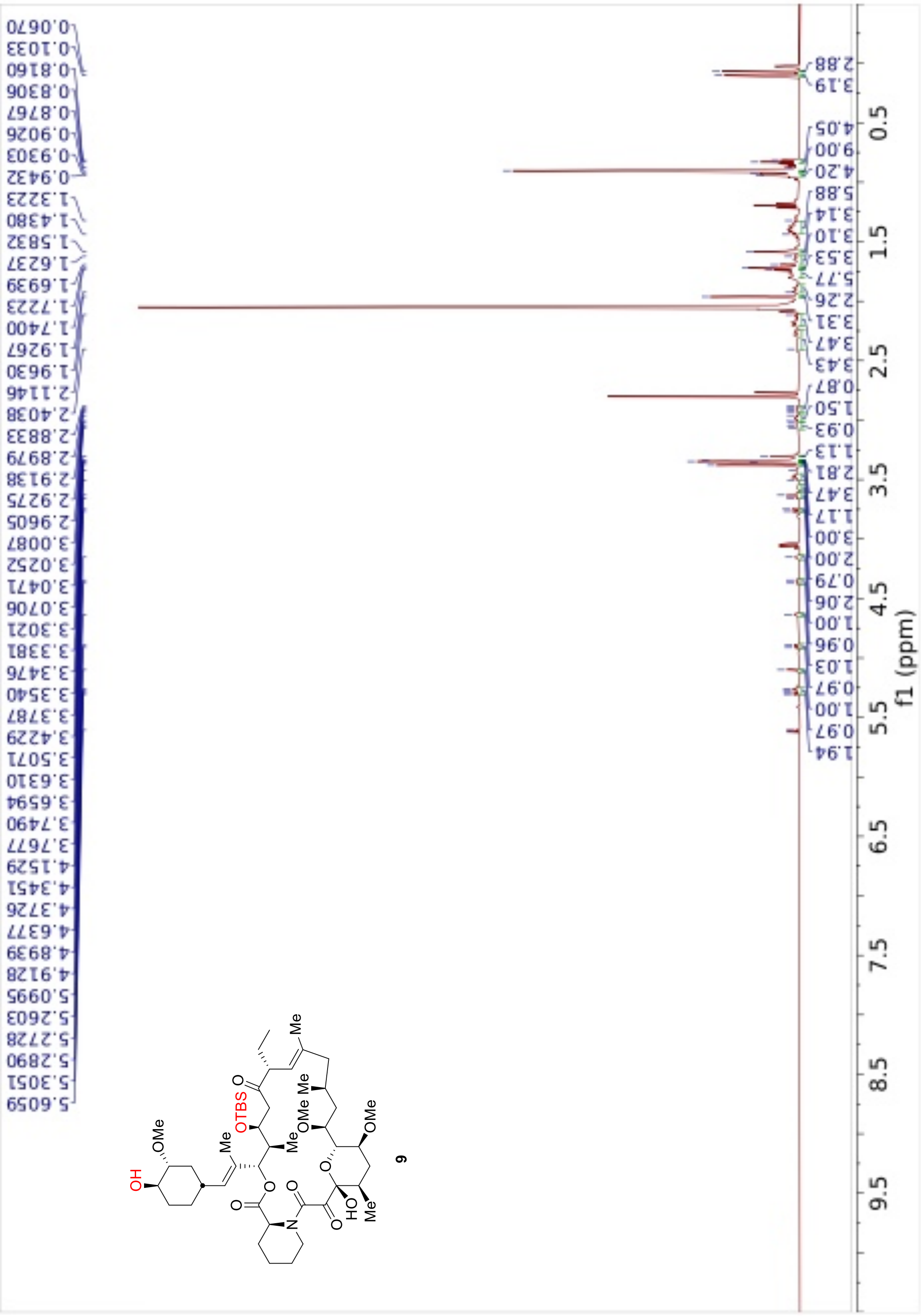

0.0670
0.1033
0.8160
0.8306
0.8767
0.9026
0.9303
0.9432
1.3223
1.4380
1.5832
1.6237
1.6939
1.7223
1.7400
1.9267
1.9630
2.1146
2.4038
2.8833
2.8979
2.9138
2.9275
2.9605
3.0087
3.0252
3.0471
3.0706
3.3021
3.3381
3.3476
3.3540
3.3787
3.4229
3.5071
3.6310
3.6594
3.7490
3.7677
4.1529
4.3451
4.3726
4.6377
4.8939
4.9128
5.0995
5.2603
5.2728
5.2890
5.3051
5.6059
2.88
3.19
4.05
9.00
4.20
5.88
3.14
3.10
3.53
5.77
2.26
3.31
3.47
3.43
0.87
1.50
0.93
1.13
2.81
3.47
1.17
3.00
2.00
0.79
2.06
1.00
0.96
1.03
0.97
1.00
0.97
1.94
0.5
1.5
2.5
3.5
4.5
5.5
6.5
7.5
8.5
9.5
f1 (ppm)
OH
OMe
Me
OTBS
O
Me
OMe
Me
OMe
N
HO
Me
9

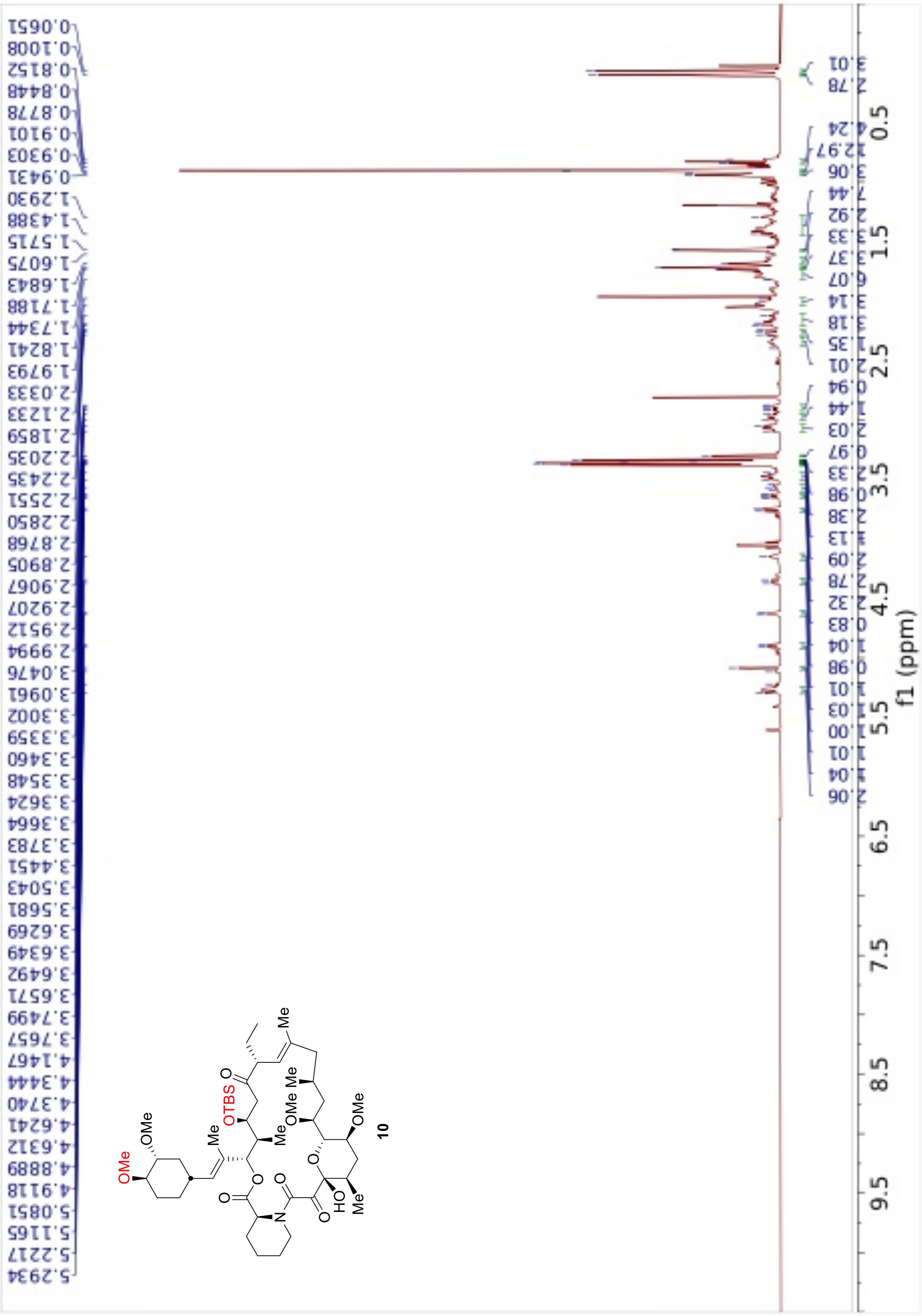
f1 (ppm)
OTBS
OMe
10

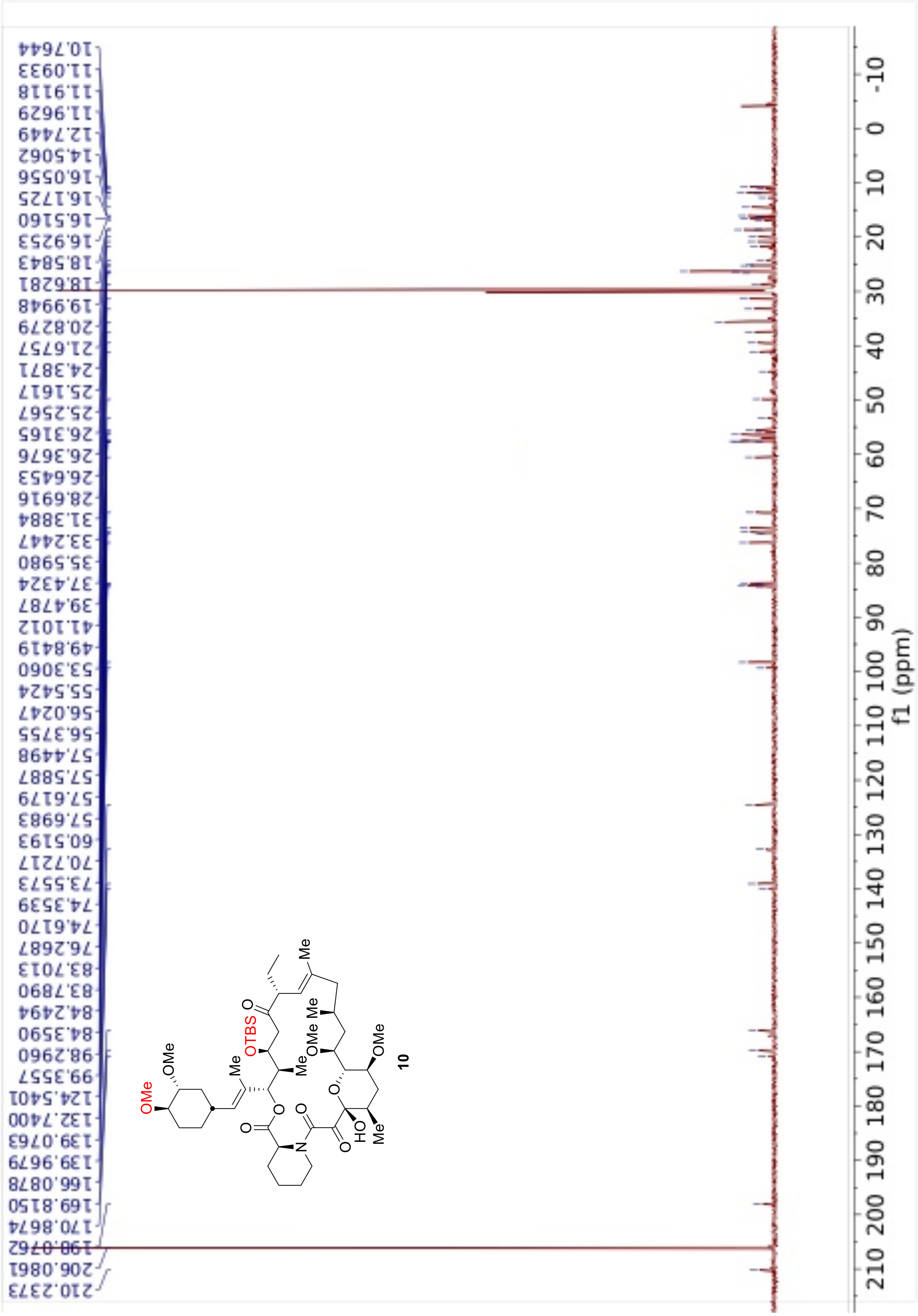
10.7644
11.0933
11.9118
11.9629
12.7449
14.5062
16.0556
16.1725
16.5160
16.9253
18.5843
18.6281
19.9948
20.8279
21.6757
24.3871
25.1617
25.2567
26.3165
26.3676
26.6453
28.6916
31.3884
33.2447
35.5980
37.4324
39.4787
41.1012
49.8419
53.3060
55.5424
56.0247
56.3755
57.4498
57.5887
57.6179
57.6983
60.5193
70.7217
73.5573
74.3539
74.6170
76.2687
83.7013
83.7890
84.2494
84.3590
98.2960
99.3557
124.5401
132.7400
139.0763
139.9679
166.0878
169.8150
170.8674
198.0762
206.0861
210.2373
f1 (ppm)
OMe
OMe
Me
OTBS
Me
OMe
Me
OMe
Me
HO
Me
N
10

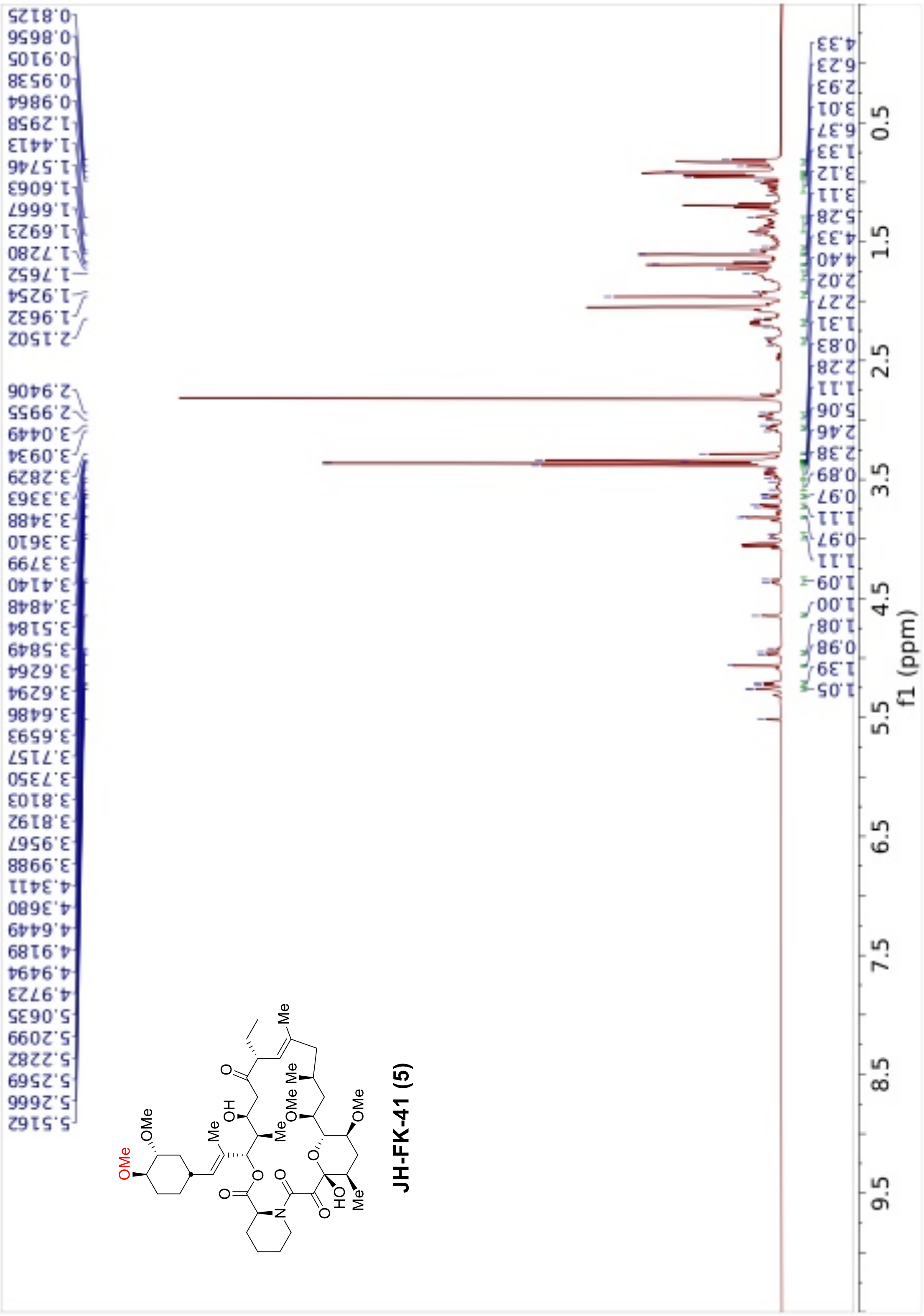

JH-FK-41 (5)
f1 (ppm)

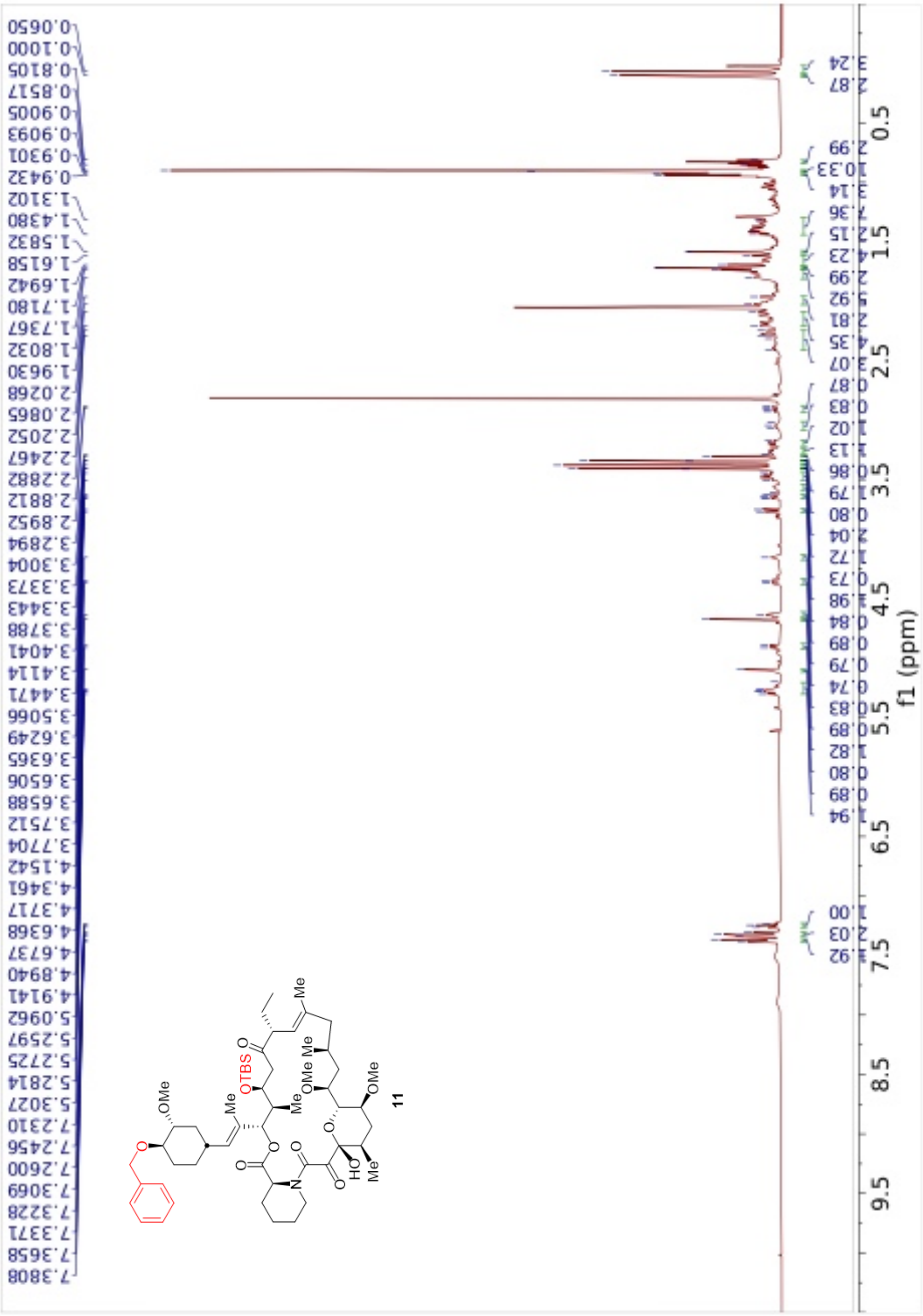

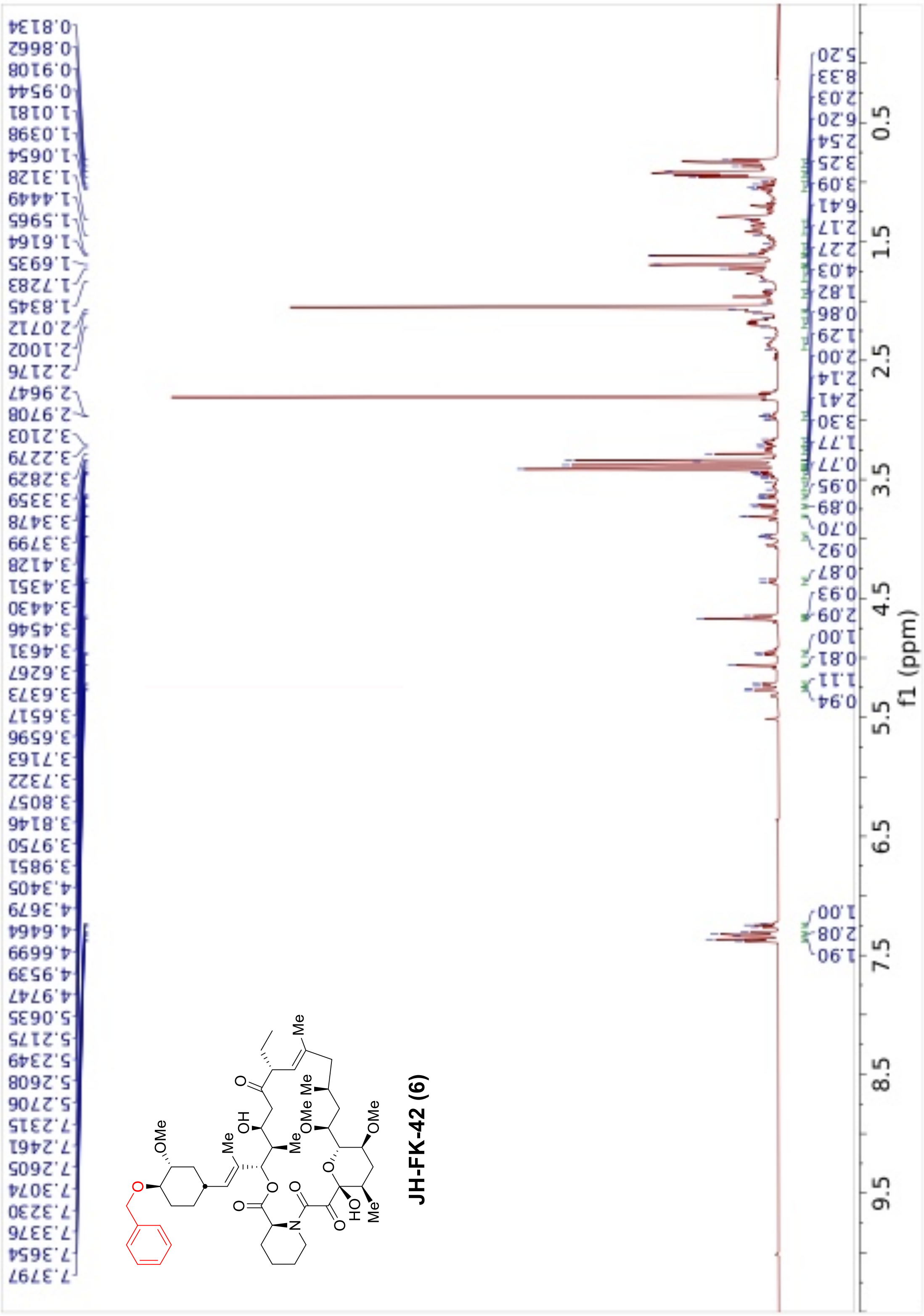
0.8134
0.8662
0.9108
0.9544
1.0181
1.0398
1.0654
1.3128
1.4449
1.5965
1.6164
1.6935
1.7283
1.8345
2.0712
2.1002
2.2176
2.9647
2.9708
3.2103
3.2279
3.2829
3.3359
3.3478
3.3799
3.4128
3.4351
3.4430
3.4546
3.4631
3.6267
3.6373
3.6517
3.6596
3.7163
3.7322
3.8057
3.8146
3.9750
3.9851
4.3405
4.3679
4.6464
4.6699
4.9539
4.9747
5.0635
5.2175
5.2349
5.2608
5.2706
7.2315
7.2461
7.2605
7.3074
7.3230
7.3376
7.3654
7.3797
JH-FK-42 (6)
f1 (ppm)

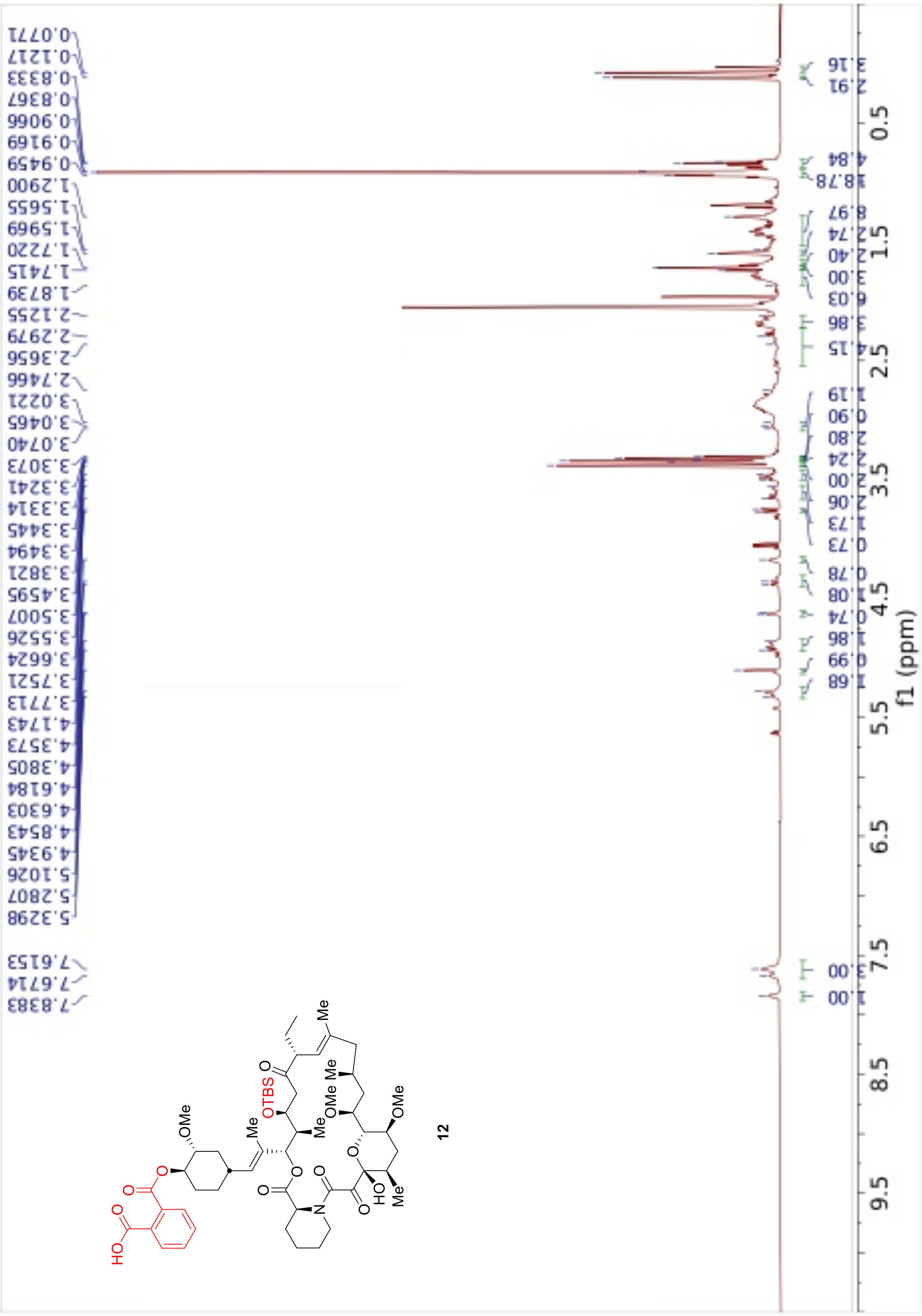
12
f1 (ppm)

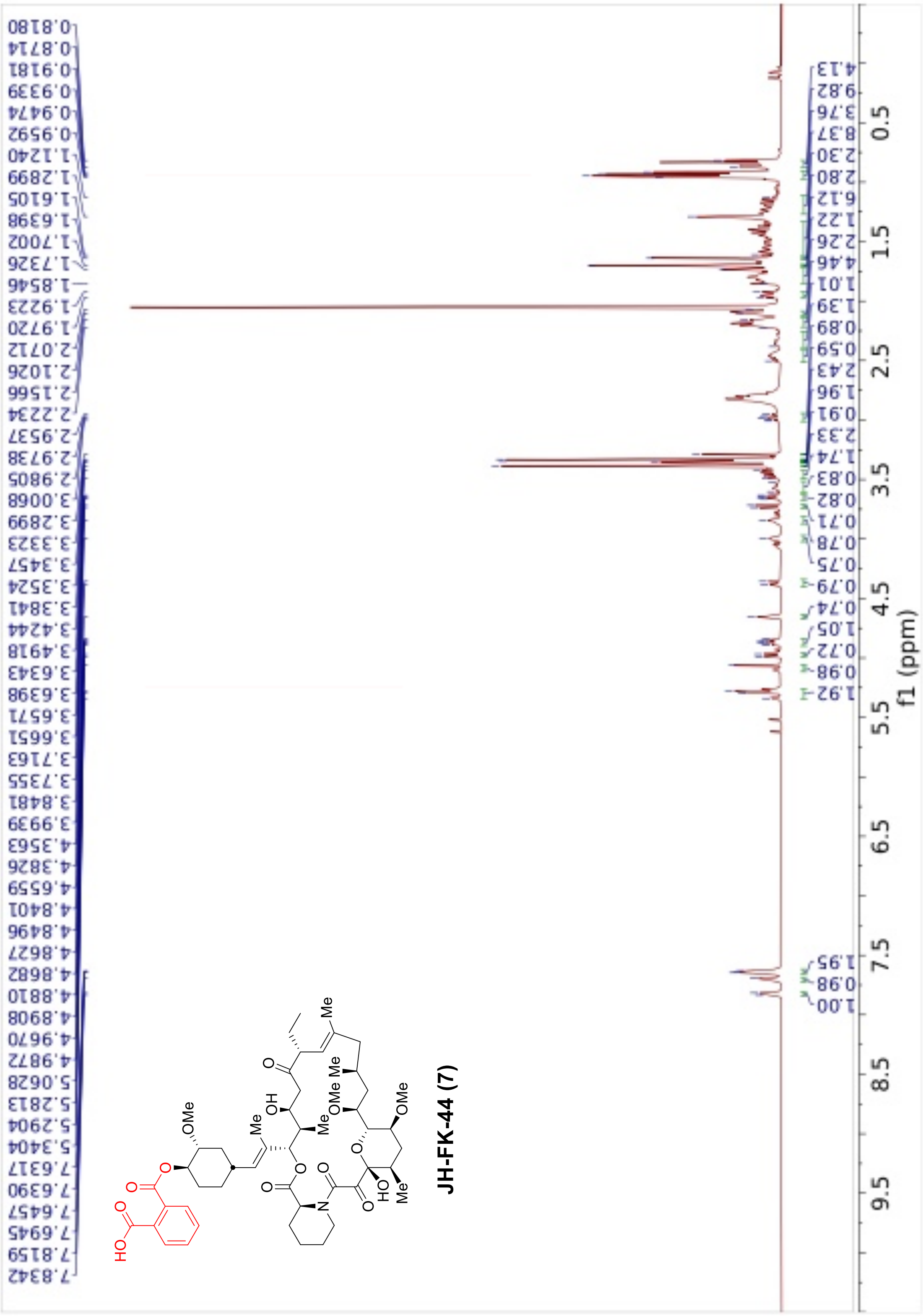

7.8342
7.8159
7.6945
7.6457
7.6390
7.6317
5.3404
5.2904
5.2813
5.0628
4.9872
4.9670
4.8908
4.8810
4.8682
4.8627
4.8496
4.8401
4.6559
4.3826
4.3563
3.9939
3.8481
3.7355
3.7163
3.6651
3.6571
3.6398
3.6343
3.4918
3.4244
3.3841
3.3524
3.3457
3.3323
3.2899
3.0068
2.9805
2.9738
2.9537
2.2234
2.1566
2.1026
2.0712
1.9720
1.9223
1.8546
1.7326
1.7002
1.6398
1.6105
1.2899
1.1240
0.9592
0.9474
0.9339
0.9181
0.8714
0.8180
f1 (ppm)
OMe
HO
Me
OH
JH-FK-44 (7)
